# pH-induced structural changes in SARS-CoV-2 spike variants

**DOI:** 10.1101/2025.11.17.688702

**Authors:** A. Sofia F. Oliveira, Lorenzo Tulli, Fiona L. Kearns, Lorenzo Casalino, Mia A. Rosenfeld, Imre Berger, Christiane Schaffitzel, Andrew D. Davidson, Allen E. Haddrell, Jonathan P. Reid, Rommie E. Amaro, Adrian J. Mulholland

## Abstract

pH critically influences SARS-CoV-2 infectivity and stability by, for example, triggering pH-dependent conformational changes in the spike glycoprotein that can facilitate membrane fusion and viral entry into host cells. Using the emerging dynamical nonequilibrium molecular dynamics (D-NEMD) simulations approach, we investigated how biologically relevant pH shifts affect the functional dynamics of the fully glycosylated spike from the ancestral, Delta, and Omicron BA.1 variants of concern. For this, over 1100 nonequilibrium simulations were conducted to capture the pH-induced structural and dynamic changes that occur following transitions from physiological to acidic and alkaline conditions, with the former mimicking the low pH environment within endosomes, and the latter the high pH conditions accessible to nascent exhaled aerosols. D-NEMD reveals that pH changes trigger distinct, variant-specific conformational responses in key regions of the spike, including the receptor binding domain (RBD), fusion peptide proximal region (FPPR), and C-terminal domain (CTD). The ancestral spike shows broad pH sensitivity, characterised by directional motions and region-specific structural rearrangements that depend on the pH conditions. The spike of the Delta variant displays increased reactivity to alkaline pH, potentially explaining its reduced stability in alkaline aerosols. The spike of Omicron BA.1, in contrast, responds strongly to acidic conditions with spontaneous RBD opening and pronounced structural rearrangements in the FPPR and CTD. This behaviour aligns with this variant’s preference for endosomal entry and reduced reliance on TMPRSS2-mediated fusion. The Omicron BA.1 spike also shows increased resilience to alkaline pH, suggesting greater environmental stability. Our findings further emphasise the key role of glycans in spike activation, with glycan N234 stabilising the RBD “up” conformation during pH-induced transitions in Omicron under acidic conditions. These insights, together, highlight pH as a potential evolutionary pressure for SARS-CoV-2 and underscore the importance of glycosylation and environmental pH variability in shaping the behaviour of viral fusion proteins.

## Introduction

pH plays a crucial role in biology as it can influence the structure, dynamics, function, stability and solubility of proteins (1–3). Intracellular pH (and its fluctuations) is known to play a broad regulatory role in a range of (normal and pathological) essential cellular processes, such as enzyme activity, membrane transport, cell growth and division and apoptosis (4). Eukaryotic cells, in particular, are highly compartmentalised, with the individual subcellular compartments providing a distinct pH environment tailored to their specific function. For example, the cytosol and nucleus typically maintain a near-neutral pH (pH ∼ 7), while mitochondria are slightly alkaline (pH ∼ 8), and lysosomes and endosomes are markedly acidic, with pH values around 5 (5). These differences in cellular or subcellular pH can, in some cases, be exploited for physiological purposes. For example, many enveloped viruses use the acidic conditions within sub-cellular endosomal compartments to facilitate host cell invasion (6, 7). These viruses possess envelope proteins that are sensitive to pH fluctuations, such as the spike in coronaviruses and hemagglutinin in influenza, which trigger the infection process once the appropriate conditions are met (6, 7).

In SARS-CoV-2, the spike protein is the primary focus of the virus’ molecular response to pH, with its pH-dependent conformational changes initiating the host cell infection process (6). Viral entry can proceed *via* two pathways: one involving direct fusion of the virus at the plasma membrane; and the other occurring through receptor-mediated endocytosis within endosomal compartments (8, 9). The latter pathway is particularly pH-dependent, as the acidic microenvironment within endosomes triggers dramatic conformational changes in the spike, leading to the exposure of the fusion machinery and ultimately facilitating the fusion of the viral envelope with the host endosomal membrane (6, 7). This fusion allows for the release of the viral genome into the host cell cytoplasm, thus initiating infection (6, 7).

Beyond their role in infection, pH changes also significantly influence the aerostability of SARS-CoV-2, which primarily spreads through the inhalation of respiratory droplets and aerosol particles exhaled by infected individuals, as well as *via* contaminated surfaces (10–12). Large respiratory droplets (i.e. >100 μm in diameter), produced by coughing or sneezing, can transmit pathogens over only short distances (<0.5 m). However, aerosol particles <100 μm in diameter, generated through speaking and exhalation, are sufficiently small to remain airborne for long periods of time, travel significant distances, accumulate in poorly ventilated spaces, and can be inhaled (10–12). For instance, fine aerosols loaded with SARS-CoV-2 particles have been shown to remain in the air up to several hours, depending on environmental conditions such as relative humidity, carbon dioxide concentration and temperature (13–18). Once inhaled, aerosol particles can deposit in various parts of the respiratory tract and initiate the infection process, with the larger particles typically depositing in the upper airway and smaller ones penetrating deeper into the lungs (10–12). Experimental studies have demonstrated that these airborne particles can undergo physical and chemical changes during transport due to their equilibration with the surrounding environment, including changes in water content and shifts in their pH (e.g. (16–23)). These pH alterations can significantly impact the stability of viruses while airborne, including the structure and dynamics of their proteins, thereby influencing their infectivity (e.g. (16–18, 22, 24)). However, a unified understanding of how pH evolves during aerosol transport is still debated (23), with some studies reporting that these particles become alkaline during transport (16–18) and others suggesting they acidify instead (22).

As previously noted, the spike, which is the central focus of the current study, plays a pivotal role in SARS-CoV-2 response to pH changes. Structurally, it is a homotrimeric glycoprotein (Figure 1A) that facilitates viral entry by binding to various human receptors, most notably angiotensin-converting enzyme 2 (ACE2) (25, 26) but also neuropilin-1 (27, 28), estrogen receptor α (29), glucose-regulated protein 78 (30), transferrin receptor (31), cluster of differentiation 147 (32), and (potentially) to sugar (33) and nicotinic acetylcholine (34–36) receptors. The full-length ancestral spike (also often referred to as wild-type, ‘early 2020’, ‘Wuhan’, or original protein) comprises 1273 amino-acid residues (37–39). Each spike monomer features three primary topological domains: the head, stalk, and cytoplasmic tail (37–39). The soluble extracellular head domain (Figure 1) contains all components required for receptor binding and membrane fusion (37–39) and is divided into two main functional regions: the receptor-binding fragment S1 and the fusion fragment S2 (37–39). It is formed by the N-terminal domains (NTDs), which contribute to immune evasion; the receptor-binding domains (RBDs), which directly engage with the host receptors; the fusion peptides (FPs), that facilitate membrane fusion; two proteolytic cleavage sites that modulate protein infectivity and stability (40–44); and, various regulatory allosteric sites, including a fatty acid binding site (45) and a heme/biliverdin site (46–48). Experiments have shown that the stability and function of the spike are highly sensitive to pH, affecting, for example, the conformation of the RBD and its interactions with host receptors (e.g. (9, 49, 50)). This modulation arises from changes in the protonation state of the tens of titratable groups in the protein (Figure 1C), which alter the protein’s charge distribution (e.g. (51–53)) and affect its structure and dynamics and its ability to form intermolecular interactions with host receptors.

**Figure 1.**
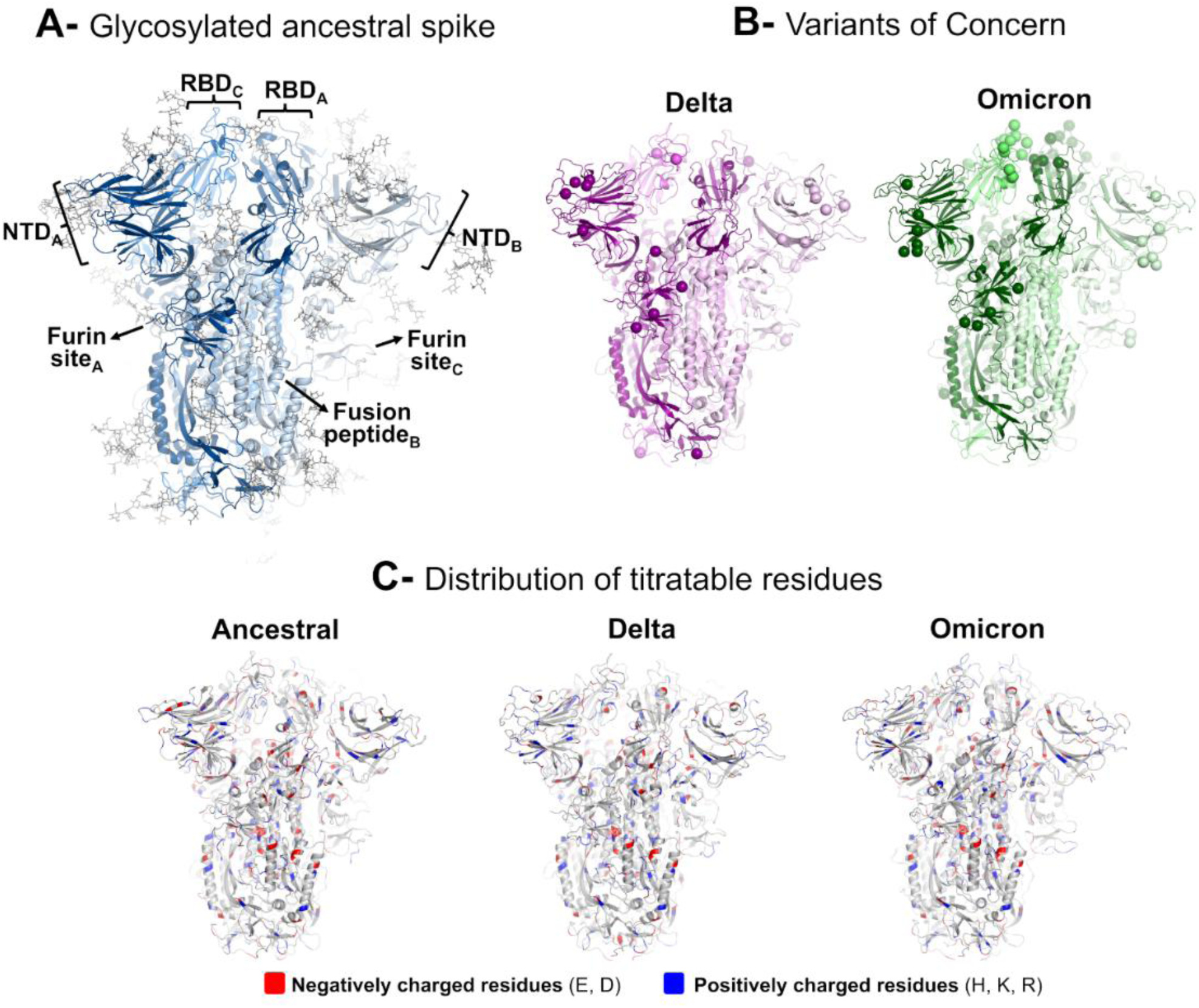
Structure of the glycosylated ectodomain of the SARS-CoV-2 spike. (**A**) Head region of the glycosylated ancestral spike. This structure, constructed from PDB IDs 7JJI (54) and 6VXX (37), corresponds to the closed conformation of the spike (in which all RBDs are in the “down” conformation, buried inside the trimer, therefore unable to bind to host ACE2 receptors) (55, 56). The spike is formed by three monomers, each one shown in a different shade of blue: monomers A, B and C are coloured in dark, light and marine blue, respectively. The RBDs and NTDs, furin sites and fusion peptides are subscripted with their chain ID (A, B or C). Glycans are represented with grey sticks. (**B**) Models for the spike head region of the Delta and Omicron BA.1 variants in the closed state. The monomers in the Delta and Omicron spikes are highlighted in shades of pink and green, respectively. The spheres show the positions of the substitutions, deletions, and insertions in the two variants of concern. Delta contains eight substitutions and two deletions relative to the ancestral protein, namely T19R, G142D, E156Δ-F157Δ, R158G, L452R, T478K, D614G, P681R, and D950N (57–60). Omicron BA.1 harbours up to forty changes relative to the ancestral spike, namely A67V, H69Δ-V70Δ, T95I, G142D, V143Δ-Y145Δ, N211Δ, L212I, EPE insertion in position 214, G339D, S371L, S373P, S375F, K417N, N440K, G446S, S477N, T478K, E484A, Q493R, G496S, Q498R, N501Y, Y505H, T547K, D614G, H655Y, N679K, P681H, N764K, D796Y, N856K, Q954H, N969K, and L981F (61–64). (**C**) Location of residues with negatively (coloured in red) and positively (coloured in blue) charged side-chains in the ancestral, Delta and Omicron BA.1 spikes. Depending on the pH conditions, aspartic (D) and glutamic (E) acids can carry negatively charged side-chains, whereas histidine (H), lysine (K), and arginine (R) can possess positively charged ones. Glycans are omitted from panels B and C for clarity, but were included in all MD simulations performed here.

Biomolecular modelling and simulation studies have provided unprecedented molecular-level insights into how pH influences the spike’s mechanism of action, including revealing aspects such as its pH-dependent molecular interactions and pH-regulated functional dynamics (e.g. (51–53, 65–76)). These studies employed various modelling approaches, ranging from continuum electrostatics calculations to equilibrium molecular dynamics (MD) and constant-pH MD simulations (51–53, 65–74, 76). For instance, Warwicker and co-workers conducted p*K*_a_ calculations across a broad set of experimentally determined spike structures to pinpoint residues involved in pH sensing (66–68). Their analysis identified several acidic residues, such as D290, D389 and D614, that are predicted to contribute to the protein’s stability under mildly acidic conditions and influence the open/closed RBD equilibrium (66–68). D614, in particular, was among the first residues to mutate in the ancestral spike protein, giving rise to the D614G variant, with this substitution having a significant impact on trimer stability (77). Equilibrium MD simulations have also been employed to explore the dynamics of the non-glycosylated spike across a range of pH conditions, showing that the protein remains structurally stable between pH 5 and 9 and undergoes pronounced changes at extreme pH conditions (pH=1, 3 and 11) (72, 74). In addition, constant-pH simulations were also conducted on the non-glycosylated ancestral spike to gain new insights into the closed-to-open transition of the RBD, with these simulations identifying D571 as a potential pH switch regulating this transition (69).

Despite significant progress, most studies examining the pH-dependent behaviour of the spike have focused on its non-glycosylated form, omitting the *N-* and *O-*linked glycans (37, 78–83). It is now well-established that glycans are critical for spike function and protection, including modulating its functional dynamics and participating in closed-to-open RBD transitions (55, 56, 75, 84–86). In our previous work, we extended the dynamical-nonequilibrium MD (D-NEMD) simulations approach to examine the response of the fully glycosylated spike from the Delta variant of concern to a pH decrease (75). These simulations identified noticeable changes in the conformational dynamics of functionally important regions, including the RBD (75). Here, we leverage the D-NEMD approach to explore how biologically relevant pH shifts affect the functional dynamics of the fully glycosylated ancestral spike, as well as those from two variants of concern, namely Delta and Omicron BA.1 (Figure 1B). D-NEMD has emerged as a valuable computational tool for probing biomolecular systems (87, 88), for example, identifying the propagation of structural rearrangements, mapping allosteric communication networks, and describing pH-induced changes, from soluble enzymes to integral membrane proteins, including several SARS-CoV-2 targets (40, 55, 75, 89–99). Our present D-NEMD simulations characterise the structural and dynamical responses of the ancestral, Delta, and Omicron spikes to acidic and basic pH shifts, revealing differences in response across the variants. The distinct motion profiles elicited by pH perturbations reveal mechanistic differences in how the ancestral, Delta, and Omicron BA.1 proteins regulate key functional regions, including the RBD, FP, and NTD, with potential implications for viral infectivity and environmental stability. Our findings further reinforce the structural and functional importance of glycans, particularly the *N*-linked glycan at position N234, which actively stabilises the RBD in its “up” conformation during spike activation.

## Results and Discussion

Extensive equilibrium MD simulations, followed by hundreds D-NEMD simulations, were performed to uncover how pH changes impact the functional dynamics of the glycosylated ancestral, Delta, and Omicron BA.1 (hereafter referred to as Omicron) spikes (Figure 1). The Delta variant, which exhibits enhanced transmissibility compared to the ancestral strain (100–102), harbours eight substitutions and two deletions in the spike (57–60). Omicron, a highly transmissible but less pathogenic variant than earlier variants (103–108), including Delta, carries 30 substitutions, three deletions, and one insertion in its spike (61–64).

Long equilibrium simulations of the three fully glycosylated systems in the closed state were conducted under physiological pH (pH ∼ 7) conditions, with all proteins remaining stable, showing structural convergence after ∼100 ns and minimal secondary structure loss after 750 ns (Figures S1-S3). Under equilibrium conditions, the overall dynamic behaviour of the variants closely resembles that of the ancestral protein, with the most notable deviations localised near sites of amino-acid substitutions, deletions, and insertions (Figure S4). However, most of these differences are not statistically significant (as indicated by the high *p*-values shown in Figure S5), with the only exception being the RBDs of Omicron, which exhibit significantly increased dynamics compared to the ancestral protein.

The SARS-CoV-2 spike is extensively glycosylated, typically featuring 22 *N*-linked and at least two variably occupied *O*-linked glycosylation sites per monomer (37, 78, 79, 82, 83). 17 out of the 22 predicted *N*-glycosylation sites were confirmed to be constantly occupied (78–81), while the *O*-linked sites exhibit low occupancy (79, 82, 83). Glycan composition and site occupancy were shown to vary across variants, expression systems, and experimental methods (78, 79, 109, 110). For example, alterations in *N*- and *O*-glycosylation profiles were observed across variants, with variants exhibiting higher resistance to neutralising antibodies (e.g. Omicron) showing a decrease in complex-type glycans with fucosylation and sialylation and an increase in the oligomannose-type glycans compared to the ancestral protein (111). All simulations presented in this work incorporate fully glycosylated models for the spike, reflecting the heterogeneity in composition and occupancy observed in experiments (78, 79). Consequently, the glycan occupancy and composition at each site can vary across monomers (for a detailed description of the glycosylation profiles, see references (56, 112)). In our models, *O-*glycan site occupancy illustrates this variability: monomer A contains glycans at both T323 and S325, whereas monomers B and C only have glycosylation at T323. Similarly, glycan composition differs between monomers, with, for example, the oligomannose linked to N234 containing eight mannose residues and two N-acetylglucosamine residues (Man8GlcNAc2) in monomer A and nine mannoses (Man9GlcNAc2) in monomers B and C (56). Differences in glycan occupancy also extend to the variants; for example, the T19R mutation in Delta ablates the *N-*glycosylation recognition sequon motif at position 17, leading to the absence of an *N-*glycan at this site (111, 113, 114). Analysis of glycan dynamics in the equilibrium simulations of the ancestral, Delta and Omicron spikes revealed that the fluctuations for each glycan are generally similar across the variants, with the *N*-glycans being more flexible than the *O*-glycans (Figure S6). The accessible surface area covered by the glycans was also determined (using various probe sizes, from the radius of a water molecule to that of a small antibody molecule) to assess the extent of the glycan shielding in the ancestral, Delta and Omicron spikes, with no noticeable differences in the protein accessible area between strains (Figure S7).

### pH-induced changes in the ancestral spike: insights from D-NEMD

A total of 576 conformations (i.e. 192 per system for the ancestral, Delta and Omicron spike simulations) were extracted from equilibrium trajectories as starting points for D-NEMD to investigate the structural and dynamic effects of pH changes (Figure S8). The D-NEMD approach, conceptualised by Ciccotti *et al.* (115, 116), integrates equilibrium and nonequilibrium simulations to directly compute (average) system properties under dynamical nonequilibrium conditions (87, 88). The Kubo-Onsager relation is used to compute the response to (either instantaneous or time-dependent) external perturbations (87, 88). D-NEMD tracks the time-resolved structural and dynamic responses of a system, while also assessing the convergence of these responses to ensure robust and reliable results (87, 88). This method is general and highly versatile, accommodating a wide range of perturbations tailored to the biological question of interest. For biomolecular systems, for example, perturbations can include chemical reactions, pH changes, or ligand binding/unbinding (87, 88). For the spike, D-NEMD identified allosteric modulation by the fatty acid binding site (55, 95), showed differences in allosteric responses between variants (117), uncovered how a pH decrease impacts the dynamics of Delta (75), and elucidated how the furin cleavage site is connected to other functionally relevant regions (40).

In this study, the perturbation applied involves altering the protonation states of specific titratable residues in the spikes to simulate acidic and basic pH shifts (see Methods section in Supporting Information for detailed list of the residues changing protonation state). The pH decrease reflects the mildly acidic conditions (pH ∼ 5) found in endocytic and secretory subcompartments, as well as in exhaled particles equilibrated with the typical composition of indoor air (5, 22, 118). In contrast, the alkaline shift (to a pH ∼ 10) mimics the sharp pH increase observed in nascent aerosol particles shortly after exhalation, where pH can exceed 10 (16, 119). By introducing a driving force into the protein (in this case, *via* changes in the protonation states of selected titratable residues), D-NEMD complements and extends equilibrium simulations, unveiling protein responses and dynamic behaviours that are difficult to capture in equilibrium conditions (87, 88). D-NEMD, utilising protonation changes in specific histidines as the external perturbation, was previously employed to investigate the effect of a pH drop on the functional dynamics of the Delta spike, revealing changes in the conformational behaviour of key functional regions (75).

Here, a total of 1152 short nonequilibrium simulations (i.e. 192 simulations per system per pH increase/decrease) were performed, each lasting 20 ns (Figure S8). The applied perturbation, involving a change in the protonation state of specific aspartate, glutamate, histidine, and lysine residues, disrupts the system equilibrium and generates the driving force for the pH-induced structural and dynamic changes to occur as the proteins relax and gradually return to (a new) equilibrium. The evolution of the average response of each spike was derived using the Kubo-Onsager relation (87, 88) by comparing equilibrium and nonequilibrium trajectories at equivalent time points and averaging across all 192 replicates for each system (Figure S8).

In addition to the nonequilibrium simulations, 576 “null perturbation” simulations were conducted for the ancestral, Delta, and Omicron systems (for further details, see Methods section in Supporting Information). These control simulations help distinguish between intrinsic protein motions and natural fluctuations from perturbation-driven responses. This “null perturbation” analysis, as initially introduced by Kamsri *et al.* (90), allows for the removal of the proteins’ equilibrium background motions from the D-NEMD responses, thereby helping to denoise the data and facilitating its analysis and interpretation (87, 89, 90). The convergence and statistical significance of the pH-induced responses were assessed by calculating the standard error of the mean (87, 88) (Figures S9-S14).

Shifts in pH trigger a complex cascade of structural and dynamical changes within the ancestral protein, impacting functionally important regions, such as the RBD and FPPR, to varying extents (Figures 2 and S15-S17). pH decrease and increase elicit distinct patterns of motion and protein responses. For example, low pH induces changes in the sub-domain 1 (SD1) in the CTD, particularly, in the D568- and D574-neighbouring regions and along with the upper portion of the FPPR (also frequently referred to as fusion domain), S2 cleavage site and the area around E1111 at the base of the head domain. In contrast, high pH triggers responses in the segments surrounding K733, including E654-S686 in sub-domain 2 (SD2), V687-A701, L861-I870, and A771-E780, as well as around K129 in the NTD, specifically residues V120-N125 and S161-P174.

**Figure 2.**
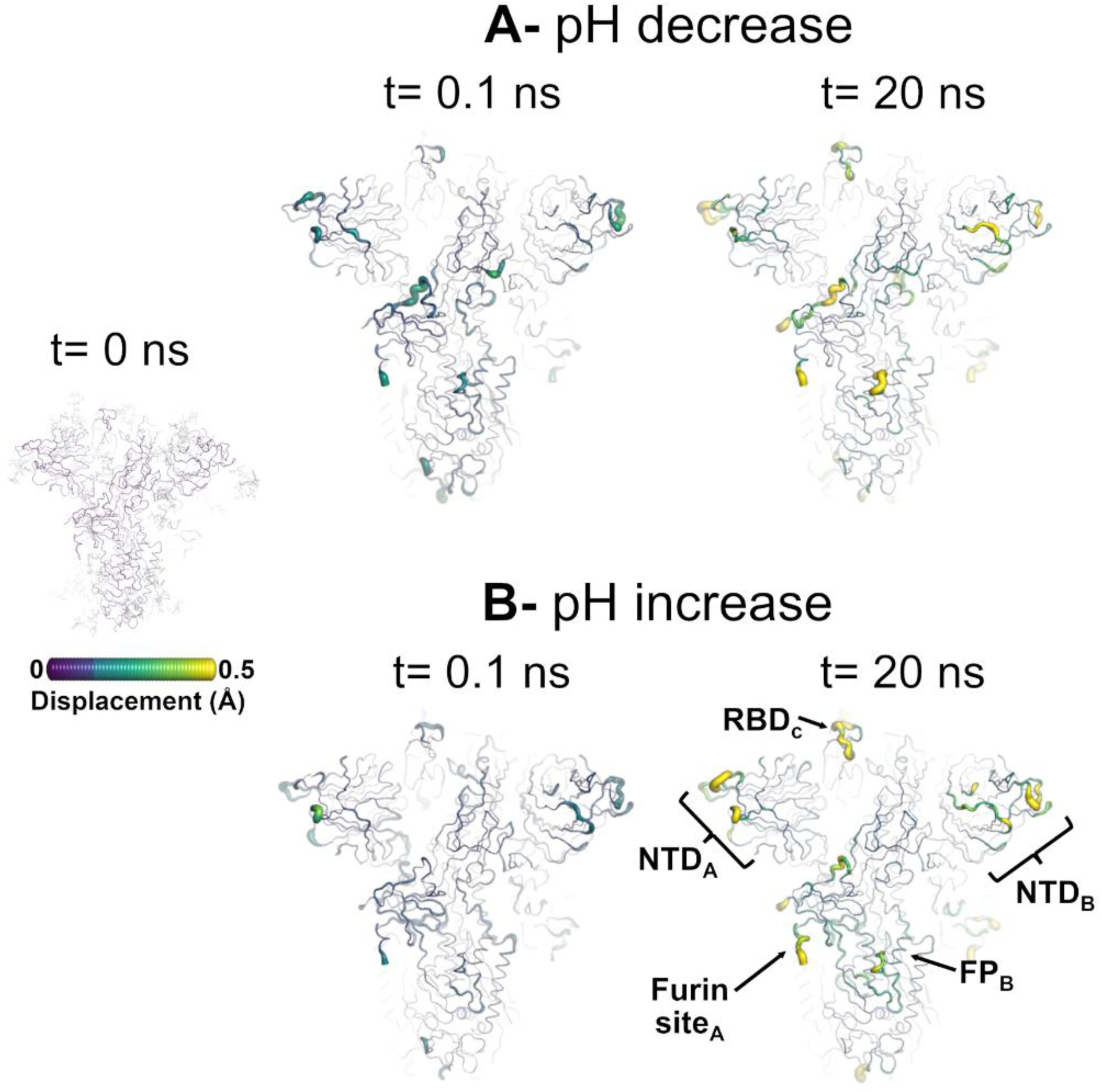
Structural response of ancestral spike to a pH decrease and increase. **A**) Average C_α_ displacements at *t* = 0.1 and 20 ns after a pH decrease. **B**) Average C_α_ displacements at *t* = 0.1 and 20 ns after a pH increase. The norm of the average C_α_ displacement vectors between the nonequilibrium (at low/high pH) and equilibrium (at physiological pH) simulations was calculated for each C_α_ atom using the Kubo-Onsager relation (87, 88). The final displacements reported correspond to the average D-NEMD responses after removing the intrinsic protein fluctuations via the “null perturbation” analysis. Both the cartoon thickness and structure colours (scale on the left) indicate the average responses to a pH change. Glycans are depicted as light grey sticks in the leftmost image but were omitted from the remaining panels (showing the responses at *t =* 0.1 and 20 ns) to facilitate visualisation. Each RBD, NTD, furin site and FP are subscripted by their chain ID (A, B or C). This figure illustrates the protein’s response from the perspective of FP_B_ (for the pH-induced changes at other time points and/or from the viewpoint of FP_A_ and FP_C_, refer to Figures S15-S17). Overall, comparable response patterns are observed across the three individual subunits

Both acidic and basic shifts induce structural responses within the RBD, with especially pronounced alterations in the receptor-binding motif (RBM) (Figures 2 and S15-S17). The RBM plays a critical functional role in the spike mechanism of action, as it directly mediates binding to the ACE2 receptor on human cells, thereby facilitating viral attachment and invasion (25). These results align with recent equilibrium simulation studies suggesting that pH modulates spike behaviour by inducing highly localised structural and dynamic changes under moderately acidic and basic conditions, with the RBD being among the most pH-sensitive regions (72, 74).

Notably, mild conformational responses to both acidic and alkaline pH changes are also detected near the heme/biliverdin binding site, specifically involving residues E169-F186 in the NTD (46, 47), with these changes being slightly more pronounced under alkaline conditions. Although the specific responses around this site can vary between monomers, their magnitude and spatial location remain relatively the same across pH conditions (Figures 2 and S15-S17). This site, located on the distal face of the NTD, has been experimentally shown to bind heme and its metabolites, biliverdin and bilirubin (46–48). While its biological role remains unclear, experimental evidence suggests that biliverdin binding to this site interferes with the attachment of specific neutralising antibodies, suggesting a potential role in immune evasion (46).

The pH-induced structural responses initially manifest as subtle alterations localised in the regions directly interacting with residues that changed protonation state (Figures 2 and S15-S17). However, as the 20 ns simulations progress, these responses intensify and gradually propagate throughout the protein structure, extending beyond the initially affected residues and triggering broader conformational responses (Figures 2 and S15-S17). An example of this behaviour can be seen around D574 in the SD1 following a pH decrease (Figure S18), with subtle responses observed around this residue immediately after its protonation state change; as time progresses and the protein adapts to the residue’s new (neutral) state, a steady growth in the responses around this residue is observed, with the changes eventually reaching the upper part of the FPPR in S2 (Figures S15-S18).

While the amplitude of the responses often varies between monomers, the overall pattern and sequence of structural changes remain generally the same across all chains (Figures 2 and S15-S17). These amplitude differences likely stem from asymmetries in protein structure and dynamics caused by variations in glycan occupancy and composition between monomers (for details on the protein’s glycosylation profile, see Supporting Information and Casalino *et al*. (56)). As mentioned above, the models used here exhibit distinct glycosylation patterns across the spike monomers, reflecting the site-specific heterogeneity observed in the experimental glycosylation data (78, 79, 109, 111).

Using D-NEMD, we can quantify not only the magnitude of the (statistically significant) structural responses induced by a perturbation but also determine the direction of these motions (Figures 3 and S19-S21). This is achieved by computing the average displacement vectors of C_α_ atoms between the equilibrium and nonequilibrium trajectories at corresponding time points (87). The average response values reported in Figures 2 and S9-S18 represent the norms (magnitudes) of these average C_α_ displacement vectors.

**Figure 3.**
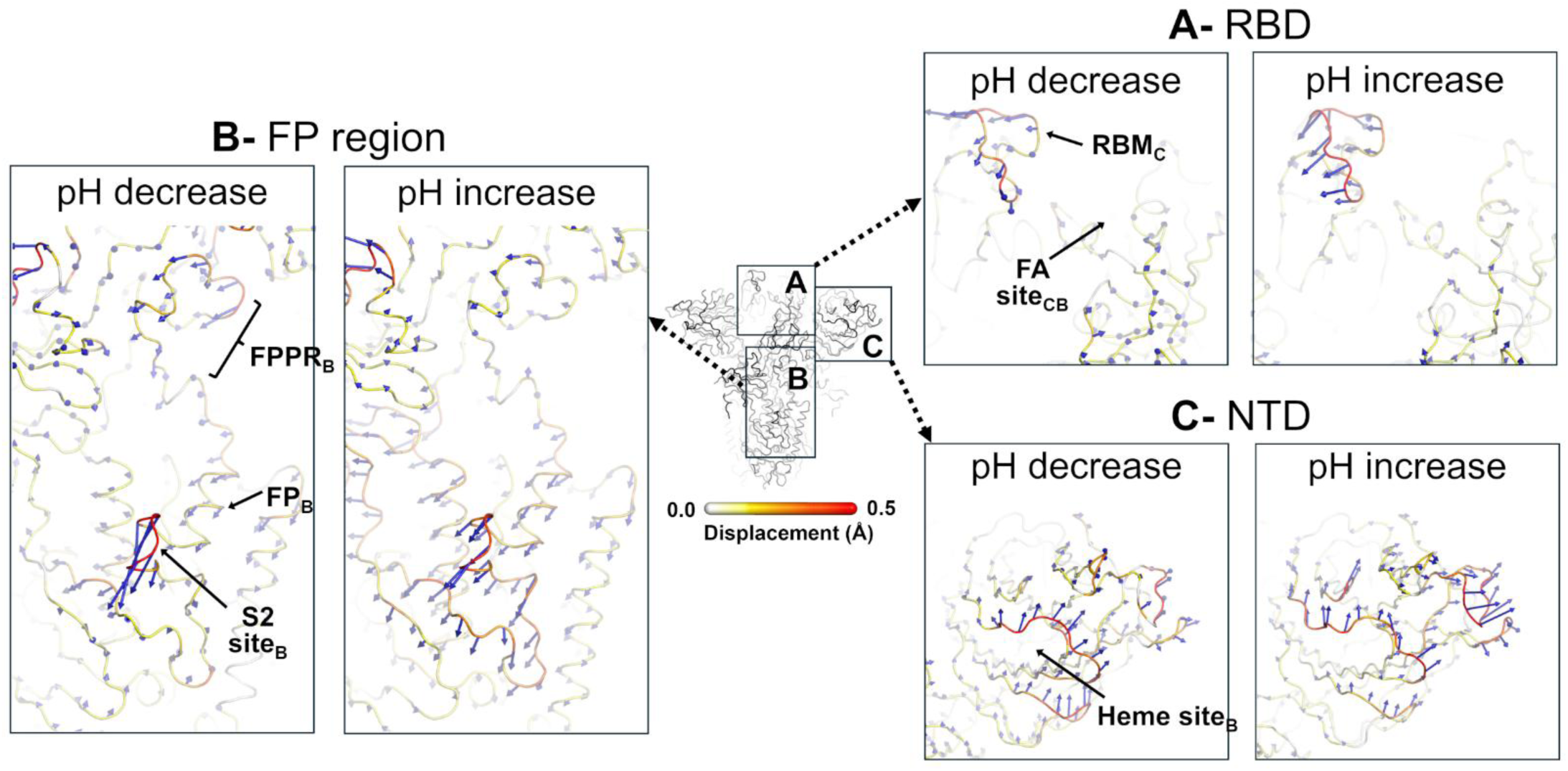
Directionality of the pH-induced structural responses in the ancestral spike at *t* = 20 ns after the pH shift. **A**) Direction of the RBD motions in response to a pH decrease and increase. **B**) Direction of motion of the FP-surrounding regions in response to a decrease and an increase in pH. **C**) Direction of the NTD motions in response to a pH decrease and increase. The blue arrows correspond to the average C_α_ displacement vectors (calculated between each pair of equilibrium and nonequilibrium trajectories over the 192 replicas for each pH shift) after removing the intrinsic protein fluctuations via the “null perturbation” analysis (see Supporting Information for more details). Vectors with a length ≥0.08 Å are displayed as blue arrows with a scale-up factor of 10. Average C_α_ displacements (i.e. the magnitude of the average C_α_ displacement vectors) are represented on a white-yellow-orange-red scale. Glycans were omitted from panels A, B and C to facilitate visualisation, but were included in all the simulations. All regions labelled are subscripted with their chain ID (A, B or C). This figure illustrates the direction of the motions from the perspective of the FP_B_, NTD_B_ and RBD_C_ (for the direction of motions from other viewpoints, see Figures S19-S22).

As illustrated in Figure 3, the response of the ancestral spike is (highly) susceptible to pH variations, with distinct motion patterns emerging depending on the environmental pH. Some regions show structural responses only under specific pH conditions, while others react to both acidic and alkaline shifts, but with differences in the direction of their movements. For example, the RBM_C_ displays notable variations in both amplitude and direction of its motion under different pH conditions, with a well-defined upward movement toward the solvent under acidic conditions and shifting inward under alkaline conditions (Figure 3A). The strong sensitivity and responsiveness of the RBM suggest that this region is finely tuned to adapt to changes in environmental pH. Given its role in mediating contact with the human ACE2 receptor (25), the distinct pH-dependent directional dynamics observed is likely to influence the virus’s ability to bind to host cells, potentially affecting its overall infectivity across different pH environments. Experimental studies have shown that the stability and infectivity of SARS-CoV-2 can vary between acidic and alkaline conditions (16, 22, 24), with infectivity remaining relatively stable within pH 5.6-9, but sharply declining above pH 9.5 (16–18); multiple factors can contribute to this (including effects on the viral membrane), but the results here indicate potentially significant pH effects on the spike, particularly those on RBD structure and dynamics.

Three other spike regions, namely the FPPR, SD1 and SD2, also show distinct movement patterns following acidic and basic pH changes, with the first two exhibiting more pronounced motions as the pH decreases and the latter generally demonstrating stronger responses under alkaline conditions (Figures 3 and S19-S22). For instance, FPPR_C_ and SD1_A_ undergo an upward displacement in acidic pH but show minimal movement at higher pH (Figures S20 and S22). In contrast, the final part of SD2_A_ (residues E661-Q675) moves outward under alkaline conditions but shows essentially no change at low pH (Figure S20). The FPPR is a ∼25-residue segment (residues 828-853) located immediately downstream of the FP, and it plays a critical role in the spike structural rearrangements required for membrane fusion (39, 120). Functioning as a flexible hinge, the FPPR facilitates the transition from the prefusion to the postfusion spike conformation (39, 77, 120, 121). In addition to its mechanical role, this region has also been proposed to act as a pH switch, potentially modulating the RBD accessibility in acidic conditions (9, 120). The SD1 region, positioned between the RBD and FPPR, is involved in the structural rearrangements required for the closed RBD-down to the open RBD-up conformation (39, 120). This region acts as a structural relay between the RBD and FPPR, detecting structural changes on either side and facilitating coordinated conformational transitions (39, 120). SD2 also interacts closely with the FPPR and is thought to contribute to the structural stability of the cleaved spike trimer, particularly in the prefusion state, by helping preserve the integrity of the S1 subunit and delaying its premature shedding (120, 121). The salt-bridge between D614 in SD2 and K854 in the FPPR was shown to be particularly important for this stabilisation, with mutations disrupting this interaction, most notably D614G (a defining feature of all major variants of concern), altering the position and dynamics of both the FPPR and 630-loop (77, 120–124). This facilitates the transition towards the open RBD-up conformation, thus enhancing the protein’s ability to engage with host receptors (77, 120–124). Consistent with the behaviour of the RBM described above, the pH-responsive changes observed in FPPR, SD1, and SD2 in this study suggest that these regions contribute to pH sensing and help tune the spike’s functional dynamics to different pH environments (Figures 3 and S19-S22).

Finally, it is also worth noting that certain regions of the spike exhibit similar structural responses to both acidic and basic pH shifts. An example of these is the heme/biliverdin binding sites located within the NTDs. While the behaviour of these sites can vary between individual monomers, the direction and magnitude of movement within each site remain essentially the same, regardless of whether the pH increases or decreases (Figures 3 and S19-S22). These findings suggest that, while this region is clearly sensitive to pH fluctuations, it does not discriminate between acidic and alkaline conditions. Such behaviour points to a potential role in general pH sensing, in which the region undergoes structural responses regardless of whether the environment becomes more acidic or more alkaline. Other spike segments, such as regions flanking the cleavage sites and FP, along with specific residues including D290, D385, D586, and D614, have also been proposed to contribute to pH sensing in the spike (66–68).

### Dynamic responses of the Delta and Omicron spikes to pH changes

D-NEMD simulations were also carried out for Delta and Omicron spike proteins to compare their structural responses to pH changes with those of the ancestral protein. Such comparison enables us to assess how substitutions, deletions and insertions in these variants influence the protein’s response dynamics under varying pH conditions. Again, 192 nonequilibrium simulations, each lasting 20 ns, were conducted for each variant (in this case, Delta and Omicron), mimicking the transition from physiological pH to acidic and basic environments (Figure S8). To account for intrinsic fluctuations in Delta and Omicron in equilibrium (i.e. at physiological pH), an additional 192 “null perturbation” simulations were performed per system. The average structural response of each variant was computed using the Kubo-Onsager relation (87, 88), with results averaged across all 192 replicates and corrected for baseline fluctuations at physiological pH using the “null perturbation” data (Figures S11-S14).

Comparison of the dynamical responses of the ancestral and variant spikes reveals that Omicron exhibits increased sensitivity to acidic conditions compared to the ancestral and Delta proteins, but shows generally attenuated responses under alkaline conditions relative to these variants (Figures 4, S15-S17 and S23-S28). Interestingly, although the number of residues undergoing protonation changes upon pH shifts increases progressively from ancestral to Delta and then to Omicron (see Supporting Information for a list of the residues changed), the magnitude of the structural responses does not simply correlate with the number of affected sites. Notably, Omicron exhibits approximately twice as many protonation changes as Delta, yet displays both the most pronounced conformational responses at low pH and the most attenuated responses at high pH among the three systems (Figure 4).

**Figure 4.**
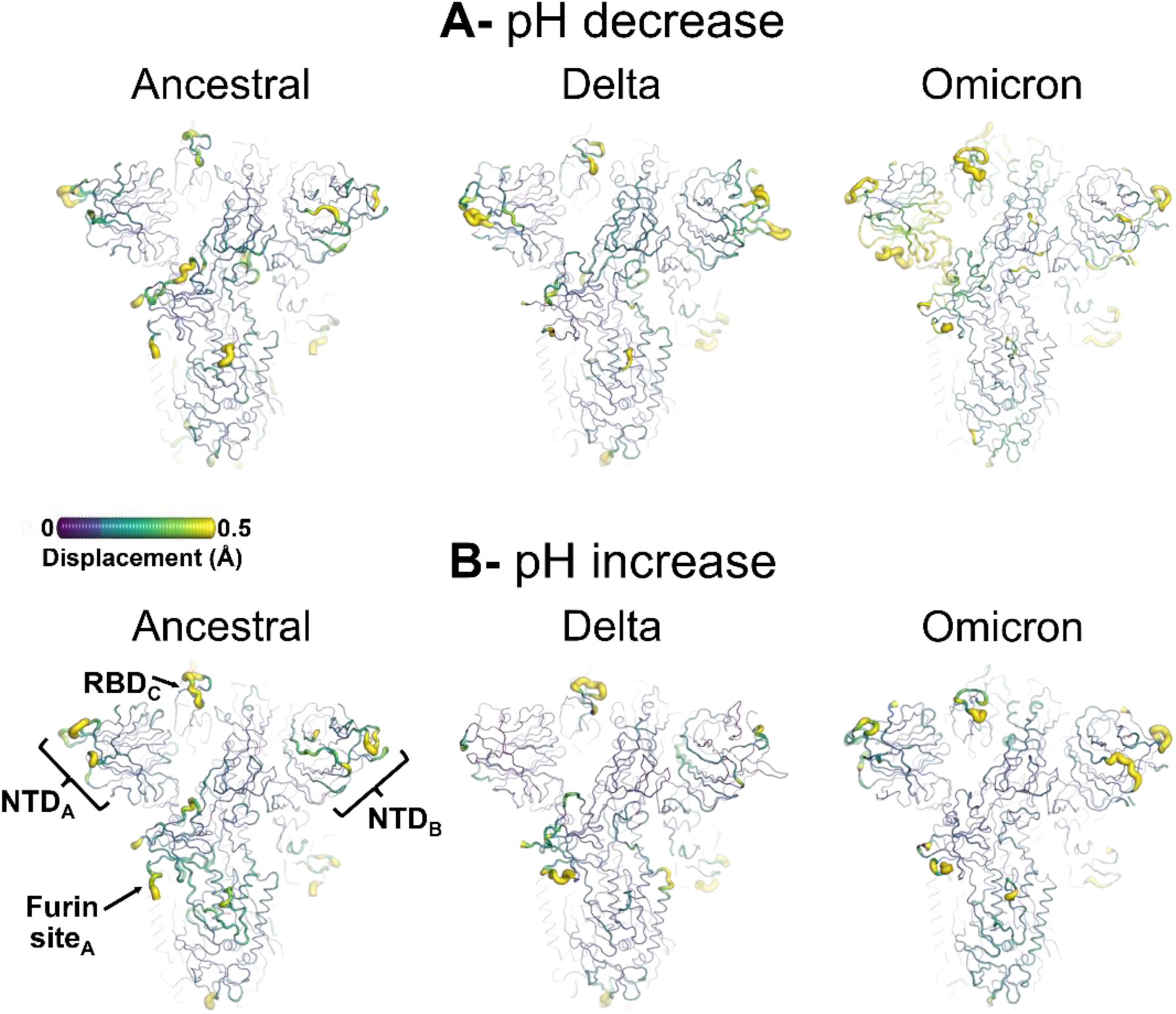
Structural response of the ancestral, Delta, and Omicron spikes to pH changes. pH-Induced structural changes in the ancestral and Delta and Omicron spikes following a **A**) pH decrease and **B**) pH increase. The norm of the average C_α_ displacement vectors between the D-NEMD (i.e. at low/high pH) and equilibrium (i.e. at physiological pH) simulations was calculated for each C_α_ atom using the Kubo-Onsager relation (87, 88). The final displacement values represent the averages obtained from the 192 pairs of simulations for each system, after removing the intrinsic protein fluctuations *via* the “null perturbation” analysis (see Supporting Information for details). The cartoon thickness and structure colours indicate the average C_α_ positional displacements. Glycans were omitted to facilitate visualisation, but were present in all equilibrium and nonequilibrium simulations. Each structural motif is subscripted with its chain ID (A, B or C). This figure shows the spikes’ responses from the viewpoint of NTD_A_, NTD_B_ and RBD_C_ (see Figures S15-S17 and S23-S28 for the responses from the viewpoints of NTD_C_, RBD_A_ and RBD_B_, respectively).

Among the systems analysed, Omicron exhibits the strongest response to acidic pH changes, whereas Delta and the ancestral spike show generally similar, but less marked, sensitivity to this shift (Figures 4, S15-S17 and S23-S28). The changes in functional dynamics observed in Omicron, along with its enhanced responsiveness under acidic conditions, suggest that this variant may have specifically adapted to operate more effectively in low-pH environments. This is particularly significant given experimental evidence indicating that, unlike earlier variants, Omicron preferentially enters host cells *via* receptor-mediated endocytosis within acidic endosomes (105, 106, 108, 125–129). The SARS-CoV-2 virus can infect host cells using two distinct membrane fusion pathways, each operating under different pH conditions (6, 8, 9). One pathway involves fusion at the plasma membrane surface under physiological pH conditions (pH ∼ 7.0-7.4), whereas the alternative route occurs within endosomal compartments, where the environment is acidic (pH ∼ 5.0-6.5) (6, 8, 9). Experimental studies have shown that the choice of entry pathway depends mainly on the relative expression levels of the type II transmembrane serine protease, TMPRSS2 (130, 131). In cell types with high TMPRSS2 expression, the virus predominantly enters *via* plasma membrane fusion (130–133), whereas in cells with low or absent TMPRSS2 levels, it relies on the endosomal route, which depends on endosomal proteases such as cathepsin L (130, 131, 134). Experimental evidence also indicates that different SARS-CoV-2 strains preferentially utilise one cellular entry pathway over the other (127, 135). Both the ancestral and Delta strains primarily rely on the TMPRSS2-dependent plasma membrane entry route (100, 102); in contrast, Omicron shows reduced reliance on TMPRSS2, instead favouring endosomal entry, reflecting a shift in cellular tropism compared to earlier strains (105, 106, 108, 125–129).

For Omicron, regions such as the RBD, FPPR, SD1, SD2 and specific NTD segments display enhanced responsiveness compared to the ancestral and Delta protein, with these regions consistently undergoing larger changes across all three Omicron monomers (Figures 5 and S29-S36). Among these, the RBM within the RBD of monomers A and C exhibits a particularly pronounced upward rotation under acidic pH conditions (Figures 5 and S35). Omicron contains 15 amino-acid substitutions in the RBD relative to the ancestral spike (61–64), with the majority clustered in the RBM, specifically N440K, G446S, S477N, T478K, E484A, Q493R, G496S, Q498R, N501Y, and Y505H. These substitutions result in the addition of five extra positive charges in the RBD alone at low pH. These mutations collectively enhance spike binding to ACE2 receptors and contribute to evading antibody neutralisation (63, 136, 137). Our findings here demonstrate that these substitutions also significantly influence RBD dynamics and its responsiveness to acidic conditions, with the direction of the observed conformational shift suggesting increased tendency for this region to transition from a closed (RBD in “down” position) to the open (RBD in “up” position) conformation, a structural rearrangement that is essential for engaging with host receptors (37–39).

**Figure 5.**
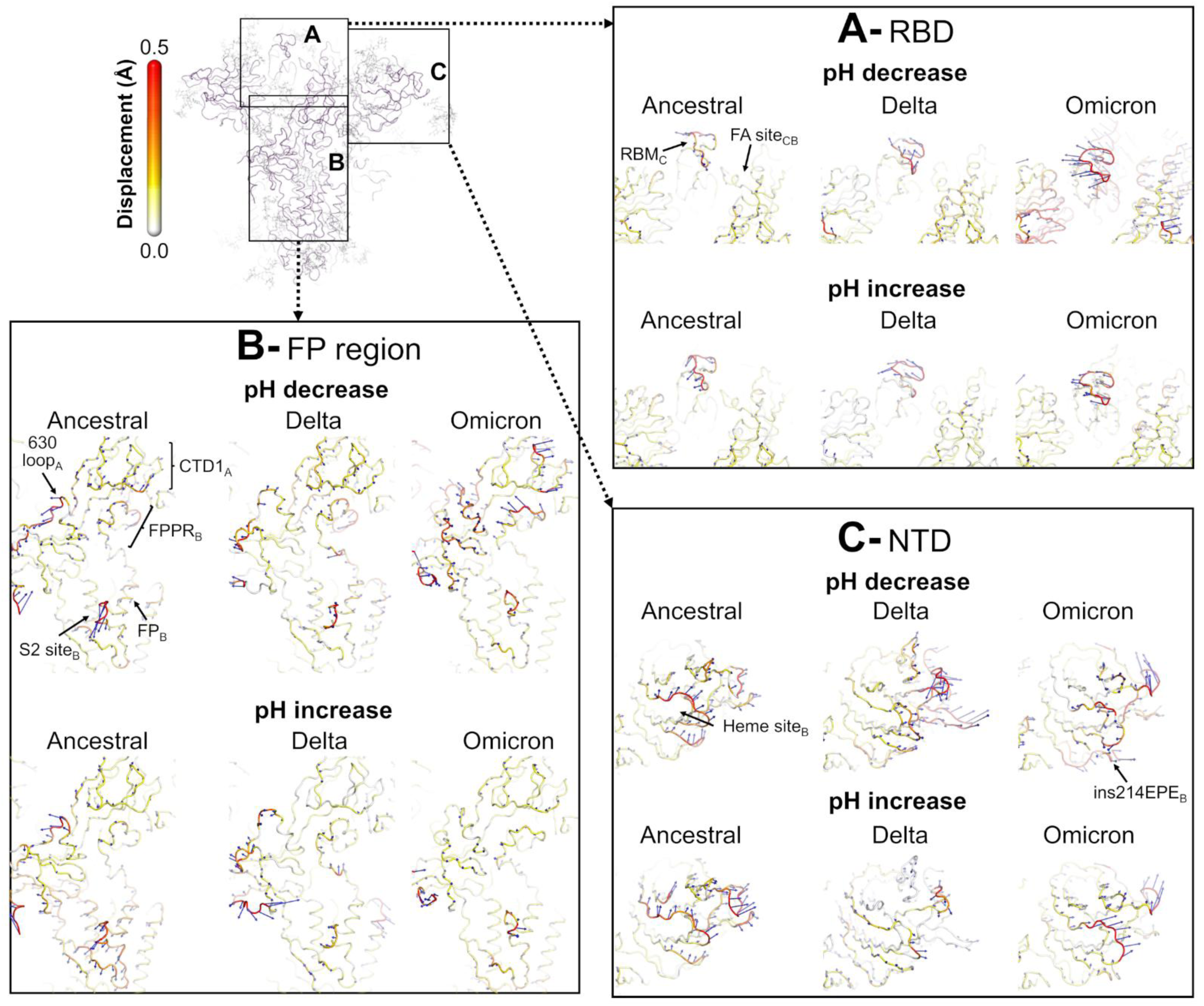
Close-up view of the direction of the RBD, FP-region and NTD responses to pH changes across the ancestral, Delta, and Omicron spikes. Comparative analysis of the pH-induced structural changes in ancestral, Delta, and Omicron 20 ns following a pH decrease/increase. Detailed view of the structural response of the **A**) RBD_C_, **B**) regions surrounding the fusion peptide_B_, and **C**) NTD_B_. The direction and amplitude of the average C_α_ displacements 20 ns after the pH decrease/increase are shown. The final displacement values represent the averages obtained from the 192 pairs of simulations for each system after removing the intrinsic protein fluctuations via the “null perturbation” analysis (see Supporting Information for details). Vectors with a length ≥0.08 Å are displayed as blue arrows with a scale-up factor of 10. Average C_α_ displacements (i.e. the magnitude of the average C_α_ displacement vectors) are represented on a white-yellow-orange-red scale. Glycans were omitted from panels A, B and C for clarity, but were included in all simulations All regions labelled are subscripted with their chain ID (A, B or C). This figure illustrates the direction of the motions from the perspective of the NTD_B_, FP_B_, and RBD_C_ (for the direction of motion from the viewpoints of NTD_A_, RBM_A_, FP_A_, RBM_B_, NTD_C_ and FP_C_, see Figures S19-S21 and S29-S34).

In addition to the RBMs, the free fatty acid (FA) binding site located at the interface between adjacent RBDs (45) also show increased responses to low pH conditions in Omicron compared to the ancestral and Delta spikes, with the most pronounced changes occurring in FA_CB_ site, located at the interface between chains B and C (Figures 4, S15-S17 and S23-S28). Building on the observed upward rotation of the RBMs, in Omicron, the RBDs forming each FA site exhibit opposing directional motions, suggesting increased separation between the two domains and a loosening of this interface (Figures 5 and S35-S36). Contrary to the RBM, Omicron does not harbour any amino-acid substitutions directly within the FA site itself, although several mutations are located in close proximity to the pocket, namely G339D, S371L, S373P, and S375F (61–64). Some of these mutations enhance the stability and structural integrity of the RBD while others contribute to immune escape (63, 136, 138). Despite the nearby mutations, experimental evidence indicates that binding of free fatty acids to this site remains intact in Omicron, unaffected by the surrounding residue changes (139). Although the precise role of the FA site in the spike’s mechanism of action remains uncertain, binding of free fatty acids to this site has been proposed to function as a molecular switch, preventing premature spike activation during viral assembly, shielding key epitopes from neutralising antibodies, and modulating viral entry in response to environmental or biochemical conditions (40, 45, 48, 139, 140). Experimental studies using surface plasmon resonance and cryo-EM demonstrated that binding of free fatty acids, as well as various other molecules, to the FA site stabilises the spike in a locked conformation (in which all RBMs are occluded) (139, 141–144), potentially preventing premature engagement with host receptors and prematurely exposing the RBD surfaces to neutralising antibodies. The FA site was also shown to be allosterically coupled to key functional regions of the spike, including the FP region and furin cleavage site, with the disruption or ligand binding/unbinding at this site influencing the dynamics of these regions (40, 55, 95, 117). The FP is a short, hydrophobic helix that inserts into the host cell membrane to initiate fusion (37–39), while the furin cleavage site (a polybasic insertion located at the S1/S2 junction (37, 38)) is involved in protein activation (42), with its removal reducing viral infectivity (8, 40, 42, 132, 145, 146). Intriguingly, the FA site is highly conserved among the spikes of highly pathogenic β-coronaviruses, including SARS-CoV, MERS-CoV and SARS-CoV-2, but is absent in less virulent strains, such as the common cold-causing HCoV-HKU1 virus (139).

In the NTD, the regions interacting with L210-I216 also display heightened responses to acidic conditions in Omicron compared to the ancestral and Delta proteins (Figures 5 and S35-S36). Residues L210-I216 are part of a flexible peripheral loop, located near the NTD antigenic supersite (147–151). Omicron harbours several sequence changes in segments neighbouring this supersite, which are thought to contribute to immune evasion (107, 152, 153). These include modifications in the L210-I216 segment itself, notably an asparagine-to-isoleucine substitution at position 211 (N211I), a deletion at position 212 (L212Δ), and a glutamate-proline-glutamate insertion at position 214 (ins214EPE) (61–64). Interestingly, our findings here indicate that these mutations (particularly ins214EPE) not only alter the dynamics of the lower part of the NTD but also render it highly sensitive and responsive to acidic pH. Nonetheless, the exact mechanistic basis for the emergence of this novel pH-responsive switch within the NTD remains to be elucidated.

The top part of the FPPR and SD1 also show stronger and clearer responses in Omicron compared to the ancestral and Delta variants following a pH decrease (Figures 5 and S35-S36). However, the direction of their motions is more diverse across the monomers, with FPPR_B_ and the CTD region directly interacting with it, SD1_A_, showing similar upward displacements, and the remaining ones displaying opposite direction motions (Figures 5 and S35-S36). As outlined above, the FPPR has been suggested to act as a pH-sensitive flexible hinge (9, 39, 77, 120, 121), while the SD1 functions as a structural relay between the FPPR and the RBD (39, 120), with both regions coordinating the conformational changes required for the spike’s transition between RBD-down and RBD-up conformations. In Omicron, there are no amino-acid substitutions directly within the FPPR (although there are some, such as N856K, D614G, and Q954H, nearby), and there is a single threonine-to-lysine change at position 547 in the SD1 (61–64). Together, these mutations were shown to improve protein stability, facilitate more efficient viral transduction into host cells, and increase infectivity (62, 137, 154–157).

Finally, the 630-loop within SD2 also generally displays more pronounced conformational changes in response to acidic pH in Omicron compared to the ancestral and Delta variants. Notably, the direction of motion of this loop diverges significantly between Omicron and the ancestral protein, with the first showing an upward shift toward the nearest NTD, and the second displaying a downward rotation (Figures 5 and S35-S36). In contrast, the 630-loop in the Delta spike exhibits consistently smaller conformational responses to acidic pH compared to both the ancestral and Omicron variants, with its motions showing less well-defined directionality. The 630-loop was shown to play a critical role in stabilising the spike’s prefusion conformation, helping to preserve its structural integrity prior to host receptor binding. In cryo-EM structures of the RBD-down conformation of the ancestral spike, this loop is relatively flexible and disordered, contributing to the destabilisation of the spike (39, 120). Neither Delta nor Omicron contain mutations within this loop; however, both variants feature the D614G substitution located nearby (57–64). In the presence of this mutation, the 630-loop was shown to adopt a well-ordered structure and form tighter interactions with adjacent regions, thereby enhancing protein stability and preventing premature RBD opening (39, 77, 120). Experimental structural data further reveal that upon the RBD’s transition from the closed to the open conformation, the 630-loop, along with the FPPR and SD1, undergoes coordinated structural rearrangements (39, 77, 120). Interestingly, our findings here reveal that the conformation and motions of SD1, 630-loop and FPPR, differ not only between variants, but also in response to pH changes, with the most pronounced effect observed in Omicron under acidic conditions. These results underscore the dynamic interplay among these regions and highlight their sensitivity to environmental factors, such as pH.

In contrast to the conformational changes observed under acidic conditions, Omicron’s response when exposed to increased pH is markedly different, with this variant exhibiting generally attenuated reactions compared to the ancestral and Delta proteins (Figures 5 and S35-S36). This unexpected observation suggests that Omicron’s functional dynamics is less affected by alkaline conditions than those of previous variants, potentially reflecting evolutionary adaptations that enhance structural stability and reduce the likelihood of early inactivation at high pH levels. These diminished responses are particularly evident in key functional regions such as the FPPR, S2’ cleavage site, 630-loop, and SD2 (Figures 5 and S35-S36). In contrast, the response of the Delta spike to alkaline pH shifts is less clear, displaying a more nuanced profile across these regions than Omicron, remaining generally closer to the ancestral protein (Figures 5 and S35-S36). While certain regions of the Delta spike, such as the SD2, display attenuated conformational motions in response to a pH increase, others, including the FPPR and RBM, undergo more pronounced changes compared to the ancestral protein. This suggests a more complex and region-specific response profile to basic environments for the Delta spike. Alkaline pH conditions, like acidic ones, are also biologically relevant and can arise, for example, in aerosol particles, where experimental studies have shown that pH levels fluctuate after exhalation and during airborne transport (16–23), including shifting toward very alkaline values (16–18, 21). The high pH (experimentally) observed in respiratory aerosols originates from the mucosal fluids they derive from, such as saliva and lung fluid, which naturally contain elevated levels of bicarbonate (158, 159). Upon droplet exhalation from the CO_2_-rich environment of the lungs into the environment, bicarbonate is converted into dissolved carbon dioxide, which subsequently evaporates, causing the initially-neutral pH within the aerosol to rise, creating an alkaline environment (18). The maximum pH reached inside respiratory aerosols and the time required to reach it remain uncertain, likely depending on factors such as droplet size, ambient concentration of carbon dioxide and the presence of carbonic anhydrase enzymes (119, 160); however, some experimental studies have reported pH values > 10 (16, 119). Experimental studies have also demonstrated that SARS-CoV-2 infectivity is influenced by pH (16–18, 21, 24), with elevated pH conditions leading to a reduction in viral infectivity (16–18). Using the advanced CELEBS (Controlled Electrodynamic Levitation and Extraction of Bioaerosol onto Substrate) technique (161, 162), Haddrell and coworkers showed that the Delta variant is more sensitive to alkaline conditions and exhibits reduced aerosol stability compared to earlier variants (17). The distinct conformational responses to pH increases observed here in the ancestral and Delta spikes suggest differing sensitivities to alkaline conditions, broadly aligning with the experimental findings (17). However, a direct comparison between the molecular motions reported here and the infectivity loss observed experimentally is challenging due to fundamental differences in scale, resolution, system complexity, and measured observables between the respective approaches.

More recently, observations (using the CELEBS technique again) indicate that Omicron BA.2 shows increased resistance to high pH conditions compared to Delta (18). Given that most mutations present in Omicron BA.2 are also found in Omicron BA.1 (the focus of this work), it is reasonable to consider that both variants may share enhanced resistance to high pH relative to Delta. Notably, Omicron BA.1 and BA.2 have more than 20 spike mutations in common (163), namely G142D, G339D, S373P, S375F, K417N, N440K, S477N, T478K, E484A, Q493R, Q498R, N501Y, Y505H, D614G, H655Y, N679K, P681H, N764K, D796Y, Q954H, and N969K. However, caution is warranted when extrapolating Omicron BA.1 behaviour from data on BA.2, as although sharing most spike mutations, the two variants also harbour unique sequence changes that may influence their biological properties (163). Specifically, Omicron BA.1 contains seven distinct spike substitutions, three deletions and one insertion (namely A67V, H69Δ-V70Δ, T95I, V143Δ-Y145Δ, N211I, L212Δ, EPE insertion in position 214, S371L, G446S, T547K, and L981F), while Omicron BA.2 features five unique spike mutations and two deletions (T19Δ, L24Δ-P26Δ, A25S, V213G, T376A, D405N and R408S) (163). These sequence changes likely contribute to variant-specific differences in host cell entry, transmissibility, pathogenicity, and immune resistance (164–166). They also underscore the need for variant-specific analyses of pH effects. Nonetheless, the reduced pH sensitivity observed in D-NEMD for Omicron BA.1 at high pH (Figures 5 and S35-S36) may reflect a shift in this variant’s environmental resilience, with potential functional implications, contributing to its distinct infectivity profile.

The pronounced amplitude and clear directionality of the responses observed in the RBM, FPPR, SD1 and NTD following a pH decrease in Omicron suggest that the spike undergoes large conformational rearrangements, consistent with a shift from a “down” to an “up” RBD state. To further characterise the nature of these structural changes, inter-RBD, inter-NTD, and RBD-helical core distances were quantified across the equilibrium (at physiological pH) and nonequilibrium (at low and high pH) simulations for the ancestral, Delta and Omicron variants (Figures S37-S39). As illustrated in Figures S37-S38, similar distances between the centres of mass of neighbouring RBDs and NTDs across the equilibrium and D-NEMD trajectories are observed for the ancestral and Delta spikes, indicating that no large-scale structural rearrangements occur in these systems in response to pH changes. In contrast, Omicron displays a marked increase in these interdomain distances during D-NEMD under acidic conditions compared to physiological pH, an effect not observed at high pH (Figure S39). This increase in the distances at low pH is most pronounced between RBD_C_–RBD_A_ and NTD_A_–NTD_B_ (Figure S39). This distance-based analysis further emphasises the distinctive pH-sensitive conformational behaviour of Omicron and its enhanced responsiveness to acidic conditions seen above.

Cryo-EM studies (e.g. (9, 26, 37, 39, 44, 48, 80, 167–173)) and single-molecule FRET imaging (174) have shown that the spike adopts a range of conformations, from a locked and closed trimer to open states, featuring one, two, or all three RBDs in the “up” position. Snapshots were extracted along the D-NEMD trajectories following a pH decrease to obtain a detailed, atomic-level characterisation of the conformational motions of Omicron as it adapts to low pH conditions. Figure 6 illustrate the structural rearrangements in Omicron following a decrease in pH, starting from a closed spike conformation in which all three RBDs are in the “down” position at the beginning of the nonequilibrium simulations. Over time, the spike transitions from a closed to a partially open state, with one RBD (specifically RBD_B_ or RBD_C,_ depending on the trajectory) shifting into an “up” conformation. This transition, which makes the RBM accessible for binding to ACE2, is an essential step in viral entry into host cells (37–39). The conformational response of the Omicron is accompanied by a reorganization of its inter- and intra-RBD salt bridges, including those involving D364, R457, K458, and K462 (Figure S40). The breaking of these interactions has previously been observed by Sztain *et al.* in their weighted ensemble simulations, where disruption of these interactions was linked to the RBD opening pathway (85).

**Figure 6.**
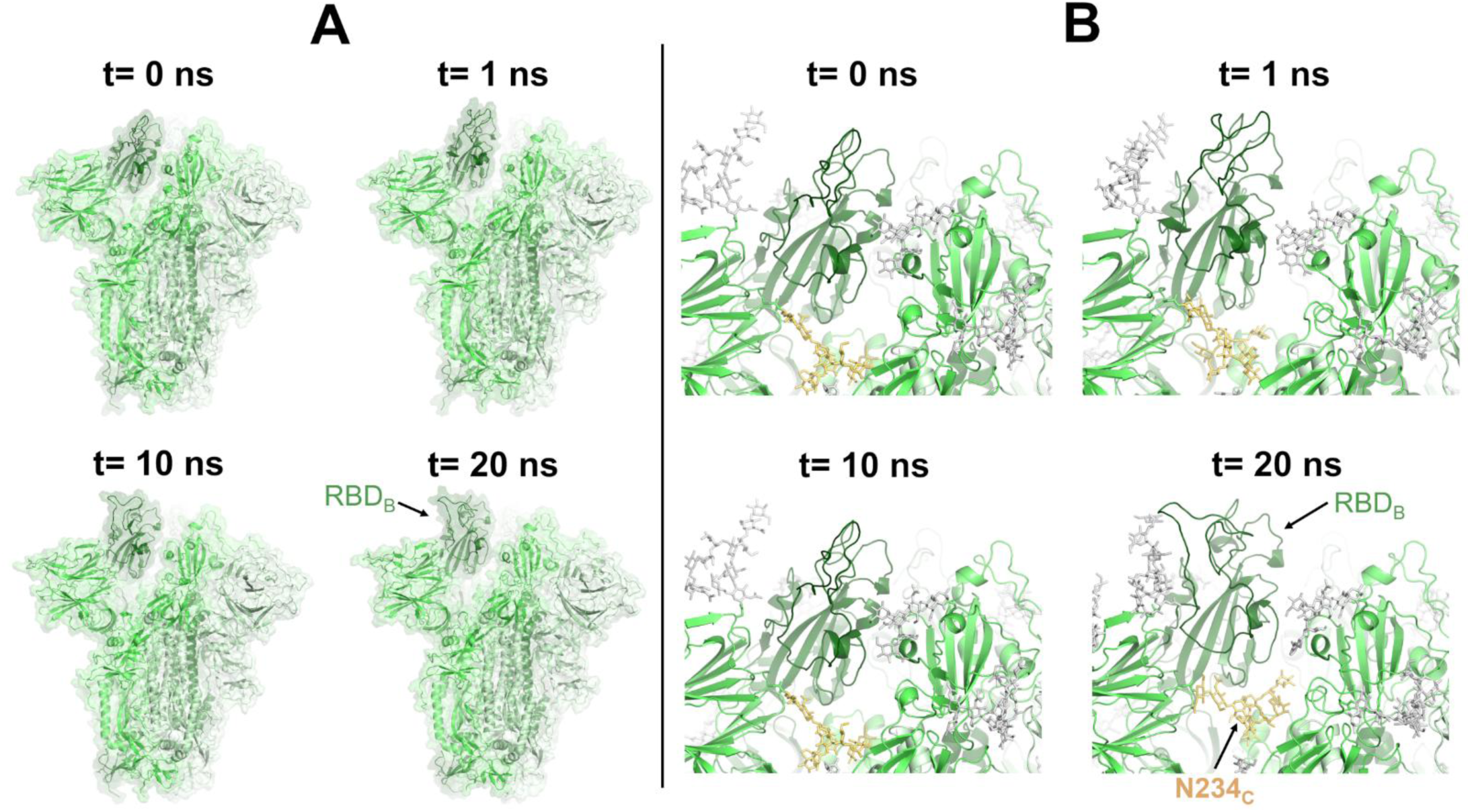
Closed-to-open transition of Omicron RBD following a pH decrease. **(A)** Snapshots along a nonequilibrium trajectory of Omicron following a pH decrease, during which one of the RBDs initiates a closed-to-open transition. Frames at the beginning (t = 0 ns), and at 1 ns, 10 ns and 20 ns after the pH drop are shown. Each spike monomer is represented in a different shade of green, with monomer B (whose RBD transitions to the open state) depicted in dark green. Glycans were omitted from the figures to facilitate visualisation, but are present in all equilibrium and nonequilibrium simulations. **(B)** Detailed view of the upward movement of RBM_B_ and the contribution of the glycan linked to N234_C_ in the process. Glycans are shown with sticks, with the yellow sticks corresponding to the glycan bound to N234_C_. Please note that during the nonequilibrium trajectory, glycan N234_C_ slides into the pocket exposed beneath RBD_B_ following its upward movement, thereby stabilising this RBD in an “up” conformation.

In some of the D-NEMD trajectories of Omicron under acidic conditions, the upward movement of the RBD was closely coupled with the structural rearrangements of surrounding glycans. A particularly interesting example involves the glycan linked to N234, which exhibited a gradual conformational change as the RBD transitioned to the “up” position (Figure 6). The upward RBD movement exposed a previously buried pocket underneath it, into which glycan N234 gradually repositioned itself (see Figure 6 for an illustrative example of glycan N234 motions). By occupying this newly accessible cavity, this glycan helped stabilise the partially open conformation of the spike, effectively anchoring the RBD in the “up” position and reducing the likelihood of its reversion to the closed state. The structural role of the glycan N234 and its influence on RBD conformational dynamics were first suggested based on the analysis of conventional equilibrium MD simulations (56), with subsequent support from weighted ensemble simulations (85). The role of glycan N234 in spike activation and host receptor engagement was experimentally confirmed by biolayer interferometry experiments (81), with the substitution of asparagine N234 by alanine, which resulted in the ablation of the glycosylation site, showing to decrease binding to ACE2, underscoring the structural and functional importance of this glycan (56).

## Conclusions

In this study, we explored how pH shifts impact the functional dynamics and structural stability of the SARS-CoV-2 spike, focusing on the ancestral strain and two major variants of concern, Delta and Omicron BA.1. Using the D-NEMD approach, we characterised spike behaviour following a pH decrease, mimicking the environment of endosomal compartments, and a pH increase, reflecting the alkaline conditions accessible by nascent exhaled aerosol particles.

D-NEMD reveal that pH changes elicit distinct dynamical responses across the variants, with different motion profiles observed in key functional regions of the protein. Overall, the ancestral spike exhibits broad pH sensitivity (i.e. responds to both acidic and alkaline conditions), with acidic pH triggering conformational responses in the SD1, the upper part of the FPPR, and the S2 cleavage site, while alkaline pH affecting the SD2 and specific parts of the NTD. Notably, the RBM shows pH-dependent directional motions, with an outward rotation under acidic conditions and an inward motion at high pH. This detailed characterisation of the pH-responsive dynamics of the ancestral spike establishes a baseline and reference point for interpreting the evolutionary adaptations in the Delta and Omicron variants.

D-NEMD further reveals that Delta largely mirrors the ancestral spike’s response to acidic pH, with some subtle, region-specific behaviour differences present, highlighting the variant’s evolutionary adaptations. For example, Delta’s RBM displays enhanced motions compared to the ancestral protein, whereas the FPPR shows a contrasting trend, with attenuated responsiveness. At alkaline pH, Delta exhibits heightened sensitivity compared to the ancestral spike, with pronounced conformational changes in both the RBM and FPPR. Such enhanced responsiveness may trigger premature spike activation for fusion or induce structural changes that can lead to inactivation, thereby offering a possible mechanistic explanation for Delta’s experimentally observed reduced stability and infectivity in alkaline aerosols.

In contrast to the ancestral and Delta proteins, Omicron displays enhanced responsiveness to acidic conditions, with significant structural rearrangements observed in the RBD, FPPR and SD1. In Omicron, a reduction in pH triggers upward movements in these regions, resulting, in some cases, in spontaneous RBD opening. This marked change in Omicron’s functional dynamics aligns well with experimental evidence indicating this variant’s preference for endosomal entry into host cells, with the observed increased responsiveness reflecting its evolutionary adaptation to low-pH environments. Unlike earlier strains, Omicron was shown to rely less on TMPRSS2-mediated plasma membrane fusion, instead favouring receptor-mediated endocytosis *via* acidic endosomes. Additionally, Omicron shows attenuated responses to alkaline pH, with this reduced sensitivity suggesting enhanced structural resilience in high-pH environments, potentially contributing to its distinct infectivity profile and transmission advantage over earlier variants.

Our findings also highlight the crucial role of glycans in modulating spike function, with the glycan linked to N234 having an active role in the RBD transition from the closed to open state. Glycan N234 stabilises the RBD in its “up” conformation during the pH-induced transition. In Omicron, as the RBD shifts upward in response to acidic conditions, the glycan N234 slides underneath it, effectively anchoring it in place and preventing it from reverting to the “down” state.

To summarise, our findings, together, provide a detailed characterisation of the pH-dependent behaviour of spike variants, highlighting pH as a potential evolutionary pressure and contributing to a deeper understanding of viral adaptation mechanisms. They also emphasise the importance of incorporating both glycosylation and environmental pH variability in future structural and functional studies of viral envelope proteins, including those from coronaviruses, influenza, and HIV, to better understand viral adaptation, stability and transmission.

## Acknowledgements

This research was supported by the Biological and Biotechnological Sciences Research Council BBRC (BB/X009831/1 for ASFO; BB/W003449/1 for ASFO and AJM; BB/Z516533/1 for CS, IB, ADD, and AJM; BB/W00884X/1 for AEH and JPR). ASFO was supported at the University of Bristol by Oracle for Research. LT studentship at the University of Bristol is funded by the “Heather Corrie Impact Fund”. This work also received funding from the European Research Council (ERC) under the European Union’s Horizon 2020 research and innovation programme (Grant agreement No. <u>101021207</u>; project information: PREDACTED). REA acknowledges support from NSF RAPID MCB-2032054, an award from the RCSA Research Corp., and a UC San Diego Moore’s Cancer Center 2020 SARS-COV-2 seed grant. MD simulations were carried out using the Oracle Public Cloud Infrastructure (under an award to AJM and ASFO from Oracle for Research for COVID-19 research), the UK national supercomputer ARCHER2 and the GW4 Alliance Isambard 3 supercomputer. ASFO and AJM thank EPSRC via HECBIOSIM (EP/R029407/1 and EP/X035603/1) for providing ARCHER2 time.

## Data availability statement

All equilibrium and D-NEMD simulation data, including input files and trajectory files, will be made openly accessible via the University of Bristol data repository, data.bris, upon manuscript acceptance.

