## Supporting Information for "pH-induced structural changes in SARS-CoV-2 spike variants"

#### Materials and methods

##### Fully glycosylated ancestral, Delta, and Omicron BA.1 spike model construction

While a summary of model construction protocols will be described here for completeness, the models of ancestral, Delta, and Omicron BA.1 (hereafter referred to as Omicron) SARS-CoV-2 spike glycoprotein head domain (residues 13-1140) were constructed and fully glycosylated according to the protocols described in detail in the supporting information of Kim *et al.* (1). All models for the glycosylated spikes contain 15 disulfide bonds per trimer and are cleaved at the furin protease cleavage site located at S1/S2 interface. Similar models for the fully glycosylated "closed" spikes have now been widely tested and used in various applications (e.g. (1-7)). The simulation conditions and protocols used here were the same as those applied successfully in our previous work (8).

Ancestral spike: The model for the "closed" ancestral protein (i.e. with all receptor binding domains, RBDs, in the down position) was based on the cryo-EM structure PDB ID 6VXX (9), with fully resolved N-terminal domains (NTDs), RBDs, and pre-fusion loops grafted from the cryo-EM structure PDB ID 7JJI (10).

Delta spike: The model for the Delta "closed" spike was constructed using the ancestral "closed" structure as a baseline. Single point mutations were incorporated *via* the "mutate" command in VMD's psfgen (11). Delta's significantly remodelled NTD was incorporated by grafting the NTD from the partially resolved cryo-EM Delta spike structure PDB ID 7SO9 (12). Our Delta model includes several mutations and deletions relative to the ancestral protein, namely T19R, G142D, E156Δ-F157Δ, R158G, L452R, T478K, D614G, P681R, and D950N.

Omicron spike: The model for the Omicron "closed" trimer was built based on the experimental cryo-EM structure PDB ID 7TF8 (13). Missing loops in the furin cleavage site were grafted into our model from PDB ID 6VSB (14). The 7FT8 structure contains various unresolved loops, particularly within the NTDs; to address this, we incorporated a fully-resolved Omicron BA.1 NTD from the experimental structure PDB ID 7K4N

(15). Our Omicron model contains more than thirty mutations, deletions and insertions relative to the ancestral protein: A67V, H69Δ-V70Δ, T95I, G142D, V143Δ-Y144Δ-Y145Δ, N211Δ, L212I, EPE insertion in position 214, G339D, S371L, S373P, S375F, K417N, N440K, G446S, S477N, T478K, E484A, Q493R, G496S, Q498R, N501Y, Y505H, T547K, D614G, H655Y, N679K, P681H, N764K, D796Y, N856K, Q954H, N969K, and L981F.

Glycosylation/Protonation/Solvation/Neutralisation: All spike models were glycosylated following the same glycoprofile as in Casalino *et al.* (5), consistent with Watanabe et *al.* (16). Since the Schrödinger Protein Preparation Wizard (17) does not support *N*- or *O*-linked glycans, whereas standalone PROPKA (18, 19) can handle them, we used later to predict the p$K_a$ values for all titratable residues and the first to predict histidine tautomeric states (i.e., whether the proton is located on the Nδ or Nε of the imidazole ring in the histidine's side-chain). All titratable residues were predicted to adopt their standard protonation states at neutral/physiological pH. Specifically, glutamate and aspartate residues were assigned negative charges, lysine and arginine residues were positively charged, and histidines were predicted to be neutral (see Table S1). Each spike model was then solvated in explicit TIP3P water (20) boxes of 215 Å x 215 Å x 215 Å and neutralised with 150 mM NaCl.

**Table S1. Complete list of all titratable residues and their selected protonation states for each spike structure at pH = ~7, as predicted by PROPKA (18, 19).** "ASP" and "GLU" refer to the deprotonated (negatively charged) forms of aspartate and glutamate, respectively, while "LYS" and "ARG" denote the protonated (positively charged) forms of lysine and arginine. "HSD" corresponds to the singly protonated (on Nδ) neutral form of histidine, and "HSE" to the singly protonated (on Nε) one.

| pH | Spike | Potonation State | Residue Number |
|---|---|---|---|
| 7 | Ancestral | ASP | 40 53 80 88 111 138 178 198 215 228 253 287 290 294 364 389 398 405 420 427 428 442 467 568 571 574 578 586 614 627 663 737 745 775 796 808 820 830 839 843 848 867 936 950 979 985 994 1041 1084 1118 1127 1139 |
| | | GLU | 96 132 154 156 169 180 191 224 281 598 309 324 340 406 465 471 484 516 554 583 619 654 661 702 725 748 773 780 819 868 918 988 990 1017 1031 1072 1092 1111 |
| | | HSD | 146 207 245 519 1058 1088 |
| | | HSE | 49 66 69 625 655 1048 1064 1083 1101 |
| | | LYS | 41 77 97 113 129 147 150 182 187 195 202 206 278 300 304 310 356 378 386 417 424 444 458 462 528 529 535 537 557 558 733 776 786 790 795 811 814 825 835 854 921 933 947 964 986 1028 1038 1045 1073 1086 |
| | | ARG | 21 34 44 78 102 158 190 214 237 246 273 319 328 346 355 357 403 408 454 457 466 509 567 577 634 646 682 683 685 765 815 847 905 983 995 1000 1014 1019 1039 1091 1107 |
| 7 | Delta | ASP | 40 53 80 88 111 138 142 178 198 215 228 253 287 290 294 364 389 398 405 420 427 428 442 467 568 571 574 578 586 627 663 737 745 775 796 808 820 830 839 843 848 867 936 979 985 994 1041 1084 1118 1127 1139 |
| | | GLU | 96 132 154 169 180 191 224 281 298 309 324 340 406 465 471 484 516 554 583 619 654 661 702 725 748 773 780 819 868 918 988 990 1017 1031 1072 1092 1111 |
| | | HSD | 519 625 655 1058 1083 1088 |
| | | HSE | 49 66 69 146 207 245 1048 1064 1101 |

| | | | |
|---|---|---|---|
| | | LYS | 41 77 97 113 129 147 150 182 187 195 202 206 278 300 304 310 356 378 386 417 424 444 458 462 478 528 529 535 537 557 558 733 776 786 790 795 811 814 825 835 854 921 933 947 964 986 1028 1038 1045 1073 1086 |
| | | ARG | 19 21 34 44 78 102 190 214 237 246 273 319 328 346 355 357 403 408 452 454 457 466 509 567 577 634 646 681 682 683 685 765 815 847 905 983 995 1000 1014 1019 1039 1091 1107 |
| 7 | Omicron | ASP | 40 53 80 88 111 138 142 178 198 215 228 253 287 290 294 339 364 389 398 405 420 427 428 442 467 568 571 574 578 586 627 663 737 745 775 808 820 830 839 843 848 867 936 950 979 985 994 1041 1084 1118 1127 1139 |
| | | GLU | 96 132 154 156 169 180 191 2141 (inserted E) 2143 (inserted E) 224 281 298 309 324 340 406 465 471 516 554 583 619 654 661 702 725 748 773 780 819 868 918 988 990 1017 1031 1072 1092 1111 |
| | | HSD | 146 207 245 519 681 1058 1088 |
| | | HSE | 49 66 505 625 954 1048 1064 1083 1101 |
| | | LYS | 41 77 97 113 129 147 150 182 187 195 202 206 278 300 304 310 356 378 386 424 440 444 458 462 478 528 529 535 537 547 557 558 679 733 764 776 786 790 795 811 814 825 835 854 856 921 933 947 964 969 986 1028 1038 1045 1073 1086 |
| | | ARG | 21 34 44 78 102 158 190 214 237 246 273 319 328 346 355 357 403 408 454 457 466 493 498 509 567 577 634 646 682 683 685 765 815 847 905 983 995 1000 1014 1019 1039 1091 1107 |

<u>Initial system minimisation/Heating/Equilibration:</u> The above systems were energy minimised, heated, and equilibrated over 55.5 ns *via* equilibrium molecular dynamics (MD) simulations using NAMD2.14 (21). For each of the ancestral, Delta, and Omicron variants, three independent replicate simulations were conducted. Simulations were performed under both constant volume (NVT ensemble) and constant pressure (NpT ensemble) conditions (22), using TACC Frontera CPU resources.

The systems were parameterised using the CHARMM36m all-atom force field (23-25) for the protein, glycans, and ions, and the TIP3P water model (20). Standard MD simulation options were used, including: timestep = 2 fs; temperature = 310 K and pressure = 1 atm (which were maintained constant *via* the Nosé-Hoover thermostat (26) and Langevin barostat (27), respectively); bonds to hydrogen were constrained using the SHAKE algorithm (28); the long-range electrostatics were handled *via* Particle Mesh Ewald (29) (interpolation order = 4 and grid spacing = 2 Å), with a 10-12-13.5 Å cutoff scheme. See supporting information in Kim *et al.* (1) for complete details on the minimization, heating and equilibration simulations performed. Please note that, beyond the equilibration step, the methods relevant to this manuscript are no longer described in Kim *et al.* (1). The procedures and approaches that follow are specific to the present study and are, therefore, detailed in full below.

<u>Waterbox reduction and re-equilibration:</u> To reduce the total number of atoms in subsequent equilibrium MD simulations, while still taking care to ensure there were no glycans crossing periodic boundary conditions to interact with the spike's image, we moderately adjusted waterbox sizes from 215 Å x 215 Å x 215 Å to 195 Å x 215 Å x 205 Å, which was still sufficient to ensure at least 20 Å of solvent buffer around the largest width of the spike protein with glycans fully outstretched. This reduced the size of each system from ~970k atoms

(970899, 970613, and 970934 atoms for the ancestral, Delta, and Omicron variants, respectively) to ~840k atoms (839367, 839395, and 839438 atoms for the ancestral, Delta, and Omicron variants), corresponding to a savings of ~130k atoms per system. This optimisation allowed for a reduction of the systems' size while ensuring sufficient solvent buffering to prevent direct self/periodic image interactions during the simulations. A short (0.5 ns) re-equilibration simulation using the NpT ensemble was initiated for each replicate across the three systems following the reduction in system size. These simulations employed the same settings as the preceding equilibrium MD runs, as described above, including: timestep = 2 fs; temperature = 310 K and pressure = 1 atm (maintained constant *via* the Nosé-Hoover thermostat (26) and Langevin barostat (27), respectively); bonds to hydrogen were constrained using the SHAKE algorithm (28); the long-range electrostatics were handled *via* Particle Mesh Ewald (29) (interpolation order = 4 and grid spacing = 2 Å), with a 10-12-13.5 Å cutoff scheme. The final frames and corresponding topologies were then converted from NAMD- to GROMACS-compatible file formats using the CHARMM-GUI webserver and the Force Field Converter tool (30-32).

**Long equilibrium molecular dynamics (MD) simulations**

Subsequently, long equilibrium simulations of 750 ns each were conducted for the three replicates of the ancestral, Delta and Omicron systems using GROMACS (33), with the same settings and conditions as in our previous work (5, 8). All long equilibrium simulations were performed on the Oracle Cloud Infrastructure (OCI), the UK national supercomputer ARCHER2 (www.archer2.ac.uk/) and the GW4 Isambard 3 high-performance computing machine.

**Dynamical-nonequilibrium MD (D-NEMD) simulations**

Over 1150 nonequilibrium simulations were performed to investigate pH-induced structural changes in the glycosylated ancestral, Delta and Omicron spikes. For each system, two sets of nonequilibrium simulations were conducted, one simulating a pH decrease and the other a pH increase (Figure S8). A total of 192 nonequilibrium simulations (i.e. 64 simulations per replicate), each 20 ns in duration, were run for each spike system-pH combination, resulting in 1152 simulations and an aggregate time exceeding >23 µs.

All starting conformations for D-NEMD were extracted from the equilibrated part of the long equilibrium trajectories at physiological pH (Figure S8). The conformations were taken every ten nanoseconds, and, in each one of them, the protonation state of specific residues was (instantaneously) changed to mimic either a decrease or an increase in pH. The predicted p$K_a$ value of titratable residues, calculated using PROPKA (18, 19), were used to identify which residues should undergo protonation state changes to mimic the desired pH shift. The following adjustments were made for each monomer in the respective spike systems:

<u>Ancestral spike</u>: to simulate a pH decrease, the protonation states of H66, H146, H519, D568, D574, H625, H655, D737, E773, E819, E918, H1064, H1083, E1092, H1101, and E1111 were modified. For a pH increase, the states of K41, K97, K129, K195, K278, K300, K304, K417, K424, K733, K921, K964, K1028, and K1038 were adjusted.

<u>Delta spike</u>: residues adjusted to simulate a pH decrease included H66, H69, D88, H146, E132, E191, H245, E298, D568, D574, H625, H655, D737, D745, E780, E819, E918, E988, E990, D1041, H1064, H1083, H1101 and E1111. For a pH increase, the protonation states of K41, K77, K97, K195, K300, K304, K386, K417, K424, K733, K854, K964, K1028 and K1038 were modified.

<u>Omicron spike</u>: the protonation of H66, E96, E132, E154, E191, D198, H207, H245, E224, D294, E298, E309, D364, D389, D398, D405, E406, E465, E516, H519, D578, D586, E619, H625, D627, H681, D737, D745, E748, E773, E819, D843, D867, E918, H954, D979, D985, E988, E990, D994, D1041, H1064, E1072, H1083, E1092, H1101 and E1111 were adjusted to simulate a pH decrease. For a pH increase, the states of K41, K77, K97, K195, K300, K304, K386, K444, K458, K462, K528, K547, K558, K764, K776, K786, K795, K854, K856, K947, K964, K969, K1028, and K1038 were modified.

At low pH, the specified histidine residues were modelled as positively charged, while aspartate and glutamate residues were treated as neutral. At high pH, lysine residues were considered neutral. Arginine residues, due to their exceptionally high p$K_a$ values (approximately 13), were not expected to undergo protonation changes within the pH range examined in this study and were therefore left unchanged in the nonequilibrium simulations. The resulting nonequilibrium systems, representing either a pH decrease or increase, were then simulated for 20 ns using GROMACS (33), employing the same settings and conditions as the long equilibrium simulations described above (Figure S8).

The perturbation applied, namely altering the protonation states of specific residues, was designed to drive the system out of equilibrium, thereby inducing structural and dynamic changes (34, 35). The spikes' response to pH shifts were extracted using the Kubo-Onsager relation (34-38). For each time point, the displacement vector of every $C_\alpha$ atom was calculated by comparing its position in equilibrium and nonequilibrium simulations (Figure S8B). These displacement vectors were then averaged across 192 simulations to obtain an average vector for each $C_\alpha$ atom. These average vectors indicate the direction of movement in response to the pH shift, while their norm represent the amplitude of the responses (Figures S9-S14). The convergence and statistical significance of the responses were assessed by determining the standard error of the mean (Figures S9-S14), as previously described (34, 35). Average displacement vectors were considered meaningful only if their magnitudes remained greater than zero after subtracting the standard error of the mean; vectors not meeting this criterion were excluded from further analysis. The average displacement vectors were visualised on the energy-minimised structures of the ancestral, Delta, and Omicron spikes using the Modevectors plugin (39) for PyMOL (40).

In addition to the nonequilibrium simulations, an extra set of 576 "null perturbation" simulations (i.e. 192 trajectories per system), each lasting 20 ns, was also conducted to further test the significance of the D-NEMD responses and ensure they do not stem from the proteins' natural equilibrium motions. These simulations used the same starting conformations as the D-NEMD runs but introduced no external perturbation, keeping the systems in an equilibrium state. As a result, any responses observed in the "null perturbation" simulations reflect only the inherent equilibrium dynamics of the proteins. This null perturbation test, originally described by Kamsri *et al*. (41), involved randomising atomic velocities according to the Boltzmann distribution while maintaining the original systems (in our case, retaining the original protonation state of all residues). The analysis of these simulations followed the same protocol as the nonequilibrium simulations above, applying the Kubo-Onsager relation (34, 35) to extract the evolution of pairwise $C_\alpha$ displacements over time. These responses allow us to distinguish intrinsic equilibrium dynamics from those responses triggered by the external perturbations, thereby facilitating the identification of the functionally relevant motions and enhancing data interpretation. All nonequilibrium and "null perturbation" simulations were performed on the Oracle Cloud Infrastructure (OCI), the UK national supercomputer ARCHER2 and the GW4 Isambard 3 high-performance computing machine.

### Supplementary Figures

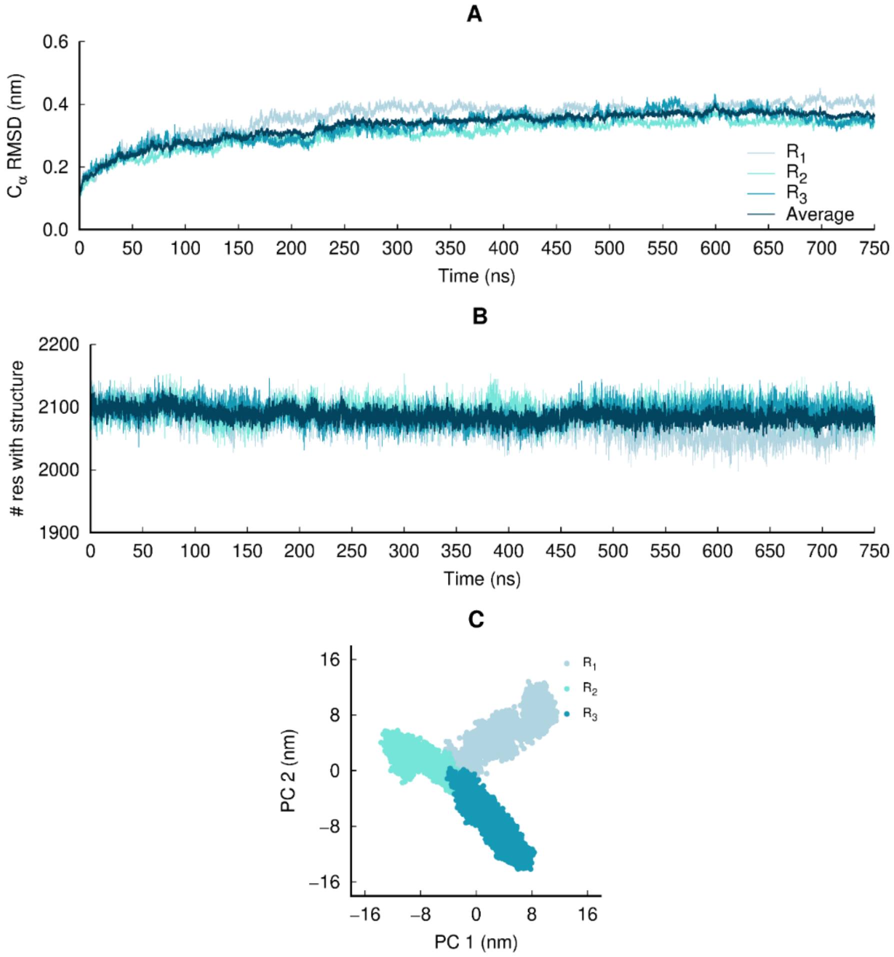


**Figure S1. Structural stability, equilibration and sampling of the equilibrium simulations of the glycosylated SARS-CoV-2 ancestral spike**. **(A)** Temporal evolution of the $C_\alpha$ RMSD relative to the starting structure. The dark blue line corresponds to the average $C_\alpha$ RMSD across all three replicates. These plots indicate that the simulations were stable over the simulation time. **(B)** Time evolution of the spike's secondary structure content (assigned by DSSP (42)), showing the number of residues assigned in total to α-helix, $3_{10}$- helix, 5-helix, β-sheet and β-bridge secondary structure classes. The dark blue line corresponds to the average number of residues with defined secondary structure across all three replicates. **(C)** Principal component analysis (PCA) for all $C_\alpha$ atoms of the three replicates. All replicates were combined for the PCA, with one conformation per 100 ps per replicate (in a total of 22501 frames).

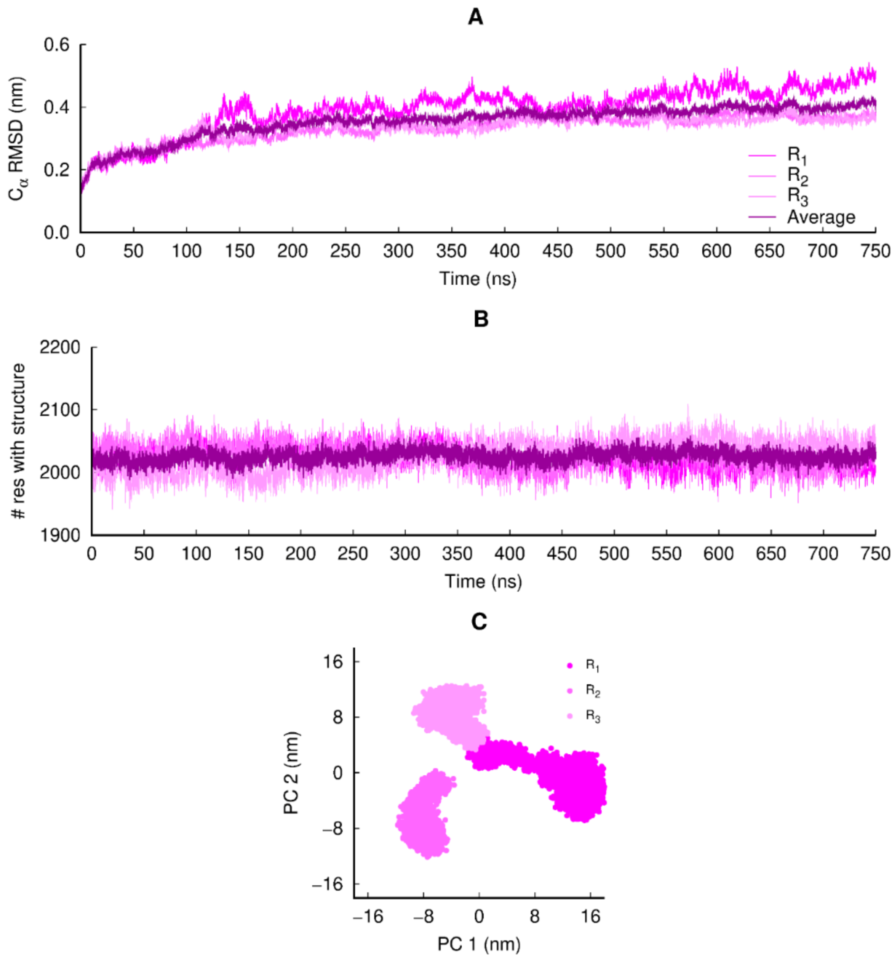


**Figure S2. Structural stability, equilibration and sampling of the equilibrium simulations of the glycosylated SARS-CoV-2 Delta spike**. **(A)** Temporal evolution of the $C_\alpha$ RMSD relative to the starting structure. The dark pink line corresponds to the average $C_\alpha$ RMSD across all three replicates. **(B)** Time evolution of the spike's secondary structure content. The dark pink line corresponds to the average number of residues with defined secondary structure across all three replicates. **(C)** PCA for all $C_\alpha$ atoms of the three replicates. For more details, see the legend of Figure S1.

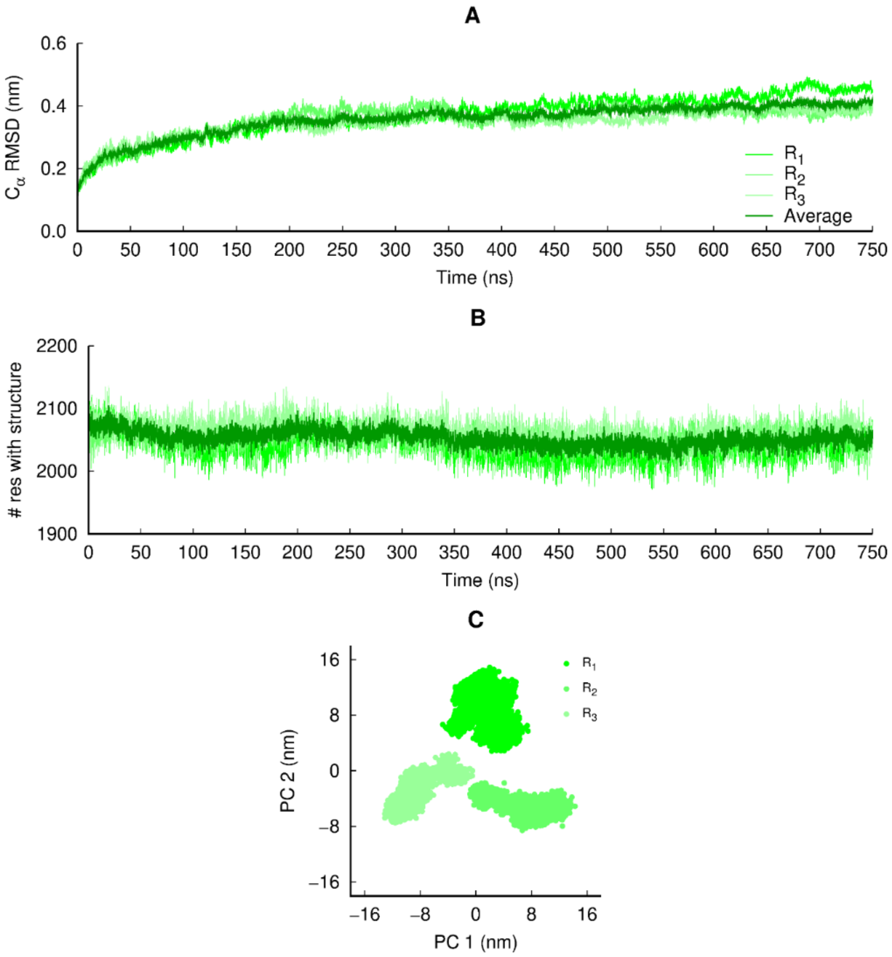


**Figure S3. Structural stability, equilibration and sampling of the equilibrium simulations of the glycosylated SARS-CoV-2 Omicron spike**. **(A)** Temporal evolution of the $C_\alpha$ RMSD relative to the starting structure. The dark green line corresponds to the average $C_\alpha$ RMSD across all three replicates. These plots indicate that the systems are stable over the simulation time. **(B)** Time evolution of the spike's secondary structure content. The dark green line corresponds to the average number of residues with defined secondary structure across all three replicates. **(C)** PCA for all $C_\alpha$ atoms of the three replicates. For more details, see the legend of Figure S1.

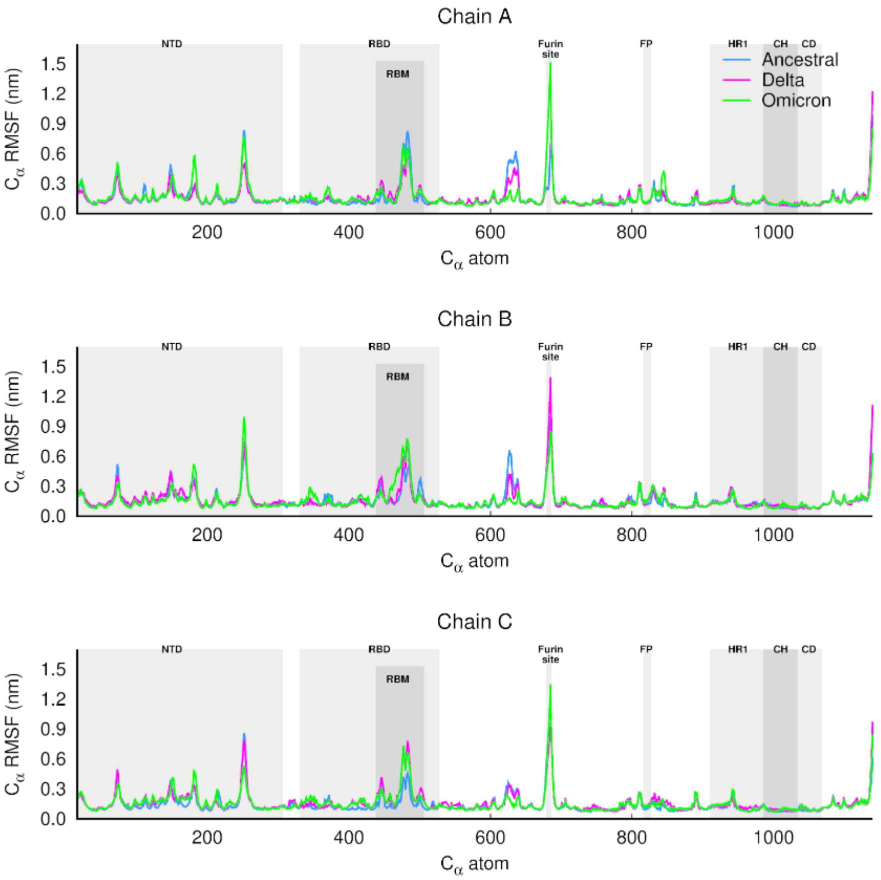


**Figure S4. Average Cα root mean square fluctuations (RMSF) for chains A, B and C of the ancestral (blue line), Delta (pink line) and Omicron (green line) spike**. The Cα RMSF was calculated using the equilibrated part of the trajectory (from 100-750 ns) and averaged across the three replicates for each system. The positions of key structural motifs are highlighted in grey, namely the N-terminal domain (NTD), receptor-binding domain (RBD), receptor-binding motif (RBM), fusion peptide (FP), heptad repeat 1 (HR1), central helix (CH), and connector domain (CD). Please zoom in on the image for detailed visualisation.

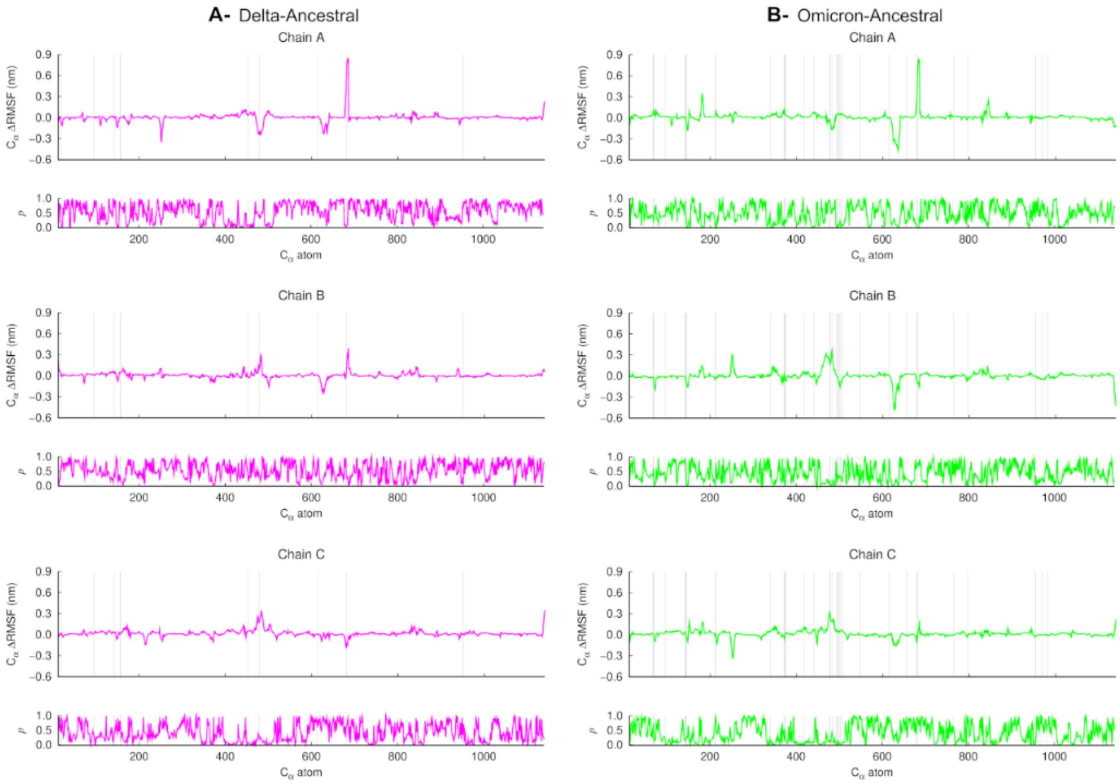


**Figure S5. Average Cα RMSF differences between the Delta (A) and Omicron (B) variants compared to the ancestral spike, along with associated *p*-values.** The statistical significance of the differences was assessed using a Student's *t*-test. Vertical grey lines highlight the position of substitutions, insertions and deletions present in each variant. Please zoom in on the image for detailed visualisation.

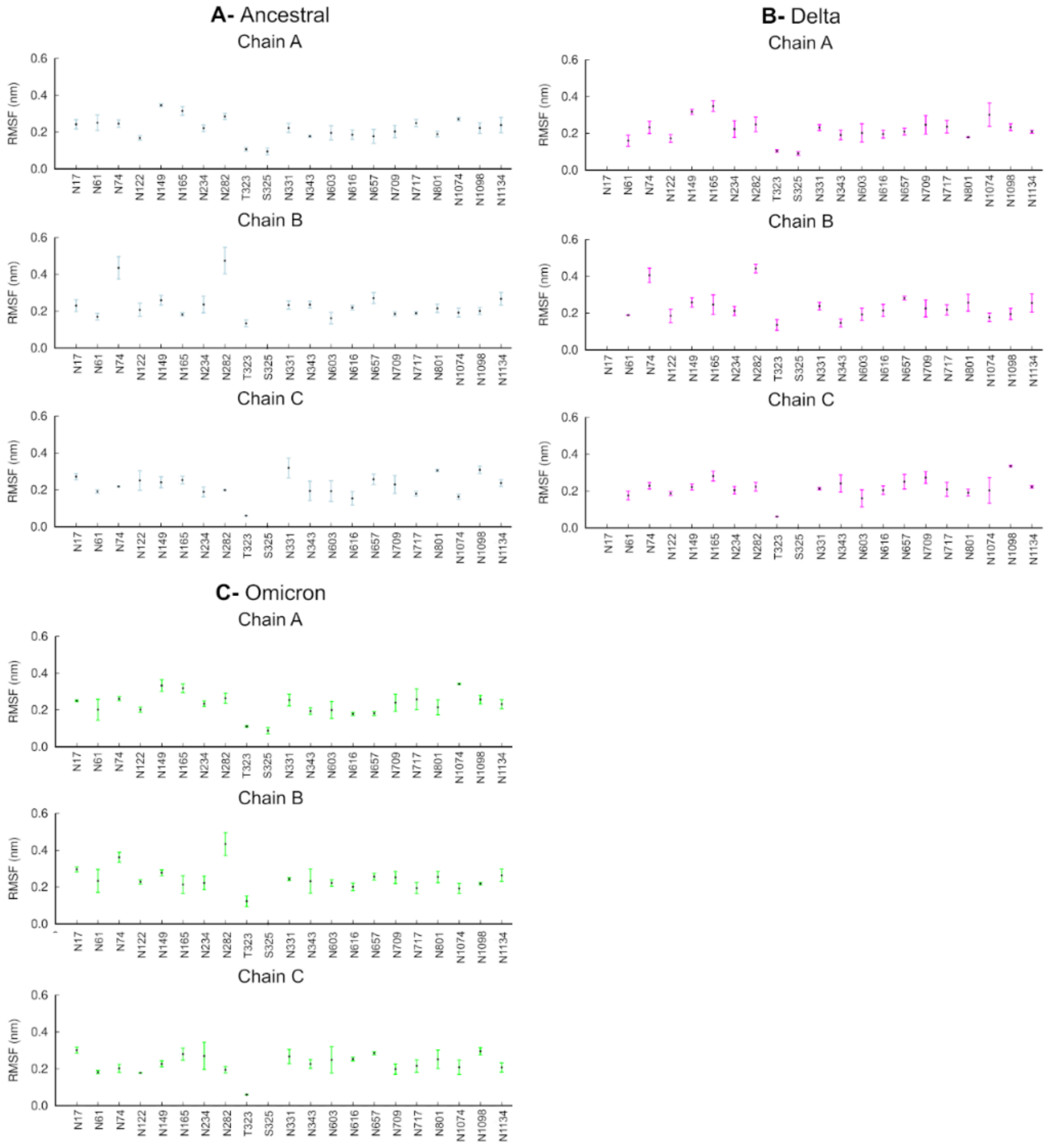


**Figure S6. Average RMSF for each glycan in chain A, B, and C of the ancestral (A), Delta (B) and Omicron (C) spike.** The RMSF was calculated using the equilibrated part of the trajectories (from 100-750 ns) and averaged across all three replicates for each complex. The vertical lines represent the standard deviation. Please zoom in on the image for detailed visualisation.

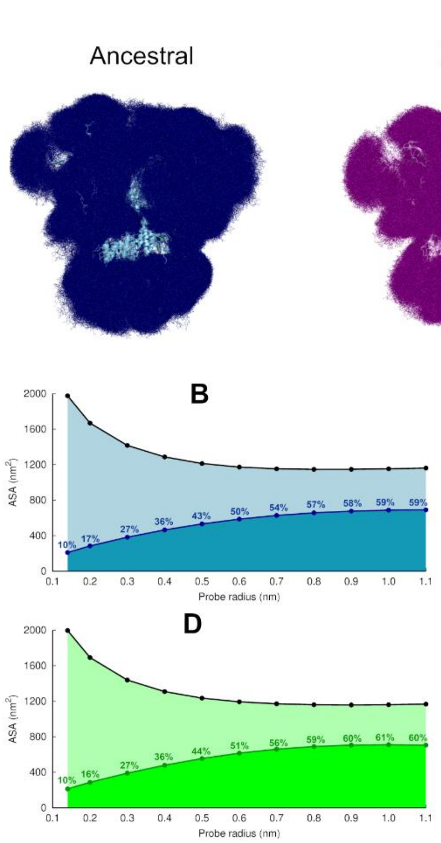


**Figure S7. Glycan shield in the ancestral, Delta and Omicron spike**. **(A)** Overlapping of the glycans' conformations in all three replicas (in a total of 1128 frames) for the ancestral, Delta and Omicron systems. The protein is shown as a cartoon, whereas the glycans are represented with sticks. **(B-C)** Solvent accessible surface area of the protein and the area shielded by glycans at multiple probe radii for the ancestral, Delta and Omicron spikes. The probe radius ranges from 0.14 nm (corresponding to a water molecule) to 1.1 nm (corresponding to a small antibody molecule). The values were averaged across all three replicates. The area shielded by the glycans corresponds to the dark blue, purple and dark green lines, whereas the black lines represent the accessible surface area of the protein without glycans (similarly to e.g. ref (5, 8, 43)).

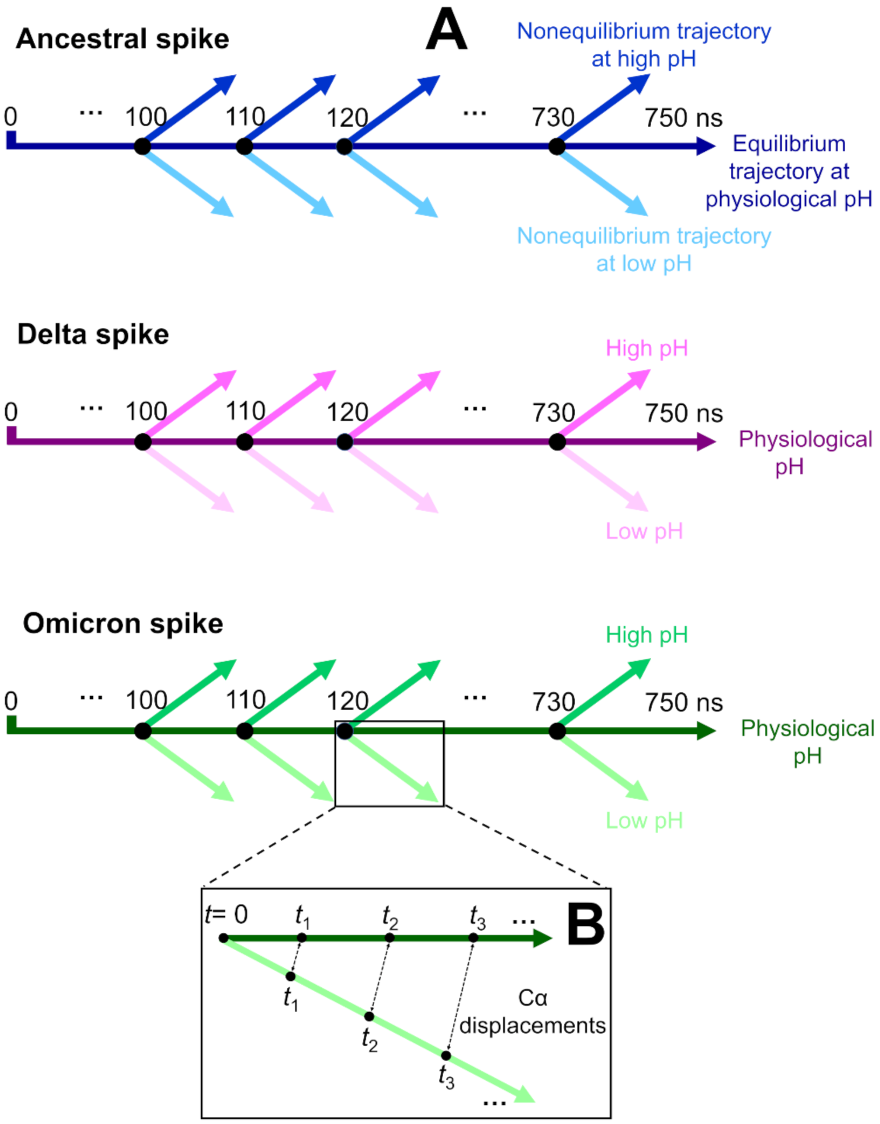


**Figure S8. Schematic overview of the D-NEMD approach and analysis**. **(A)** Three equilibrium MD simulations, 750 ns each, were performed for the glycosylated, cleaved (with cleavage at the S1/S2 interface) ancestral, Delta and Omicron spikes in the closed state at physiological pH. From the equilibrated portion of each trajectory (from 100–750 ns), frames were extracted every 10 ns to serve as starting points for short nonequilibrium simulations under low and high pH conditions. Perturbations were introduced by modifying the protonation states of specific residues to mimic pH shifts, and each nonequilibrium simulation was run for 20 ns. **(B)** The Kubo-Onsager relation (34, 35) was used to extract the response of the system to pH change. For that, for each pair of equilibrium (at physiological pH) and nonequilibrium (at low/high pH) trajectories, $C_\alpha$ displacement vectors were calculated at selected time points (0, 0.1, 1, 10, and 20 ns) and averaged across 192 simulations to capture the collective response.

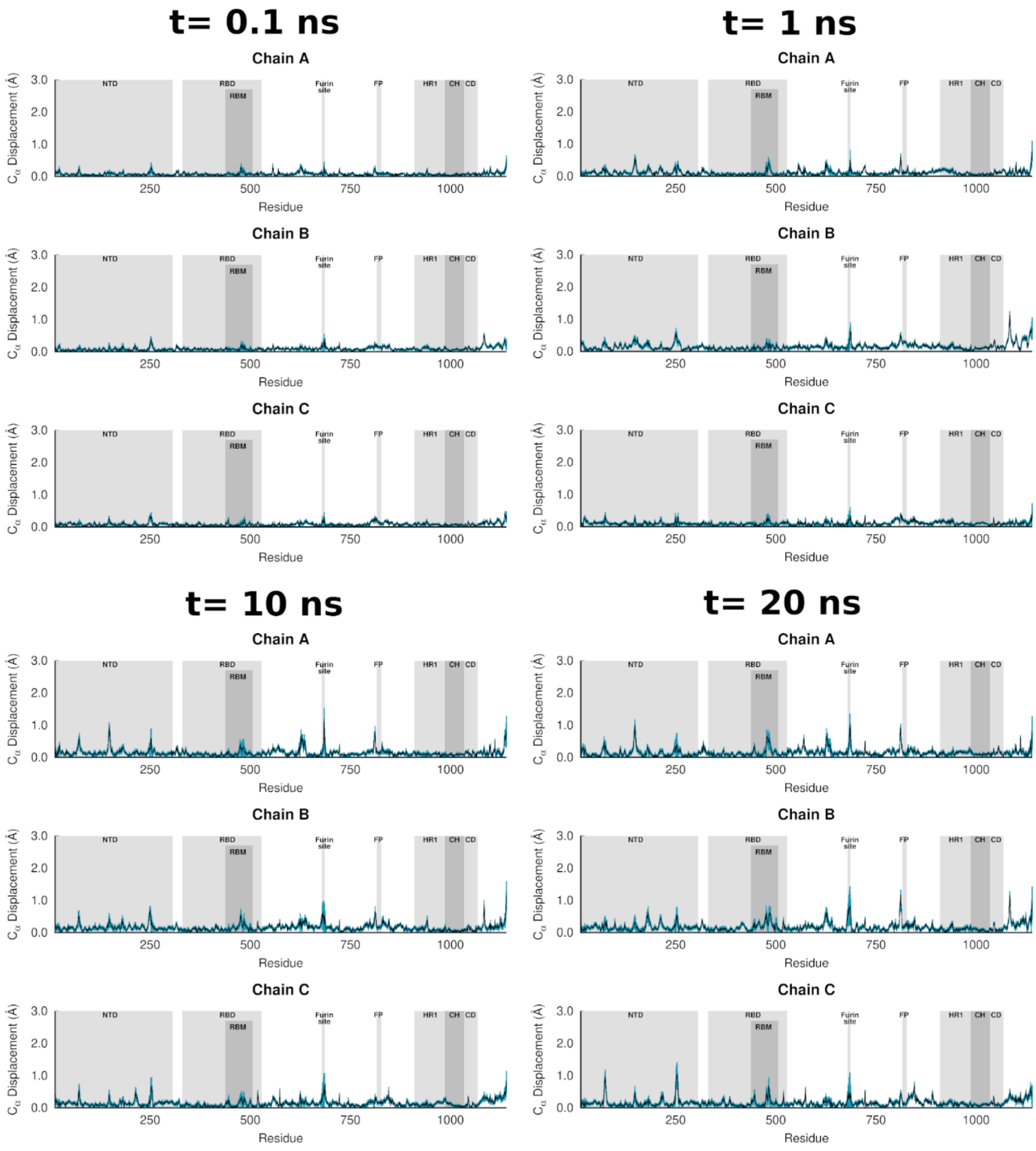


**Figure S9. Average $C_\alpha$-positional displacement and corresponding standard errors at *t* = 0.1, 1, 10 and 20 ns following a pH decrease for the ancestral spike**. The average displacements were determined using the Kubo-Onsager relation (34, 35) by comparing the nonequilibrium (at low pH) and equilibrium (at physiological pH) simulations at equivalent time points. The averages were calculated across the 192 pairs of simulations. The values plotted correspond to the average $C_\alpha$ responses after removing the intrinsic protein fluctuations *via* the "null perturbation" analysis. The vertical blue lines represent the standard error of the mean. The positions of key structural motifs are highlighted in grey, namely the NTD, RBD, receptor-binding motif (RBM), fusion peptide (FP), heptad repeat 1 (HR1), central helix (CH), and connector domain (CD). Please zoom in on the image for detailed visualisation.

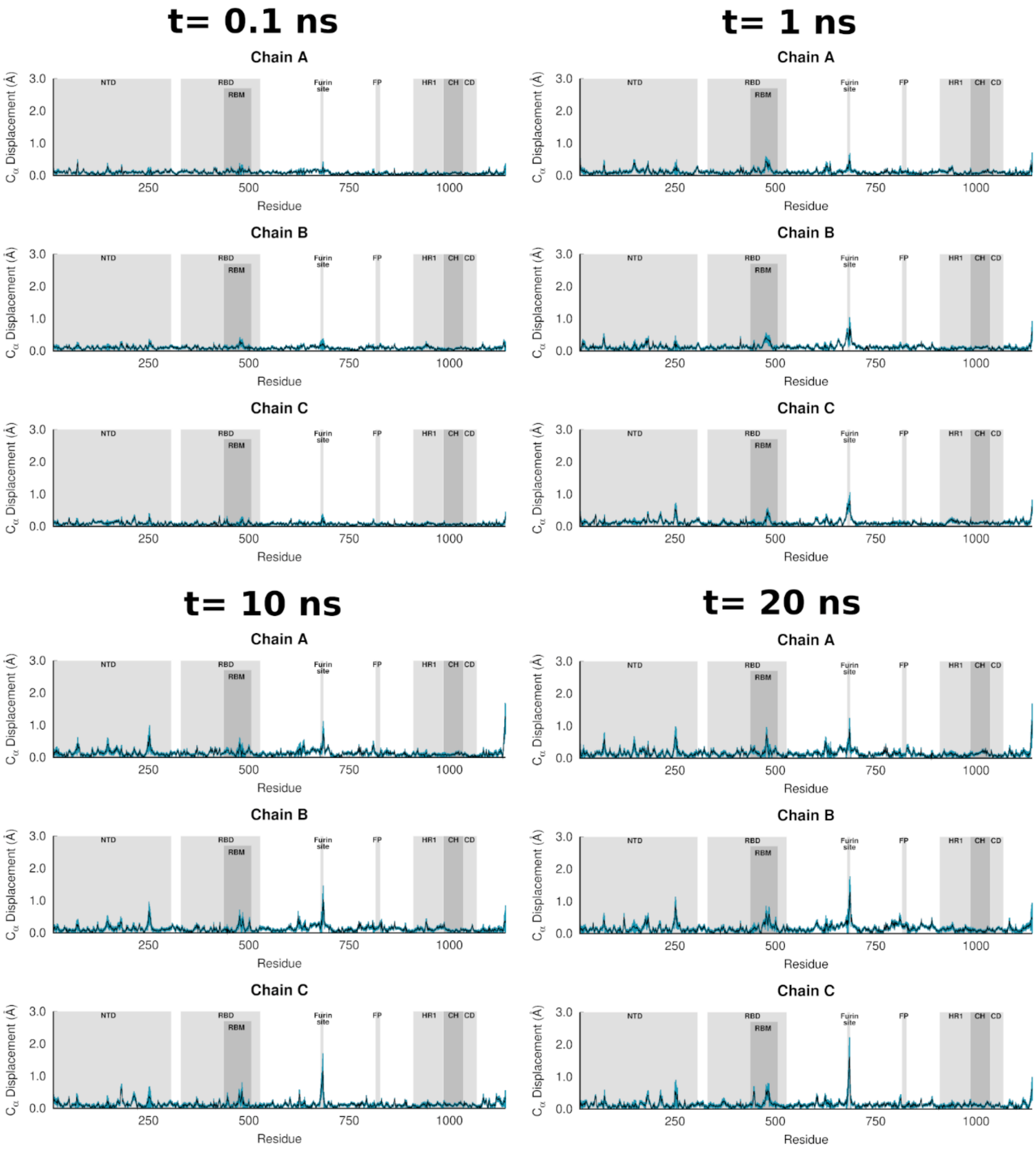


**Figure S10. Average Cα-positional displacement and corresponding standard errors at *t* = 0.1, 1, 10 and 20 ns following a pH increase for the ancestral spike**. The average displacements were determined using the Kubo-Onsager relation (34, 35) by comparing the nonequilibrium (at high pH) and equilibrium (at physiological pH) simulations at equivalent time points. For more details, see the caption of Figure S9. Please zoom in on the image for detailed visualisation.

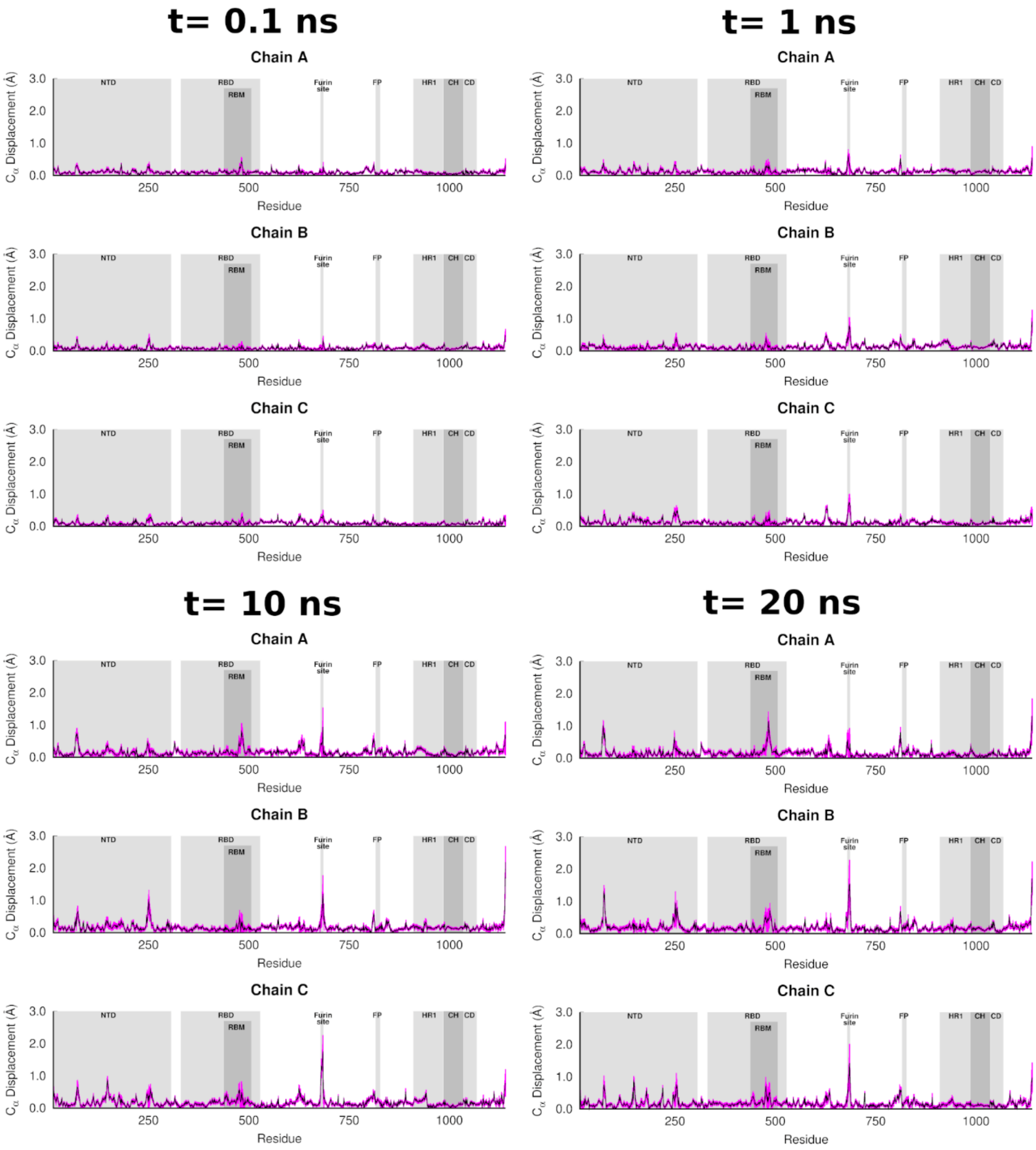


**Figure S11. Average $C_\alpha$-positional displacement and corresponding standard errors at *t* = 0.1, 1, 10 and 20 ns following a pH decrease for the Delta spike**. The average displacements were determined using the Kubo-Onsager relation (35, 36) by comparing the nonequilibrium (at low pH) and equilibrium (at physiological pH) Delta simulations at equivalent time points. The vertical pink lines represent the standard error of the mean. For more details, see the caption of Figure S9. Please zoom in on the image for detailed visualisation.

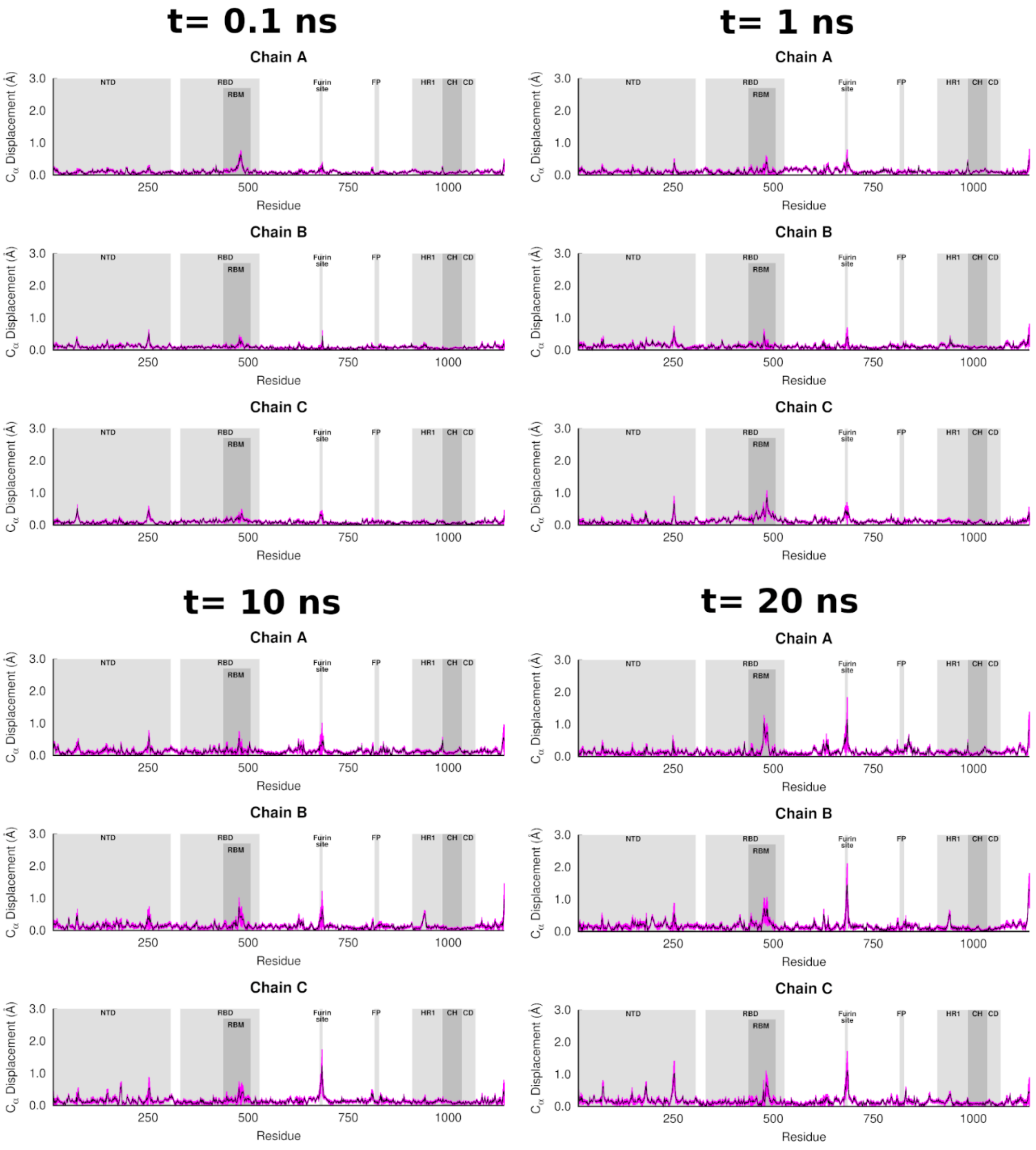


**Figure S12. Average $C_{\alpha}$-positional displacement and corresponding standard errors at *t* = 0.1, 1, 10 and 20 ns following a pH increase for the Delta spike**. The average displacements were determined using the Kubo-Onsager relation (35, 36) by comparing the nonequilibrium (at high pH) and equilibrium (at physiological pH) Delta simulations at equivalent time points. The vertical pink lines represent the standard error of the mean. For more details, see the caption of Figure S9. Please zoom in on the image for detailed visualisation.

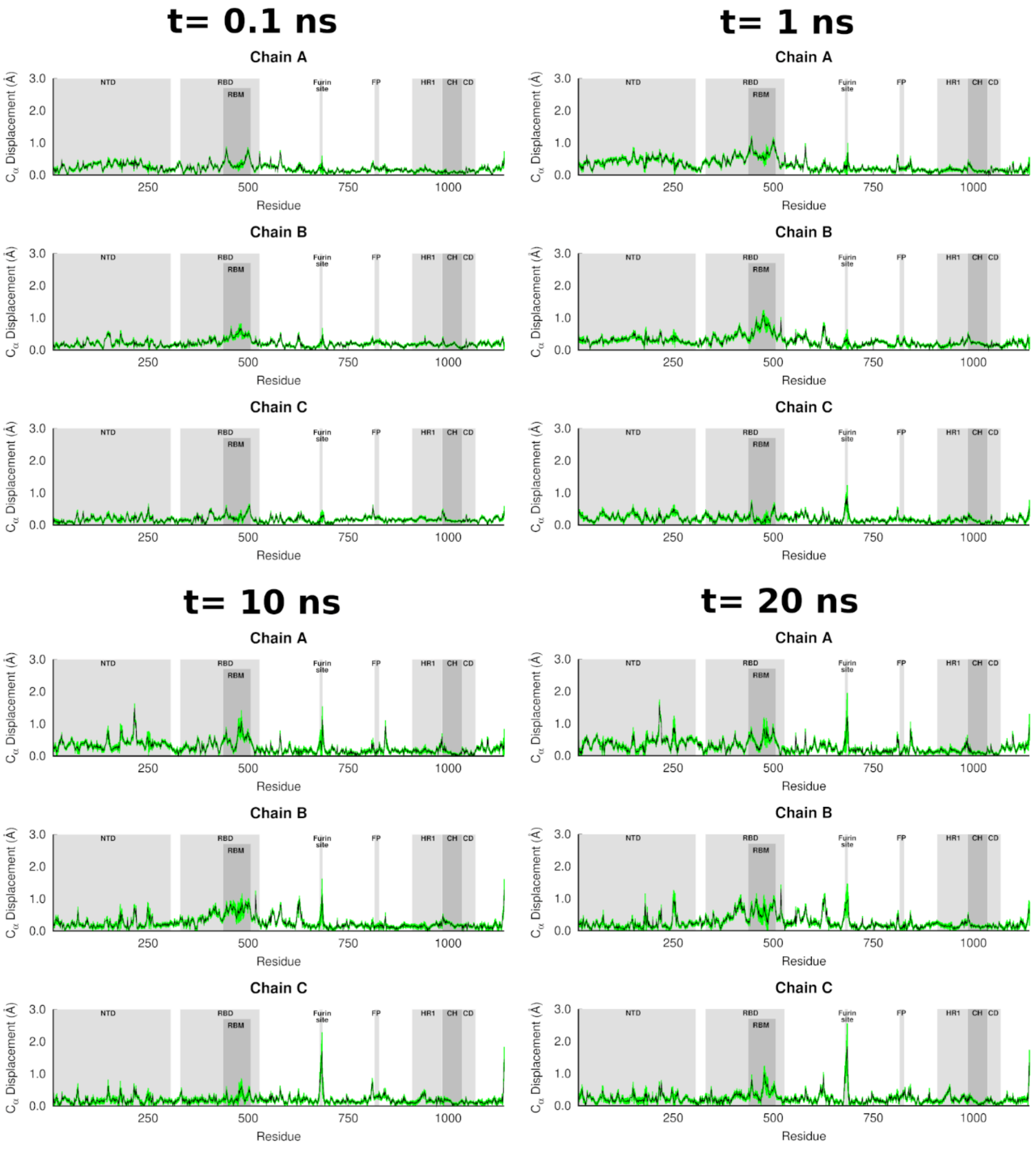


**Figure S13. Average $C_\alpha$-positional displacement and corresponding standard errors at *t* = 0.1, 1, 10 and 20 ns following a pH decrease for the Omicron spike**. The average displacements were determined using the Kubo-Onsager relation (35, 36) by comparing the nonequilibrium (at low pH) and equilibrium (at physiological pH) Omicron simulations at equivalent time points. The vertical green lines represent the standard error of the mean. For more details, see the legend of Figure S9. Please zoom in on the image for detailed visualisation.

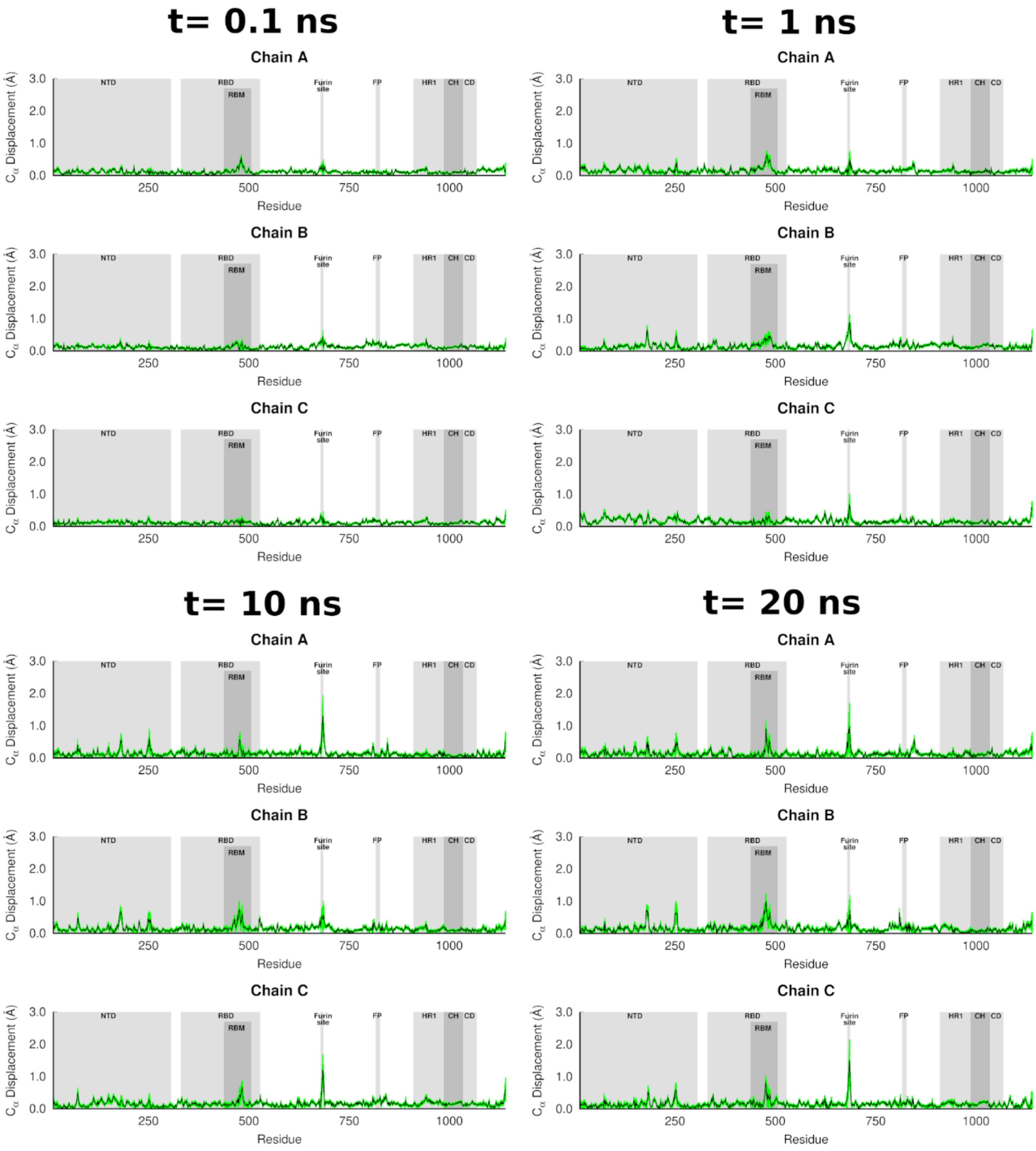


**Figure S14. Average Cα-positional displacement and corresponding standard errors at *t* = 0.1, 1, 10 and 20 ns following a pH increase for the Omicron spike**. The average displacements were determined using the Kubo-Onsager relation (35, 36) by comparing the nonequilibrium (at high pH) and equilibrium (at physiological pH) Omicron simulations at equivalent time points. The vertical green lines represent the standard error of the mean. For more details, see the caption of Figure S9. Please zoom in on the image for detailed visualisation.

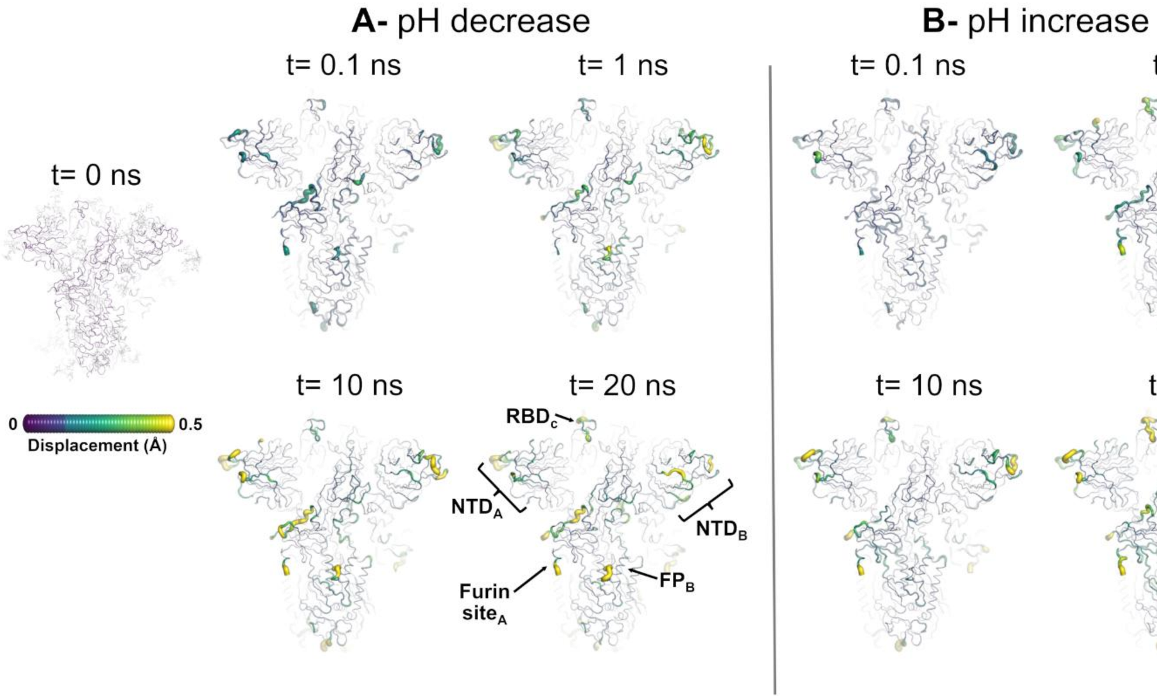


**Figure S15. Structural response of the ancestral spike to pH changes from the viewpoint of the NTDA, NTDB, FPB, and RBDC.** The average $C_\alpha$ displacements at $t$ = 0, 0.1, 1, 10, and 20 ns following a pH decrease **(A)** and increase **(B)** are shown. The displacements are mapped onto the starting structure for the equilibrium simulations of the ancestral protein at physiological pH. The magnitude of each residue's response was calculated using the Kubo-Onsager relation (34, 35) and corresponds to the norm of the average $C_\alpha$ displacement vector between every pair of equilibrium and nonequilibrium trajectories. The final displacements correspond to the average D-NEMD responses after removing the intrinsic protein fluctuations via the "null perturbation" analysis. Cartoon thickness and colours (scale shown on the left) indicate the extent of the average $C_\alpha$-positional displacements. Glycans are depicted as light grey sticks in the leftmost image but were omitted from the remaining panels (showing the responses at $t$=0.1, 1, 10 and 20 ns) to facilitate visualisation. Each RBD, NTD, furin site and FP are subscripted with their chain ID (A, B or C).

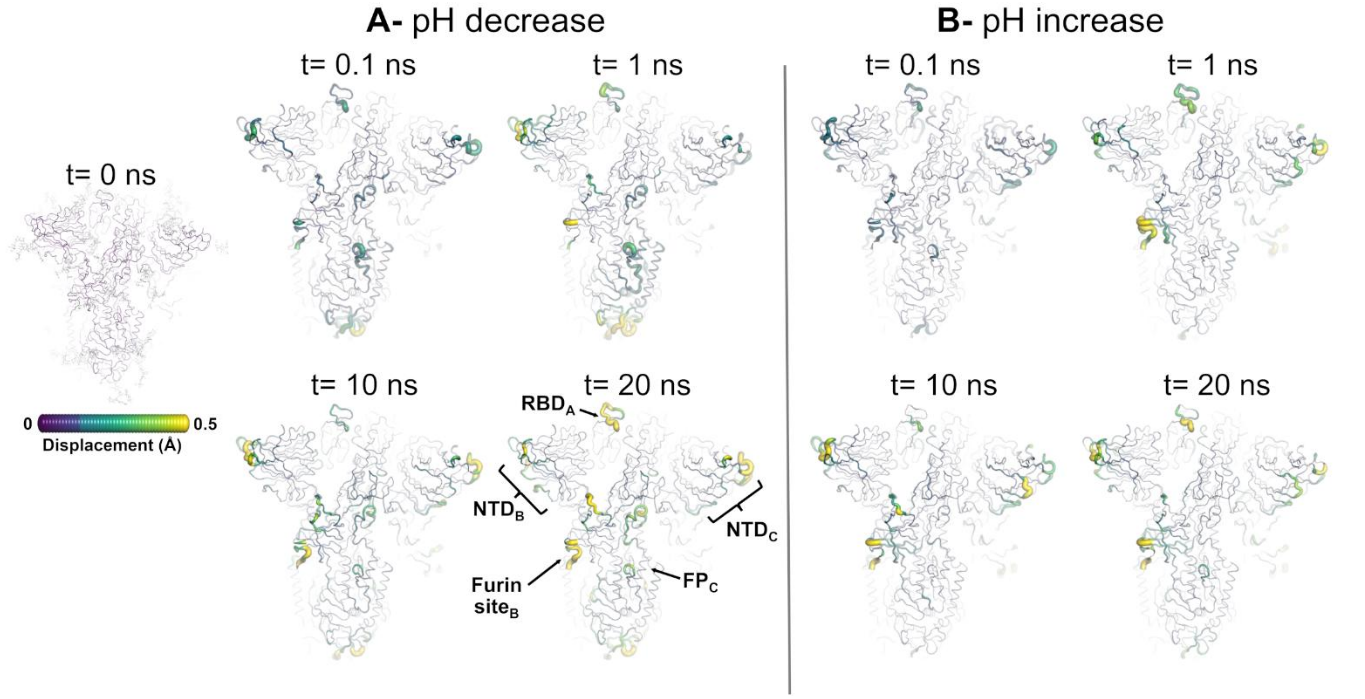


**Figure S16. Structural response of the ancestral spike to pH changes from the viewpoint of the RBD$_A$, NTD$_B$, NTD$_C$, and FP$_C$**. The average C$_\alpha$ displacements at $t$ = 0, 0.1, 1, 10 and 20 ns following a pH decrease **(A)** and increase **(B)** are shown. For more details, see the caption of Figure S15.

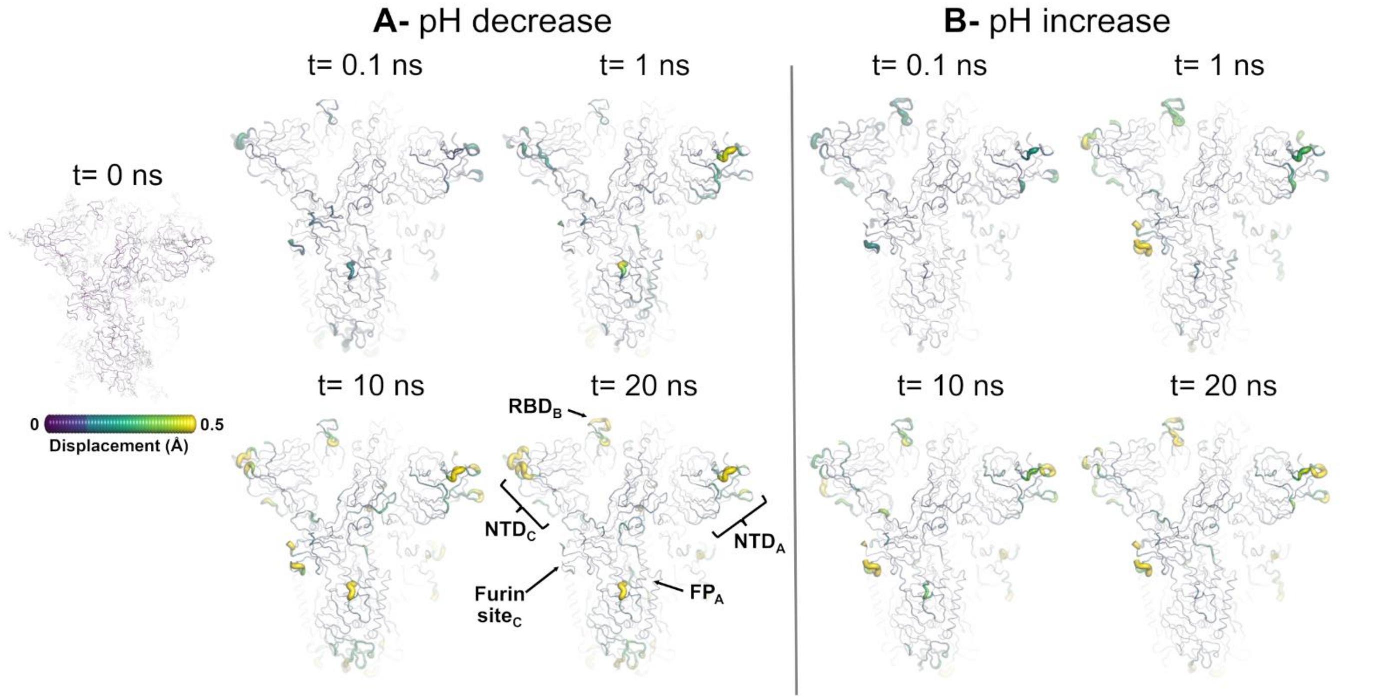


**Figure S17. Structural response of the ancestral spike to pH changes from the viewpoint of the NTD$_A$, FP$_A$, RBD$_B$, and NTD$_C$.** The average C$_\alpha$ displacements at $t$ = 0, 0.1, 1, 10 and 20 ns following a pH decrease **(A)** and increase **(B)** are shown. For more details, see the caption of Figure S15.

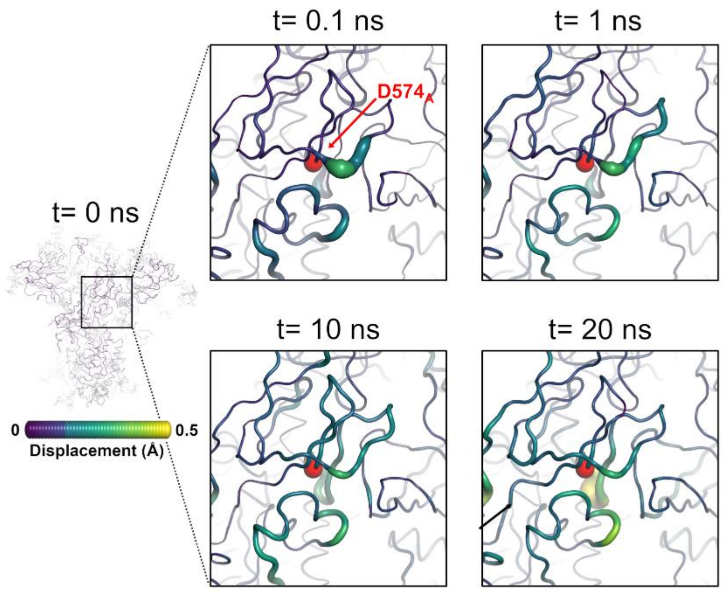


**Figure S18. Structural response of the ancestral spike to pH changes from the viewpoint of D574$_A$ and FPPR$_B$**. The average $C_\alpha$ displacements at $t$ = 0, 0.1, 1, 10 and 20 ns following a pH decrease **(A)** and increase **(B)** are shown. The displacements are mapped onto the starting structure for the equilibrium simulations of the ancestral protein at physiological pH. The magnitude of each residue's response was calculated using the Kubo-Onsager relation (34, 35) and corresponds to the norm of the average $C_\alpha$ displacement vector between every pair of equilibrium and nonequilibrium trajectories. The final displacements correspond to the average D-NEMD responses after removing the intrinsic protein fluctuations via the "null perturbation" analysis. Cartoon thickness and colours (scale shown on the left) indicate the extent of the average $C_\alpha$-positional displacements. Glycans are depicted as light grey sticks in the leftmost image but were omitted from the remaining panels (showing the responses at $t$=0.1, 1, 10 and 20 ns) to facilitate visualisation.

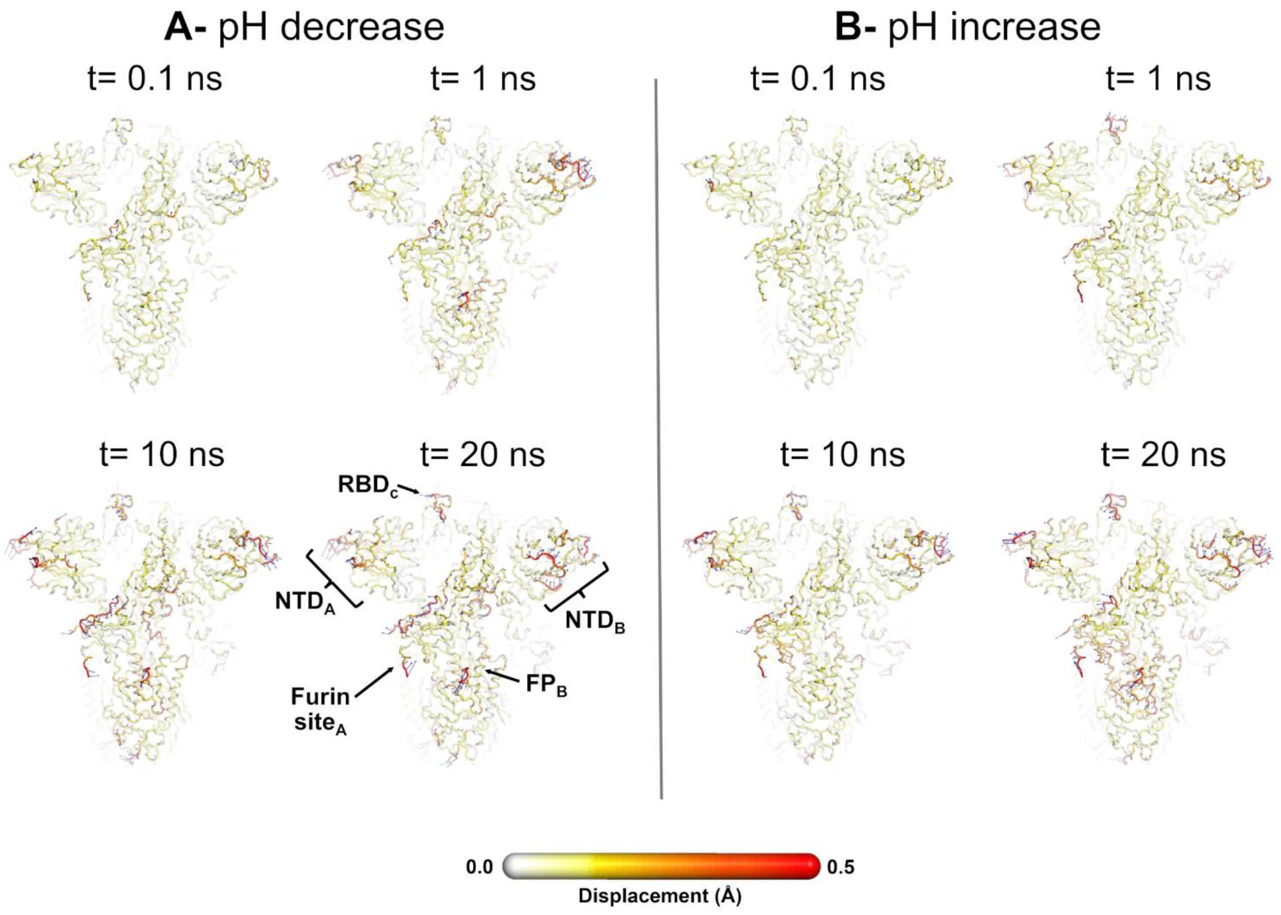


**Figure S19. Average displacement vectors from D-NEMD in response to a pH decrease (A) and increase (B) from the viewpoint of the $NTD_A$, $NTD_B$, $FP_B$, and $RBD_C$ for the ancestral spike.** This figure shows a full view of the protein, with average displacement vectors at $t$ = 0.1, 1, 10, and 20 ns following a pH change. The blue arrows correspond to the average $C_\alpha$ displacement vectors (calculated between the equilibrium and nonequilibrium trajectories averaged across all 192 pairs of trajectories) after removing the intrinsic protein fluctuations via the "null perturbation" analysis. Vectors with length ≥0.08 Å are displayed as blue arrows with a scale-up factor of 10. Average $C_\alpha$ displacements (i.e. the norm of the average $C_\alpha$ displacement vectors) are represented on a white-yellow-orange-red scale. Glycans were omitted from the figures to facilitate visualisation, but were present in the simulations. Each RBD, NTD, furin site and FP are subscripted with their chain ID (A, B or C). Please zoom in on the image for a detailed visualisation.

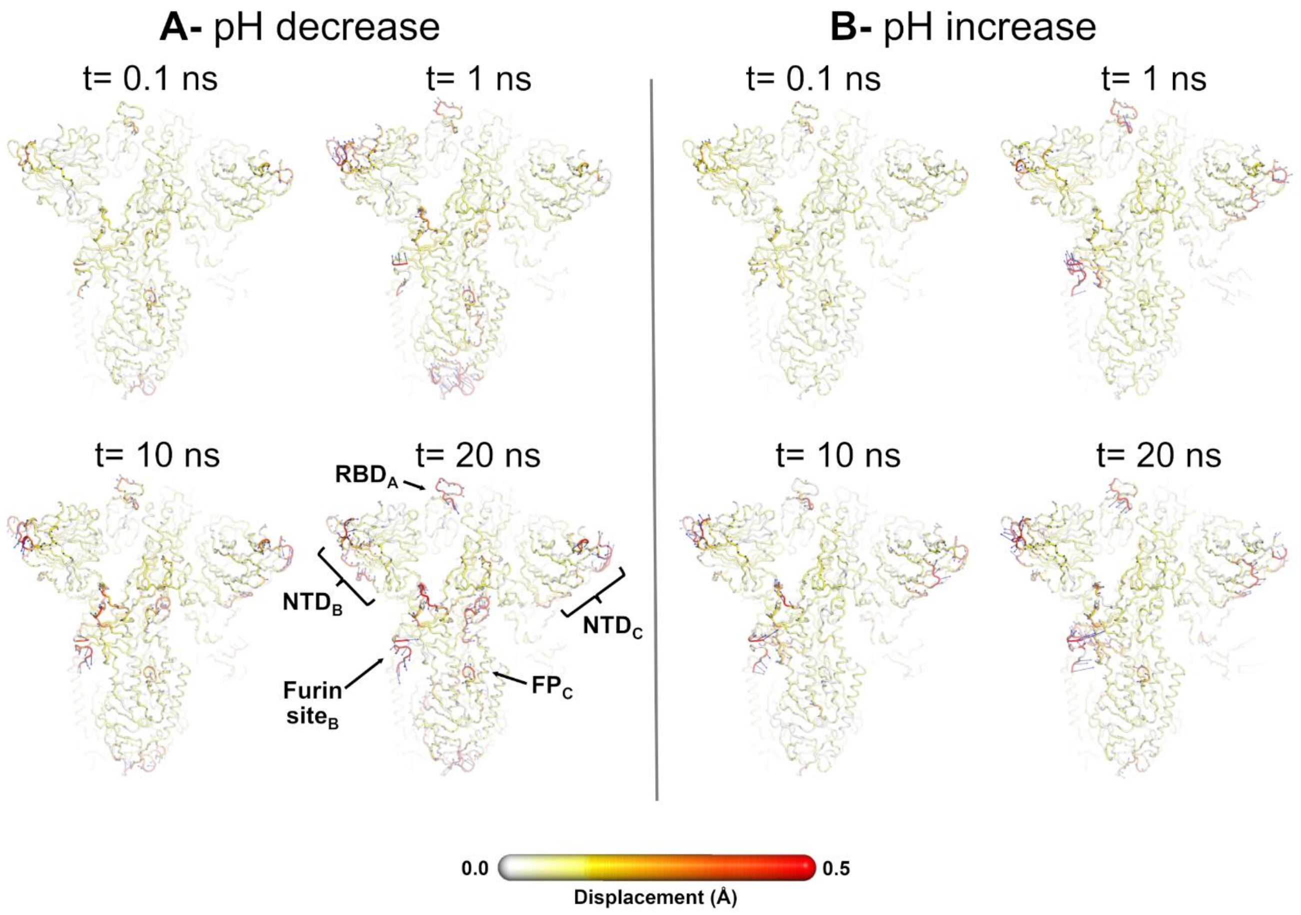


**Figure S20. Average displacement vectors from D-NEMD in response to a pH decrease (A) and increase (B) from the viewpoint of the RBD$_A$, NTD$_B$, NTD$_C$, and FP$_C$ for the ancestral spike.** This figure shows a full view of the protein, with average displacement vectors at *t* = 0.1, 1, 10, and 20 ns following a pH change. For details, see the caption of Figure S19. Please zoom in on the image for detailed visualisation.

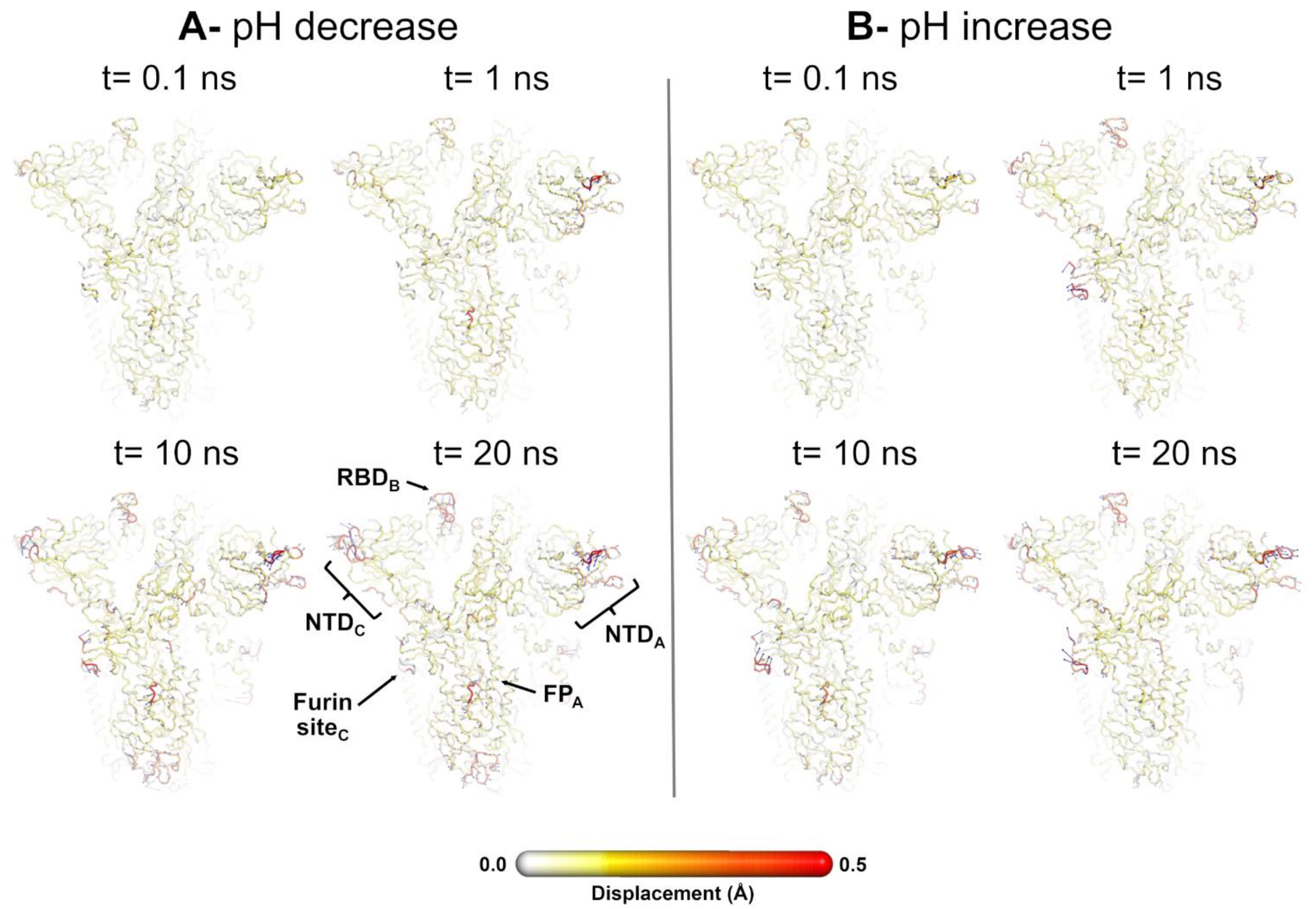


**Figure S21. Average displacement vectors from D-NEMD in response to a pH decrease (A) and increase (B) from the viewpoint of the $NTD_A$, $FP_A$, $RBD_B$, and $NTD_C$ for the ancestral spike.** This figure shows a full view of the protein, with average displacement vectors at *t* = 0.1, 1, 10, and 20 ns following a pH change. For details, see the caption of Figure S19. Please zoom in on the image for detailed visualisation.

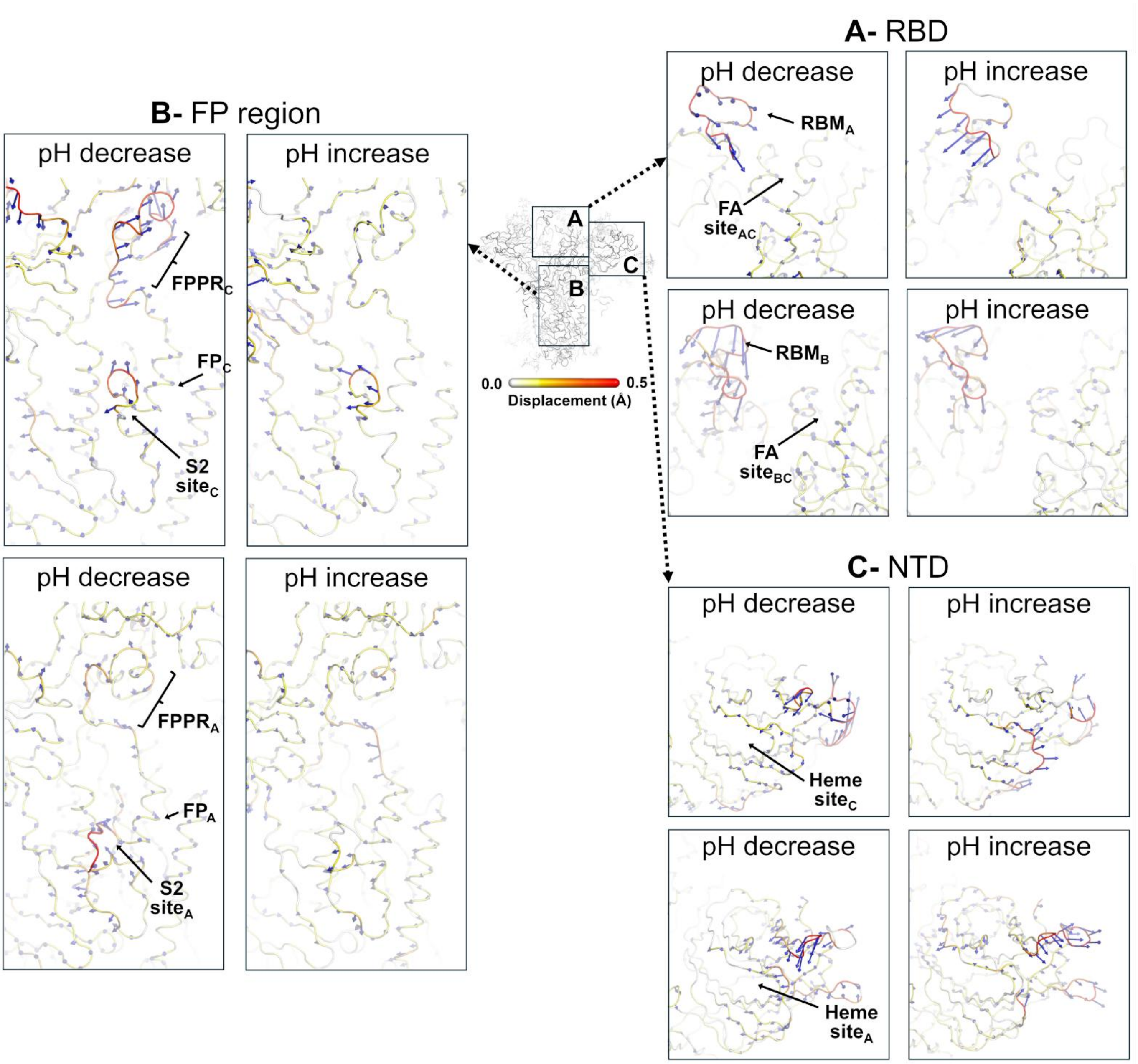


**Figure S22. Directionality of the pH-induced structural responses in the ancestral spike at *t* = 20 ns following the pH shift. (A)** Direction of the $RBM_A$ and $RBM_B$ motions in response to a pH decrease and increase. **(B)** Direction of the motions of the $FP_A$ and $FP_C$-surrounding regions in response to a pH decrease and increase. **(C)** Direction of the $NTD_A$ and $NTD_C$ motions in response to a pH decrease and increase. The blue arrows correspond to the average $C_\alpha$ displacement vectors (calculated between the equilibrium and nonequilibrium trajectories averaged across all 192 pairs of trajectories) after removing the intrinsic protein fluctuations via the "null perturbation" analysis. Vectors with a length ≥0.08 Å are displayed as blue arrows with a scale-up factor of 10. Average $C_\alpha$ displacements (i.e. the norm of the average $C_\alpha$ displacement vectors) are represented on a white-yellow-orange-red scale. Glycans were omitted from panels A, B and C to facilitate visualisation, but are present in all simulations. All regions labelled are subscripted with their chain ID (A, B or C). Please zoom in on the image for a detailed visualisation.

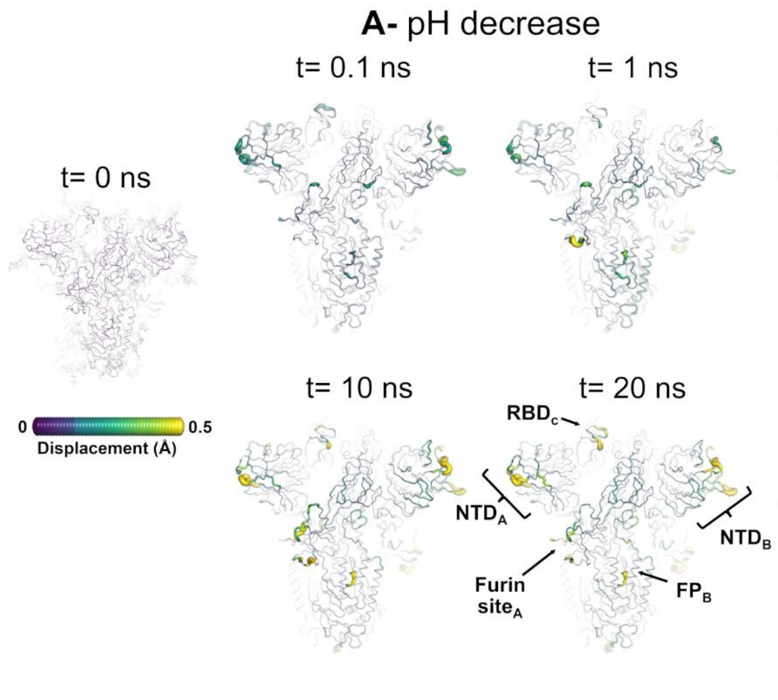


**Figure S23. Structural response of Delta to pH changes from the viewpoint of the $NTD_A$, $NTD_B$, $FP_B$ and $RBD_C$**. The average $C_\alpha$ displacements at $t$ = 0, 0.1, 1, 10, and 20 ns following a pH decrease **(A)** and increase **(B)** are shown. The displacements are mapped onto the starting structure for the equilibrium simulations of Delta at physiological pH. The magnitude of each residue's response was calculated using the Kubo-Onsager relation (34, 35) and corresponds to the norm of the average $C_\alpha$ displacement vector between every pair of equilibrium and nonequilibrium trajectories. The final displacements correspond to the average D-NEMD responses after removing the intrinsic protein fluctuations via the "null perturbation" analysis. Cartoon thickness and colours (scale shown on the left) indicate the extent of the average $C_\alpha$-positional displacements. Glycans are depicted as light grey sticks in the leftmost image but were omitted from the remaining panels (showing the responses at $t$=0.1, 1, 10 and 20 ns) to facilitate visualisation. Each RBD, NTD, furin site and FP are subscripted with their chain ID (A, B or C).

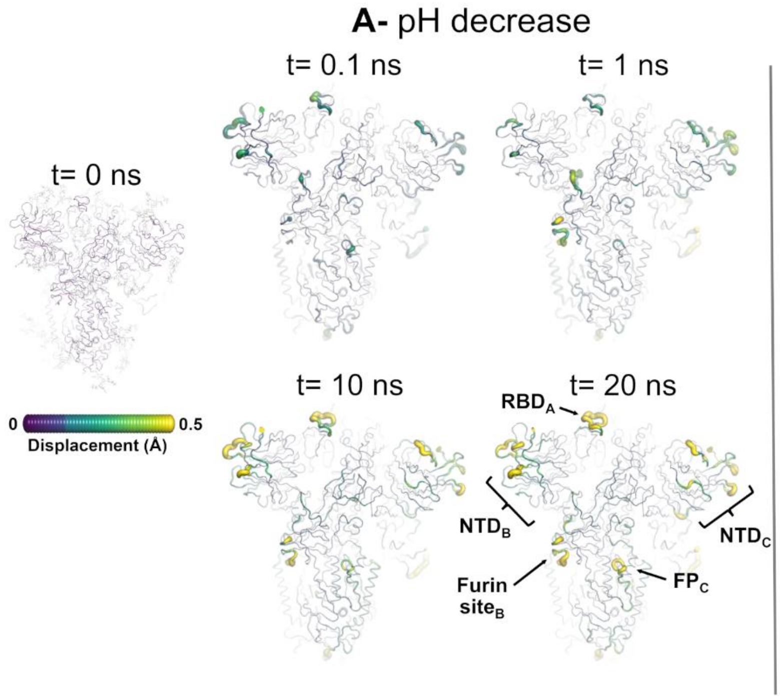


**Figure S24. Structural response of Delta to pH changes from the viewpoint of the RBD$_A$, NTD$_B$, NTD$_C$, and FP$_C$.** The average C$_\alpha$ displacements at *t* = 0, 0.1, 1, 10 and 20 ns following a pH decrease **(A)** and increase **(B)** are shown. For details, see the caption of Figure S23.

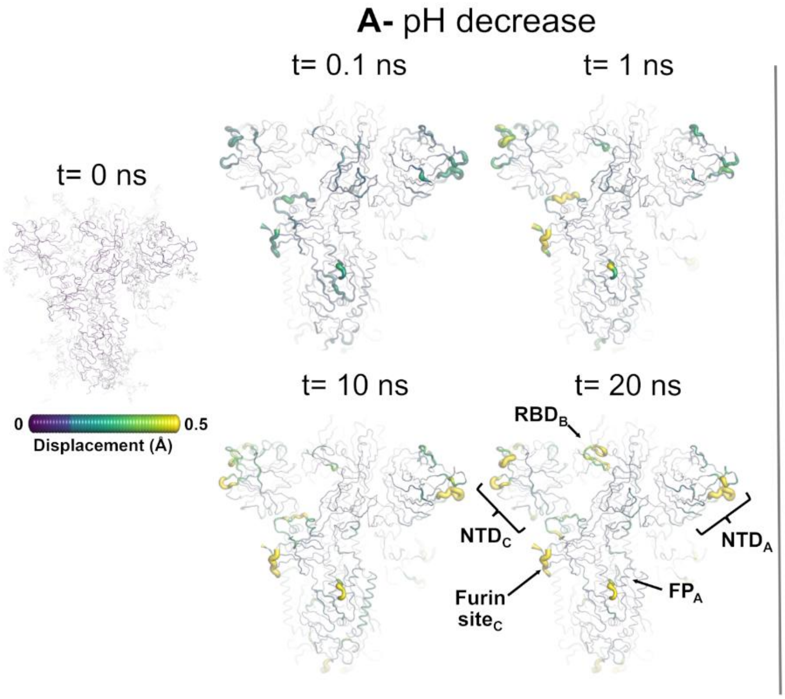


**Figure S25. Structural response of Delta to pH changes from the viewpoint of the $NTD_A$, $FP_A$, $RBD_B$, and $NTD_C$**. The average $C_\alpha$ displacements at $t$ = 0, 0.1, 1, 10 and 20 ns following a pH decrease **(A)** and increase **(B)** are shown. For details, see the caption of Figure S23.

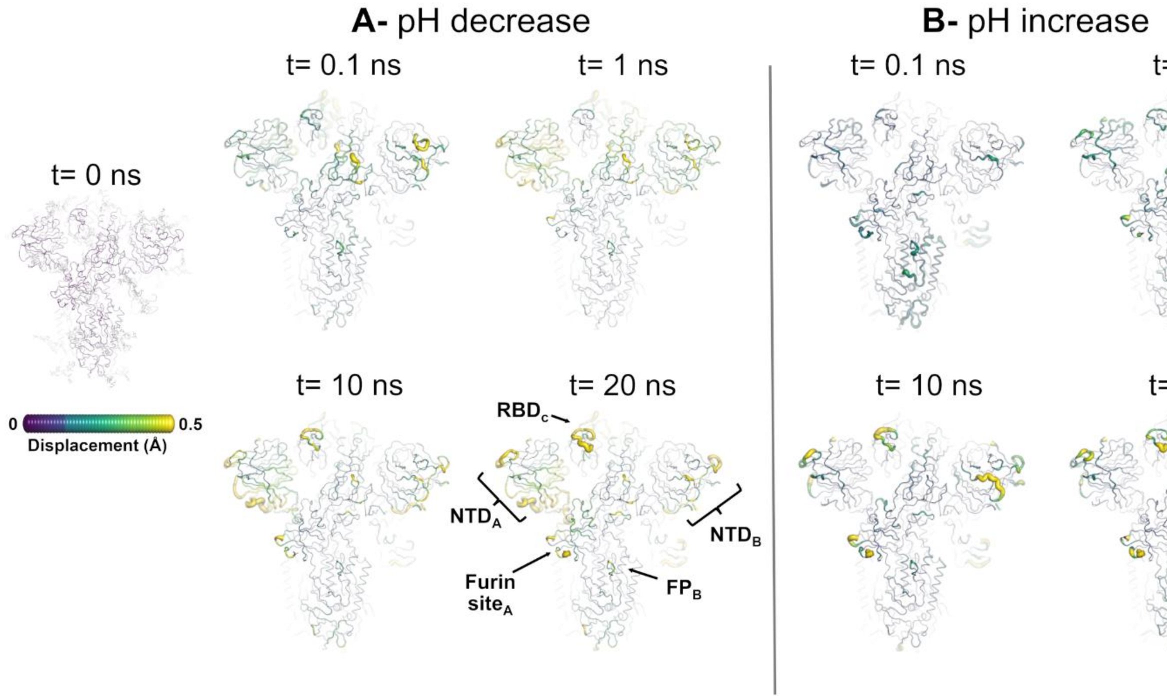


**Figure S26. Structural response of Omicron to pH changes from the viewpoint of the NTD$_A$, NTD$_B$, FP$_B$ and RBD$_C$.** The average C$_\alpha$ displacements at $t$ = 0, 0.1, 1, 10, and 20 ns following a pH decrease **(A)** and increase **(B)** are shown. The displacements are mapped onto the starting structure for the equilibrium simulations of Omicron at physiological pH. The magnitude of each residue's response was calculated using the Kubo-Onsager relation (34, 35) and corresponds to the norm of the average C$_\alpha$ displacement vector between every pair of equilibrium and nonequilibrium trajectories. The final displacements correspond to the average D-NEMD responses after removing the intrinsic protein fluctuations via the "null perturbation" analysis. Cartoon thickness and colours (scale shown on the left) indicate the extent of the average C$_\alpha$-positional displacements. Glycans are depicted as light grey sticks in the leftmost image but were omitted from the remaining panels (showing the responses at $t$=0.1, 1, 10 and 20 ns) to facilitate visualisation. Each RBD, NTD, furin site and FP are subscripted with their chain ID (A, B or C).

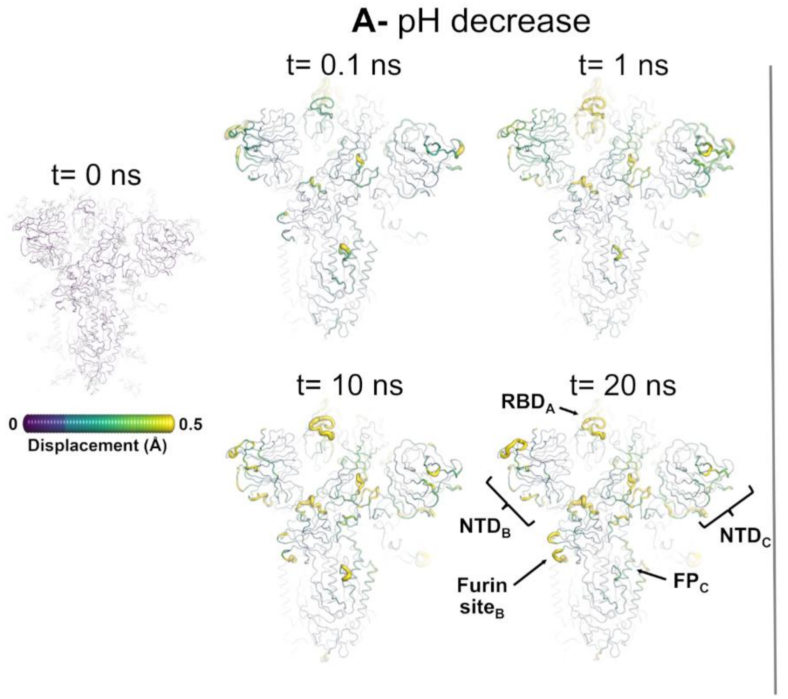


**Figure S27. Structural response of Omicron to pH changes from the viewpoint of the $RBD_A$, $NTD_B$, $NTD_C$, and $FP_C$**. The average $C_\alpha$ displacements at $t$ = 0, 0.1, 1, 10 and 20 ns following a pH decrease **(A)** and increase **(B)** are shown. For details, see the caption of Figure S26.

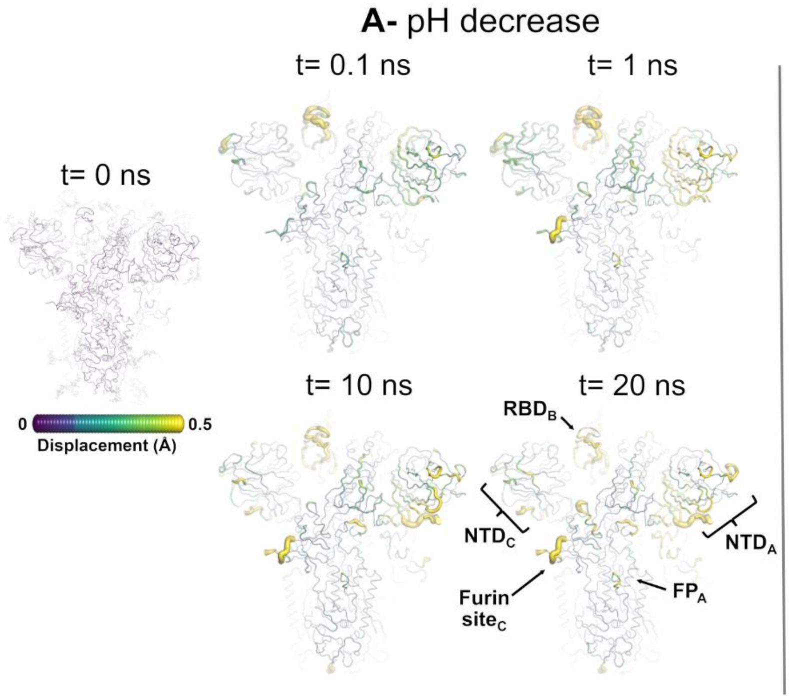


**Figure S28. Structural response of Omicron to pH changes from the viewpoint of the $NTD_A$, $FP_A$, $RBD_B$, and $NTD_C$**. The average $C_\alpha$ displacements at $t$ = 0, 0.1, 1, 10 and 20 ns following a pH decrease **(A)** and increase **(B)** are shown. For details, see the caption of Figure S26.

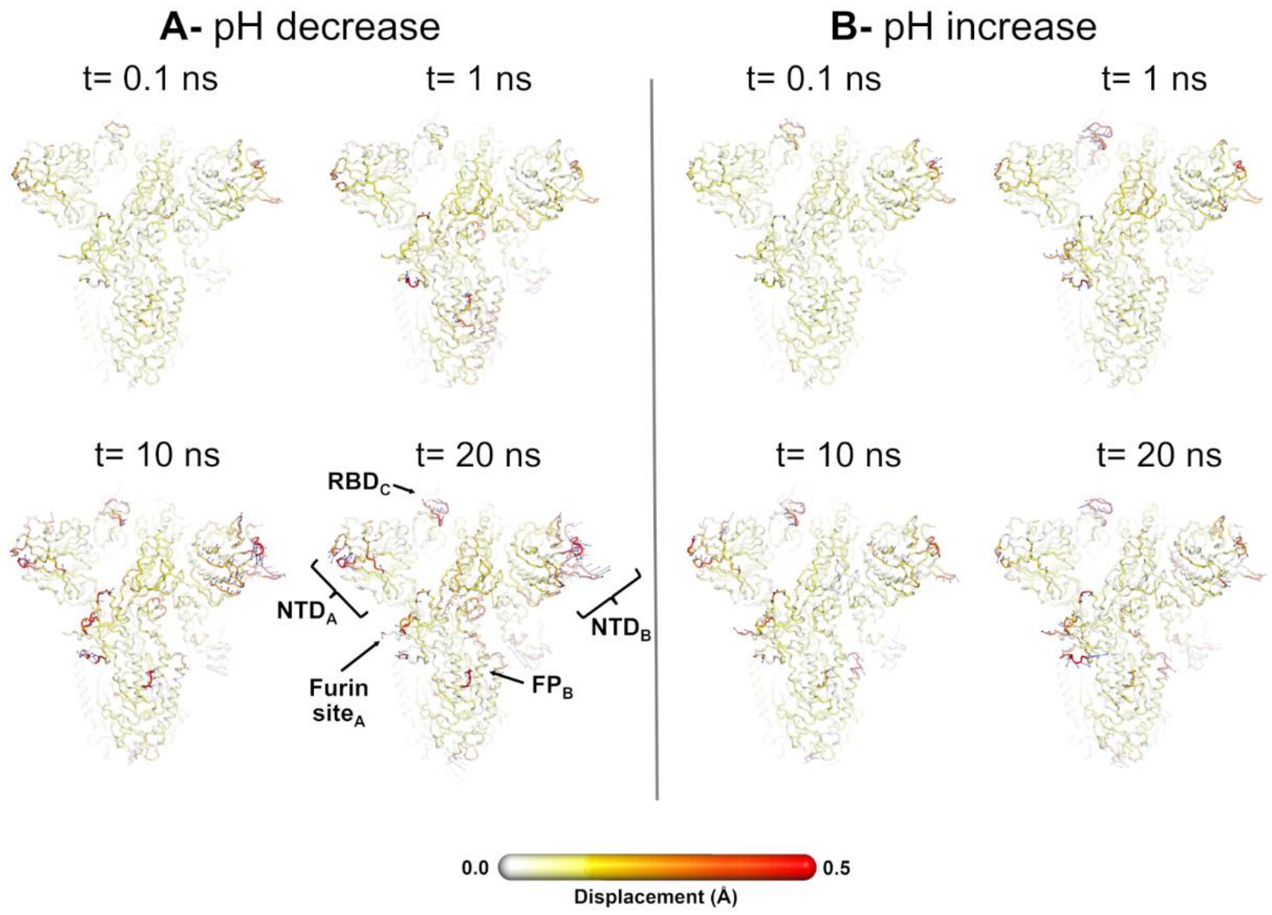


**Figure S29. Average displacement vectors from D-NEMD in response to a pH decrease (A) and increase (B) from the viewpoint of the NTD$_A$, NTD$_B$, FP$_B$, and RBD$_C$ for Delta.** This figure shows a full view of the protein, with average displacement vectors at $t$ = 0.1, 1, 10, and 20 ns following a pH change. The blue arrows correspond to the average $C_\alpha$ displacement vectors (calculated between the equilibrium and nonequilibrium trajectories averaged across all 192 pairs of trajectories) after removing the intrinsic protein fluctuations via the "null perturbation" analysis. Vectors with length ≥0.08 Å are displayed as blue arrows with a scale-up factor of 10. Average $C_\alpha$ displacements (i.e. the norm of the average $C_\alpha$ displacement vectors) are represented on a white-yellow-orange-red scale. Glycans were omitted from the figures to facilitate visualisation, but were present in the simulations. Each RBD, NTD, furin site and FP are subscripted with their chain ID (A, B or C). Please zoom in on the image for a detailed visualisation.

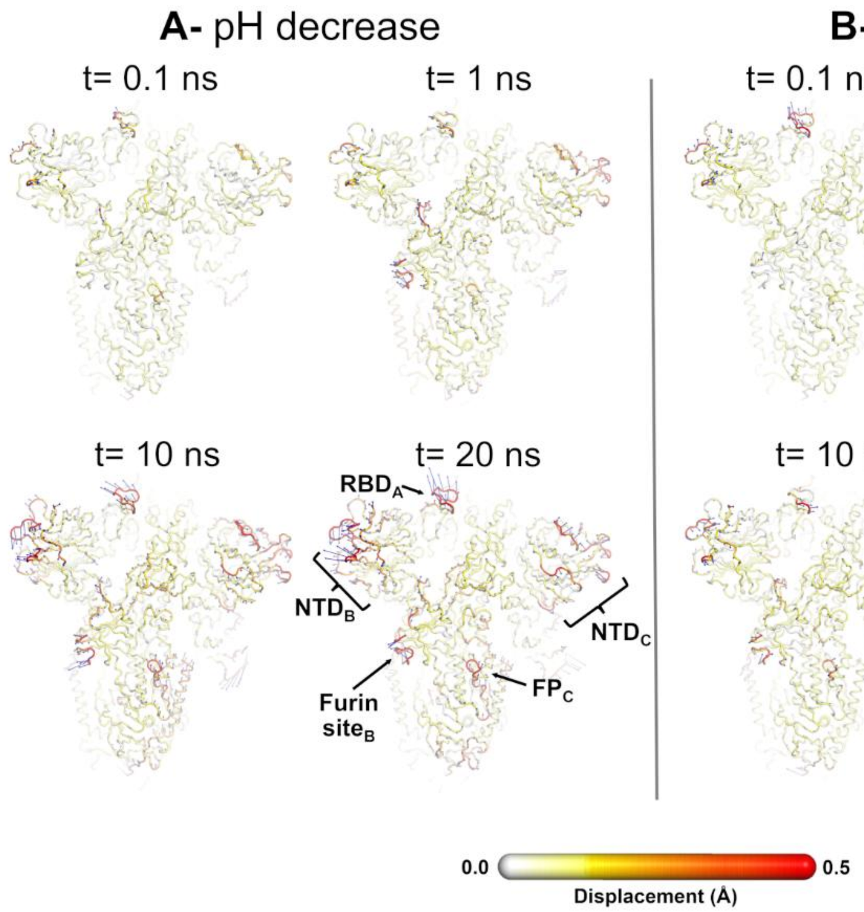


**Figure S30. Average displacement vectors from D-NEMD in response to a pH decrease (A) and increase (B) from the viewpoint of the RBD$_A$, NTD$_B$, NTD$_C$, and FP$_C$ for Delta.** This figure shows a full view of the protein, with average displacement vectors at $t$ = 0.1, 1, 10, and 20 ns following a pH change. For details, see the caption of Figure S29. Please zoom in on the image for detailed visualisation.

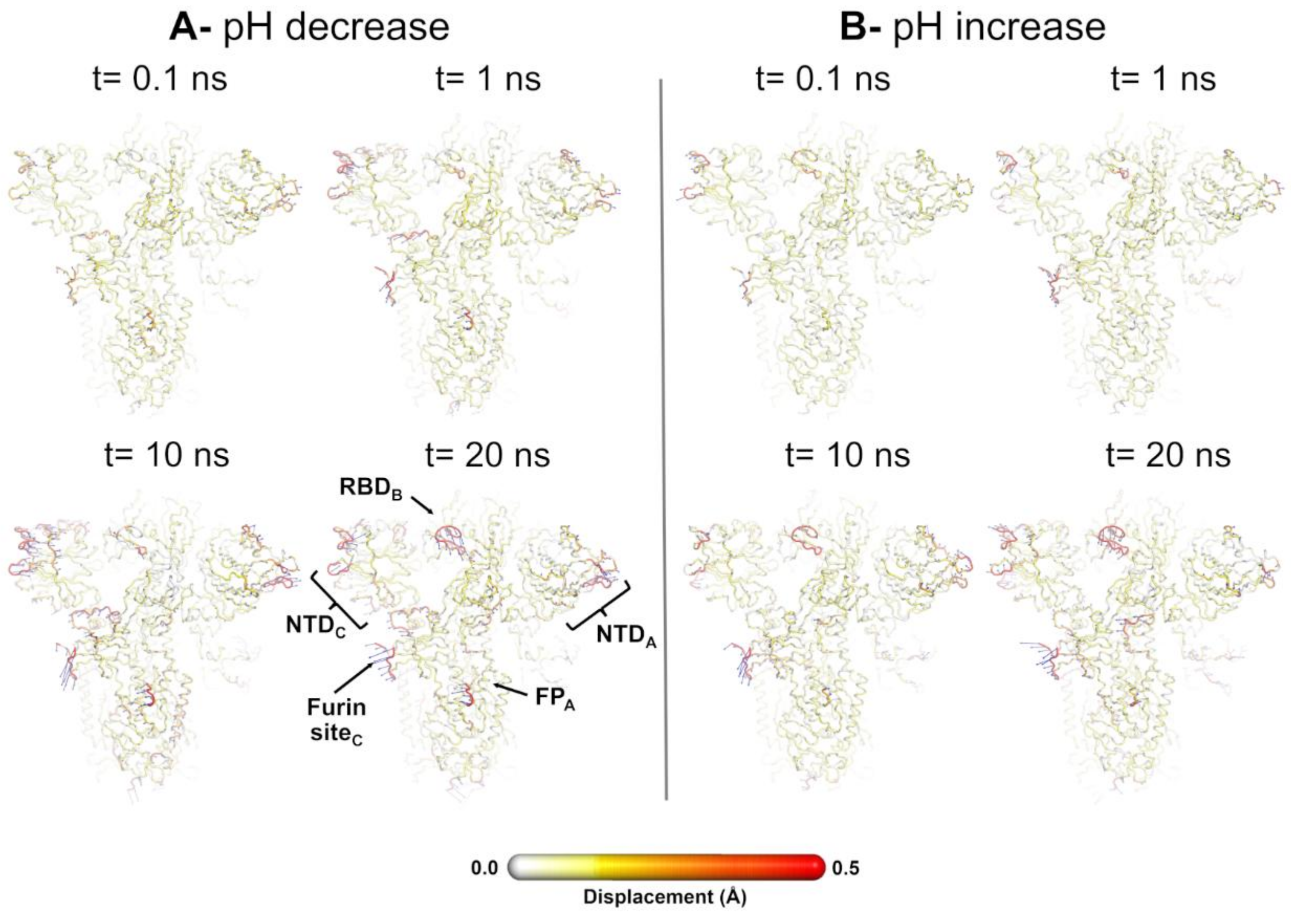


**Figure S31. Average displacement vectors from D-NEMD in response to a pH decrease (A) and increase (B) from the viewpoint of the $NTD_A$, $FP_A$, $RBD_B$, and $NTD_C$ for Delta.** This figure shows a full view of the protein, with average displacement vectors at $t$ = 0.1, 1, 10, and 20 ns following a pH change. For details, see the caption of Figure S29. Please zoom in on the image for detailed visualisation.

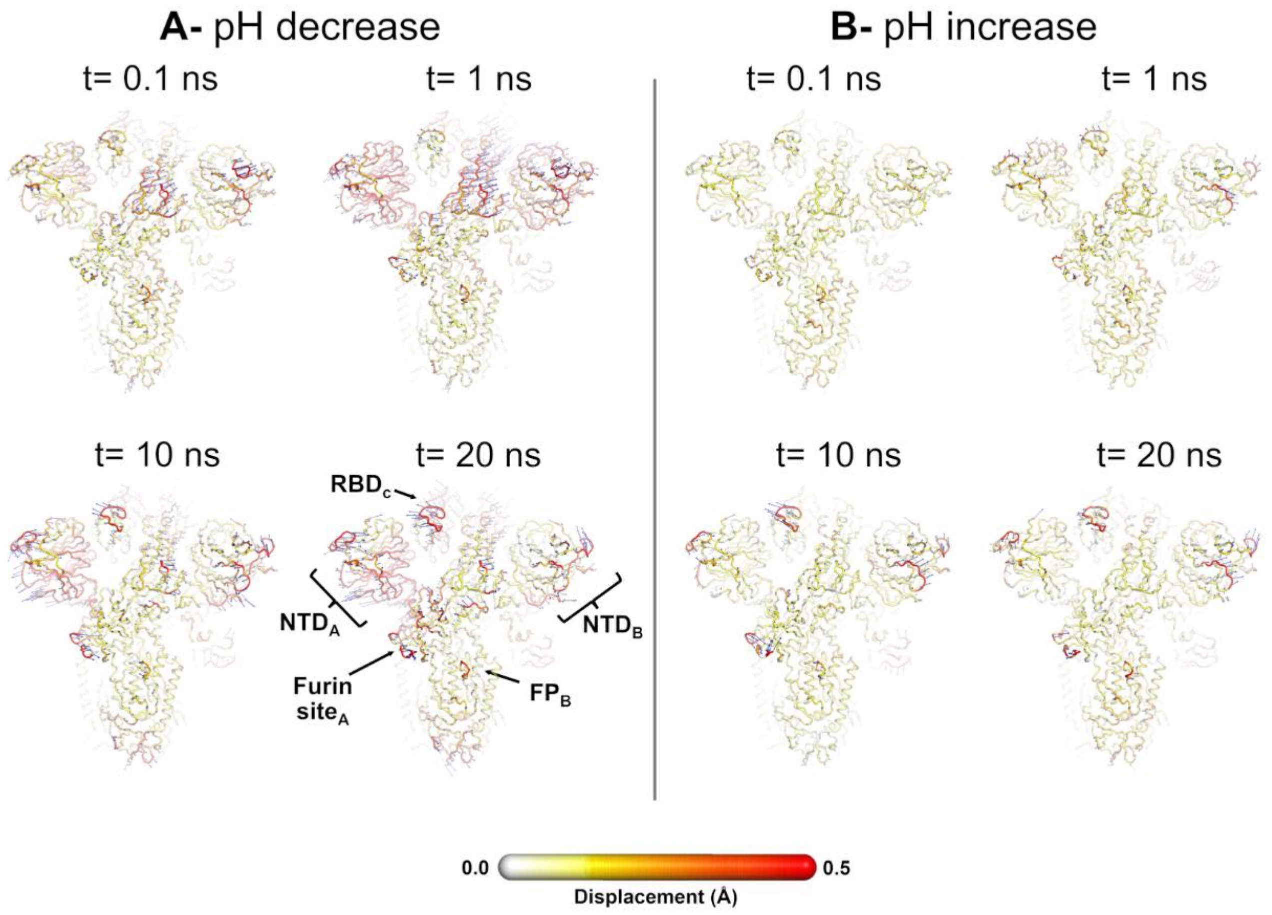


**Figure S32. Average displacement vectors from D-NEMD in response to a pH decrease (A) and increase (B) from the viewpoint of the $NTD_A$, $NTD_B$, $FP_B$, and $RBD_C$ for Omicron.** This figure shows a full view of the protein, with average displacement vectors at *t* = 0.1, 1, 10, and 20 ns following a pH change. The blue arrows correspond to the average $C_\alpha$ displacement vectors (calculated between the equilibrium and nonequilibrium trajectories averaged across all 192 pairs of trajectories) after removing the intrinsic protein fluctuations via the "null perturbation" analysis. Vectors with length ≥0.08 Å are displayed as blue arrows with a scale-up factor of 10. Average $C_\alpha$ displacements (i.e. the norm of the average $C_\alpha$ displacement vectors) are represented on a white-yellow-orange-red scale. Glycans were omitted from the figures to facilitate visualisation, but were present in the simulations. Each RBD, NTD, furin site and FP are subscripted with their chain ID (A, B or C). Please zoom in on the image for a detailed visualisation.

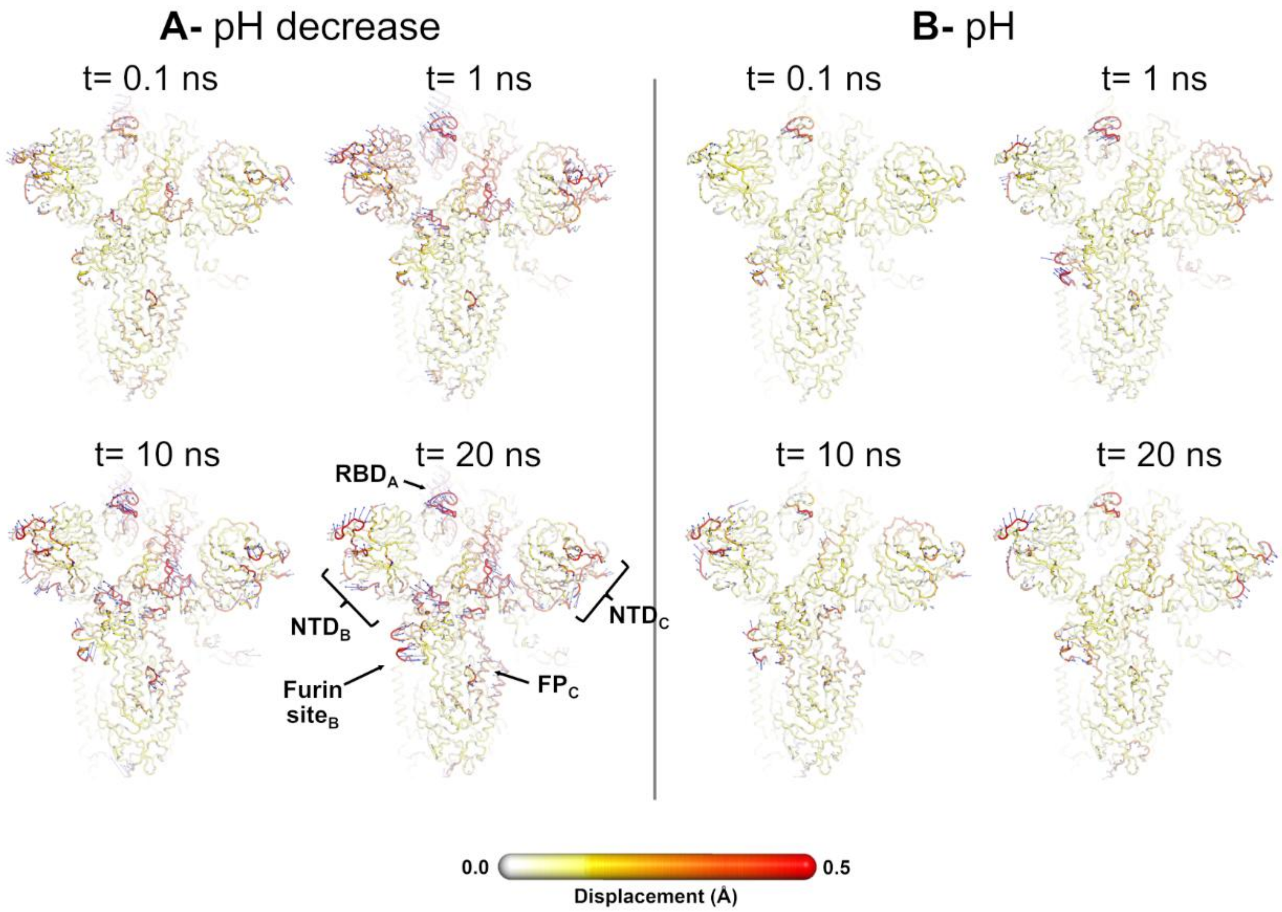


**Figure S33. Average displacement vectors from D-NEMD in response to a pH decrease (A) and increase (B) from the viewpoint of the $RBD_A$, $NTD_B$, $NTD_C$, and $FP_C$ for Omicron.** This figure shows a full view of the protein, with average displacement vectors at $t$ = 0.1, 1, 10, and 20 ns following a pH change. For details, see the caption of Figure S32. Please zoom in on the image for detailed visualisation.

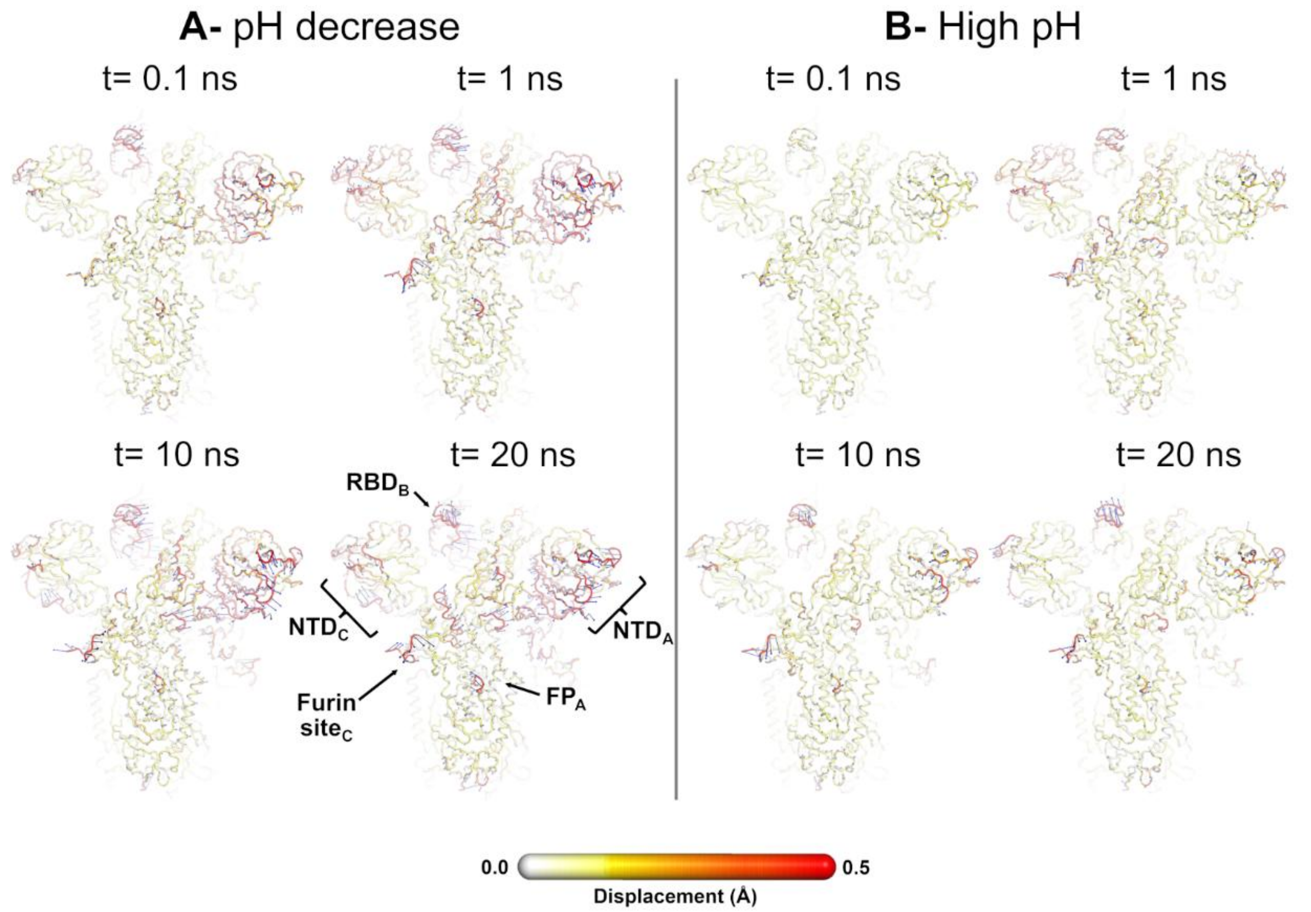


**Figure S34. Average displacement vectors from D-NEMD in response to a pH decrease (A) and increase (B) from the viewpoint of the $NTD_A$, $FP_A$, $RBD_B$, and $NTD_C$ for Omicron.** This figure shows a full view of the protein, with average displacement vectors at *t* = 0.1, 1, 10, and 20 ns following a pH change. For details, see the caption of Figure S32. Please zoom in on the image for detailed visualisation.

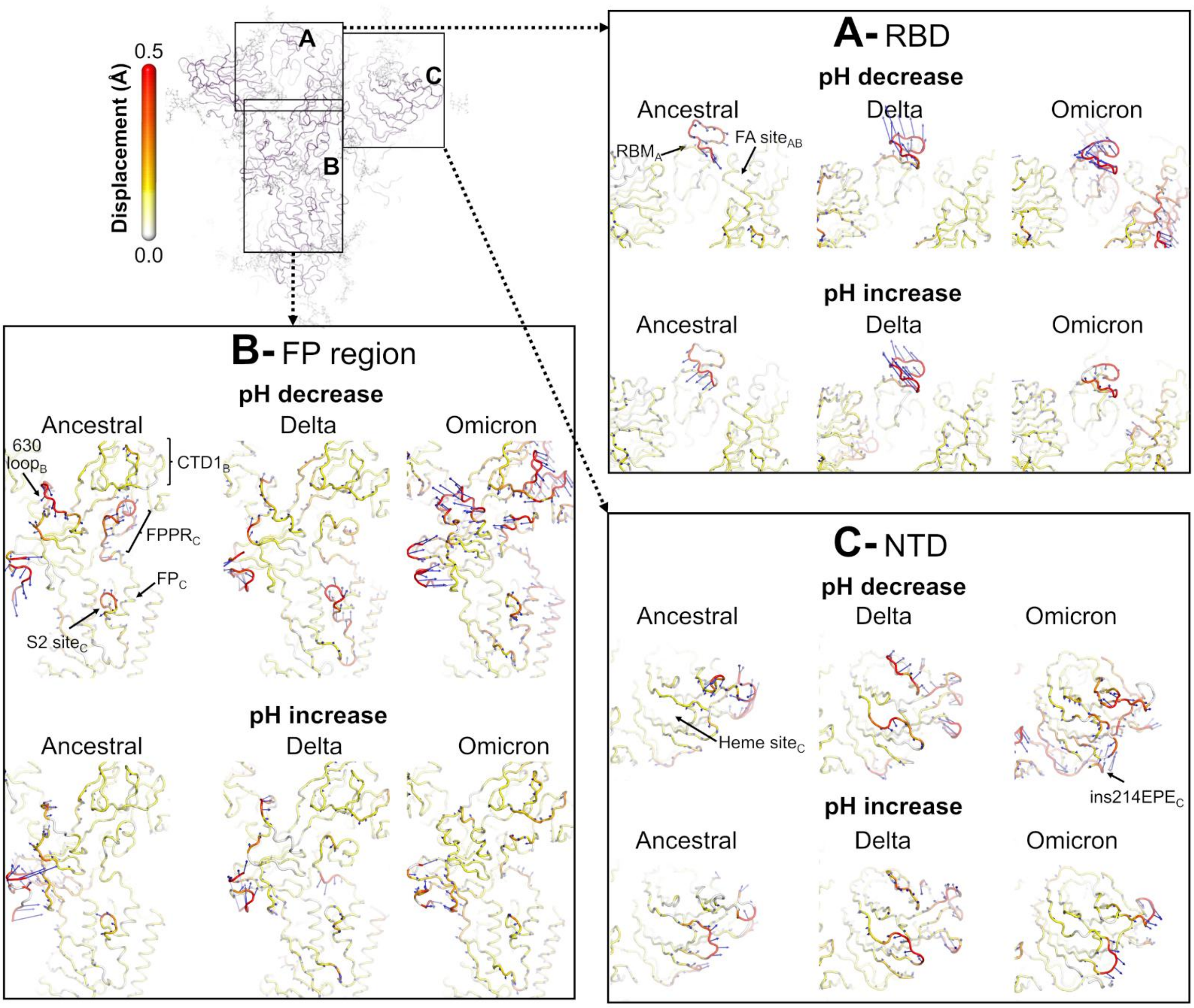


**Figure S35. Structural responses of ancestral, Delta, and Omicron spikes to pH changes from the viewpoint of $FP_C$, $RBM_A$ and $NTD_C$.** Comparative analysis of the pH-induced structural responses in ancestral, Delta, and Omicron at $t$ = 20 ns after the pH decrease/increase. Close-up view of the structural response of the **(A)** $RBD_A$; **(B)** $FP_C$-surrounding regions; and **(C)** $NTD_C$. The average $C_\alpha$ displacements at $t$ = 20 ns after the pH decrease/increase are shown, mapped onto the starting structure for the equilibrium simulations of the ancestral, Delta and Omicron proteins at physiological pH. The blue arrows correspond to the average $C_\alpha$ displacement vectors (calculated between the equilibrium and nonequilibrium trajectories across all 192 pairs of simulations) after removing the intrinsic protein fluctuations via the "null perturbation" analysis. Vectors with length ≥0.08 Å are displayed as blue arrows with a scale-up factor of 10. Average $C_\alpha$ displacement magnitudes are also represented on a white-yellow-orange-red scale. Glycans were omitted from the figures to facilitate visualisation, but were present in the simulations. Each RBD, NTD, furin site and FP are subscripted with their chain ID (A, B or C).

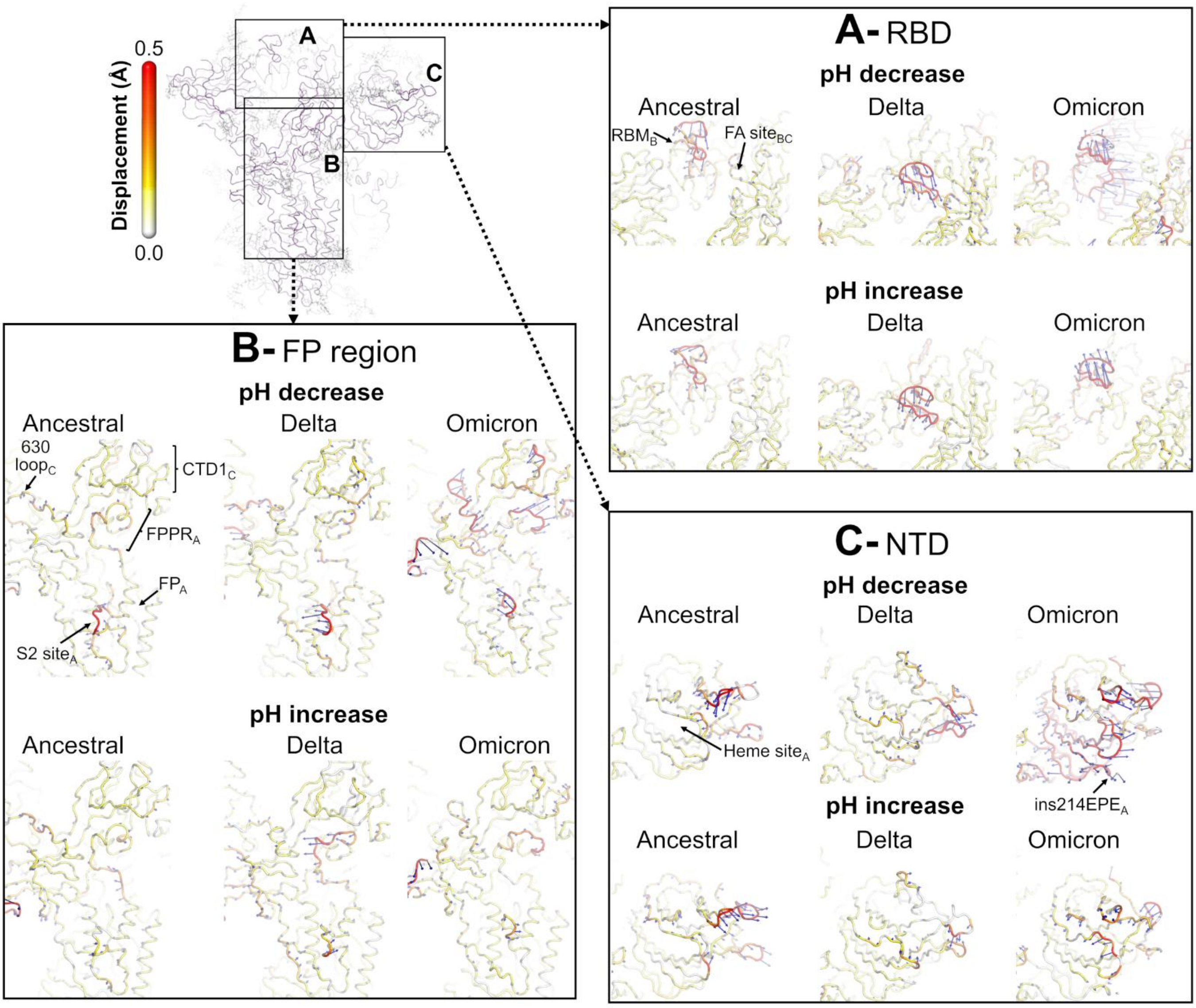


**Figure S36. Structural responses of ancestral, Delta, and Omicron spikes to pH changes from the viewpoint of $NTD_A$, $FP_A$, and $RBD_B$**. Comparative analysis of the pH-induced structural responses in ancestral, Delta, and Omicron at $t = 20$ ns after the pH decrease/increase. Close-up view of the structural response of the **(A)** $RBD_B$; **(B)** $FP_A$-surrounding regions; and **(C)** $NTD_A$. For details, see the caption of Figure S35.

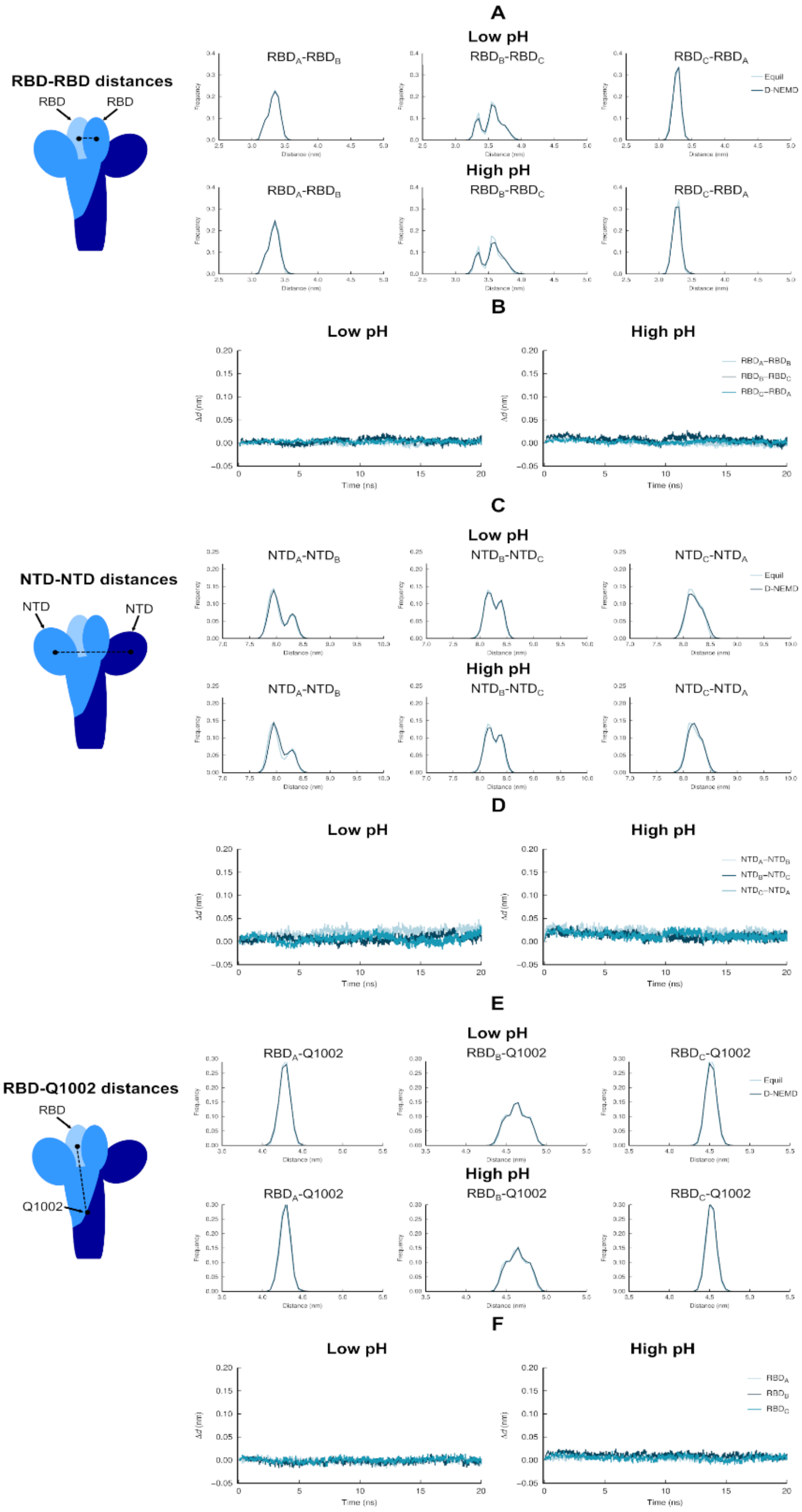


**Figure S37. RBD-RBD, NTD-NTD, and RBD-Q1002 distances in the equilibrium and nonequilibrium simulations of the ancestral spike. (A)** Distributions of distances between the centre of masses of $RBD_A$ and $RBD_B$, $RBD_B$ and $RBD_C$, and $RBD_C$ and $RBD_A$ in the equilibrium (at physiological pH) and nonequilibrium (at low/high pH) simulations. **(B)** Time evolution of the differences between the equilibrium and nonequilibrium trajectories for the RBD-RBD distances. Average Δ*d* (y-axis) is defined as the difference of the distance *d* between the centre of masses of $RBD_A$ and $RBD_B$, $RBD_B$ and $RBD_C$, and $RBD_C$ and $RBD_A$ in the nonequilibrium ($d_{neq}$) and equilibrium ($d_{eq}$) trajectories ($\Delta d = d_{eq} - d_{neq}$). **(C)** Distributions of distances between the centre of masses of the $NTD_A$ and $NTD_B$, $NTD_B$ and $NTD_C$, and $NTD_C$ and $NTD_A$ in the equilibrium (at physiological pH) and nonequilibrium (at low/high pH) simulations. **(D)** Time evolution of the differences between the equilibrium and nonequilibrium trajectories for the NTD-NTD distances. Average Δ*d* is the difference of the distance *d* between the centre of masses of the $NTD_A$ and $NTD_B$, $NTD_B$ and $NTD_C$, and $NTD_C$ and $NTD_A$ in the nonequilibrium ($d_{neq}$) and equilibrium ($d_{eq}$) trajectories ($\Delta d = d_{eq} - d_{neq}$). **(E)** Distributions of distances between the centre of mass of the $RBD_A$, $RBD_B$, and $RBD_C$ and the centre of mass of the three Q1002 residues (located in the central helices in the helical core of the protein) in the equilibrium (at physiological pH) and nonequilibrium (at low/high pH) simulations. **(F)** Time evolution of the differences between the equilibrium and nonequilibrium trajectories for the RBD-Q1002 distances. Average Δ*d* (y-axis) is defined as the difference of the distance *d* between the centre of mass of the $RBD_A$, $RBD_B$, and $RBD_C$ and the centre of mass of the three Q1002 in the nonequilibrium ($d_{neq}$) and equilibrium ($d_{eq}$) trajectories ($\Delta d = d_{eq} - d_{neq}$).The vertical blue lines in panels **B, D**, and **E** represent the standard error of the mean. Note that no significant differences are observed between the RBD-RBD, NTD-NTD and RBD-Q1002 distances in the equilibrium and nonequilibrium simulations. Please zoom in on the image for detailed visualisation.

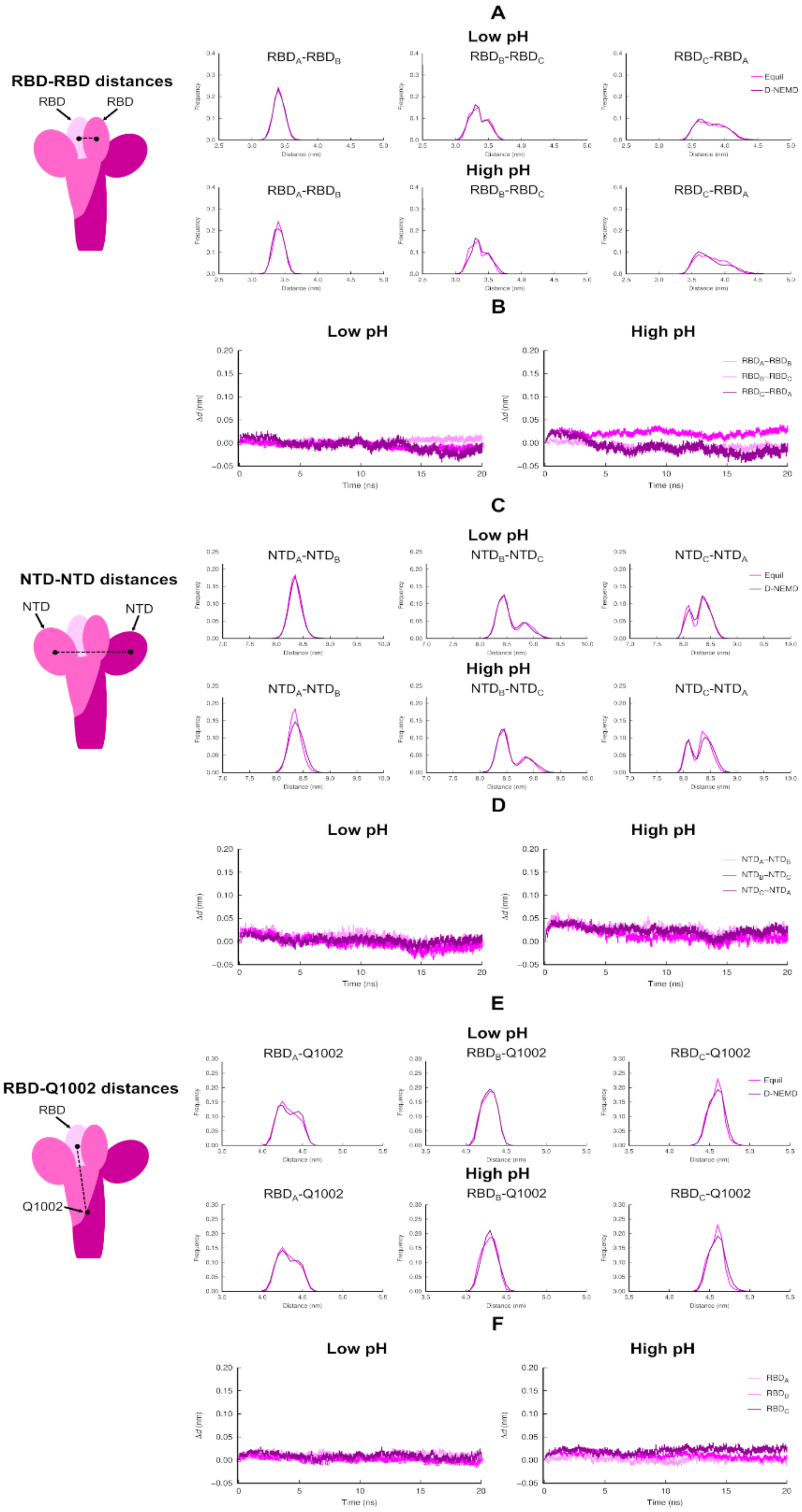


**Figure S38. RBD-RBD, NTD-NTD, and RBD-Q1002 distances in the equilibrium and nonequilibrium simulations of Delta.** **(A)** Distributions of distances between the centre of masses of $RBD_A$ and $RBD_B$, $RBD_B$ and $RBD_C$, and $RBD_C$ and $RBD_A$ in the equilibrium (at physiological pH) and nonequilibrium (at low/high pH) simulations. **(B)** Time evolution of the differences between the equilibrium and nonequilibrium trajectories for the RBD-RBD distances. Average Δ$d$ (y-axis) is defined as the difference of the distance $d$ between the centre of masses of $RBD_A$ and $RBD_B$, $RBD_B$ and $RBD_C$, and $RBD_C$ and $RBD_A$ in the nonequilibrium ($d_{neq}$) and equilibrium ($d_{eq}$) trajectories ($\Delta d = d_{eq} - d_{neq}$). **(C)** Distributions of distances between the centre of masses of the $NTD_A$ and $NTD_B$, $NTD_B$ and $NTD_C$, and $NTD_C$ and $NTD_A$ in the equilibrium (at physiological pH) and nonequilibrium (at low/high pH) simulations. **(D)** Time evolution of the differences between the equilibrium and nonequilibrium trajectories for the NTD-NTD distances. Average Δ$d$ is the difference of the distance $d$ between the centre of masses of the $NTD_A$ and $NTD_B$, $NTD_B$ and $NTD_C$, and $NTD_C$ and $NTD_A$ in the nonequilibrium ($d_{neq}$) and equilibrium ($d_{eq}$) trajectories ($\Delta d = d_{eq} - d_{neq}$). **(E)** Distributions of distances between the centre of mass of the $RBD_A$, $RBD_B$, and $RBD_C$ and the centre of mass of the three Q1002 residues (located in the central helices in the helical core of the protein) in the equilibrium (at physiological pH) and nonequilibrium (at low/high pH) simulations. **(F)** Time evolution of the differences between the equilibrium and nonequilibrium trajectories for the RBD-Q1002 distances. Average Δ$d$ (y-axis) is defined as the difference of the distance $d$ between the centre of mass of the $RBD_A$, $RBD_B$, and $RBD_C$ and the centre of mass of the three Q1002 in the nonequilibrium ($d_{neq}$) and equilibrium ($d_{eq}$) trajectories ($\Delta d = d_{eq} - d_{neq}$).The vertical blue lines in panels **B, D**, and **E** represent the standard error of the mean. Note that Δ$d$ (RBD-RBD) < 0 observed for Delta at high pH in panel **B** indicates that the distance between the RBDs is smaller in the nonequilibrium **c**ompared to the equilibrium simulations, thus suggesting changes leading towards a more closed state of the protein. Please zoom in on the image for detailed visualisation.

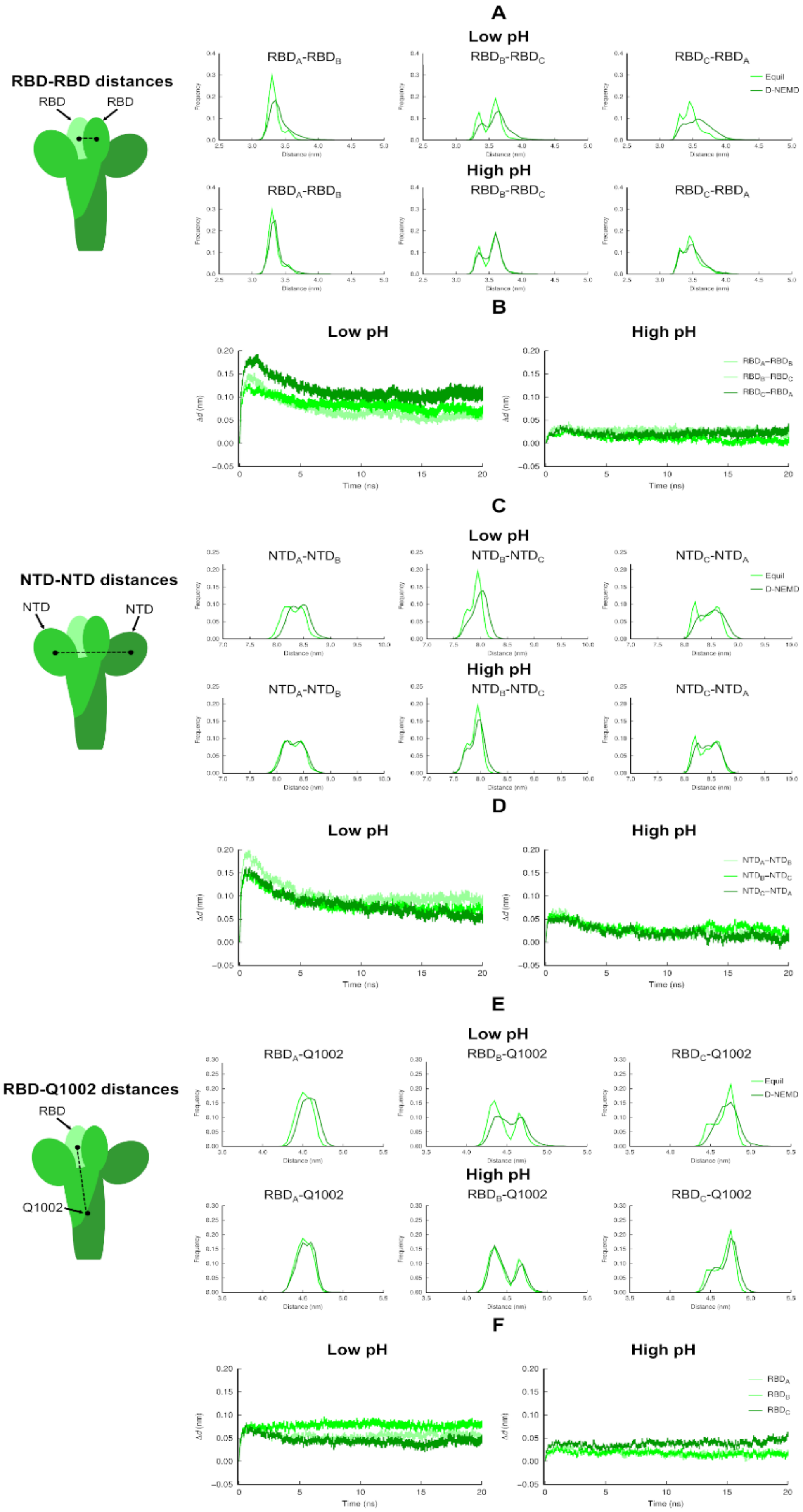


**Figure S39. RBD-RBD, NTD-NTD, and RBD-Q1002 distances in the equilibrium and nonequilibrium simulations of Omicron. (A)** Distributions of distances between the centre of masses of $RBD_A$ and $RBD_B$, $RBD_B$ and $RBD_C$, and $RBD_C$ and $RBD_A$ in the equilibrium (at physiological pH) and nonequilibrium (at low/high pH) simulations. **(B)** Time evolution of the differences between the equilibrium and nonequilibrium trajectories for the RBD-RBD distances. Average Δ*d* (y-axis) is defined as the difference of the distance *d* between the centre of masses of $RBD_A$ and $RBD_B$, $RBD_B$ and $RBD_C$, and $RBD_C$ and $RBD_A$ in the nonequilibrium ($d_{neq}$) and equilibrium ($d_{eq}$) trajectories ($\Delta d = d_{eq} - d_{neq}$). **(C)** Distributions of distances between the centre of masses of the $NTD_A$ and $NTD_B$, $NTD_B$ and $NTD_C$, and $NTD_C$ and $NTD_A$ in the equilibrium (at physiological pH) and nonequilibrium (at low/high pH) simulations. **(D)** Time evolution of the differences between the equilibrium and nonequilibrium trajectories for the NTD-NTD distances. Average Δ*d* is the difference of the distance *d* between the centre of masses of the $NTD_A$ and $NTD_B$, $NTD_B$ and $NTD_C$, and $NTD_C$ and $NTD_A$ in the nonequilibrium ($d_{neq}$) and equilibrium ($d_{eq}$) trajectories ($\Delta d = d_{eq} - d_{neq}$). **(E)** Distributions of distances between the centre of mass of the $RBD_A$, $RBD_B$, and $RBD_C$ and the centre of mass of the three Q1002 residues (located in the cental helices in the helical core of the protein) in the equilibrium (at physiological pH) and nonequilibrium (at low/high pH) simulations. **(F)** Time evolution of the differences between the equilibrium and nonequilibrium trajectories for the RBD-Q1002 distances. Average Δ*d* (y-axis) is defined as the difference of the distance *d* between the centre of mass of the $RBD_A$, $RBD_B$, and $RBD_C$ and the centre of mass of the three Q1002 in the nonequilibrium ($d_{neq}$) and equilibrium ($d_{eq}$) trajectories ($\Delta d = d_{eq} - d_{neq}$).The vertical blue lines in panels **B, D**, and **E** represent the standard error of the mean. Note that $\Delta d > 0$ observed for Omicron at low pH in panels **B** and **D** means that the distance between the RBD-RBD/NTD-NTD is greater in the nonequilibrium compared to the equilibrium simulations, thus indicating pH-induced conformational responses. This increase in distance in Omicron at low pH indicates that the spike trimer undergoes conformational changes consistent with a transition from a closed to an up RBD state. Please zoom in on the image for detailed visualisation.

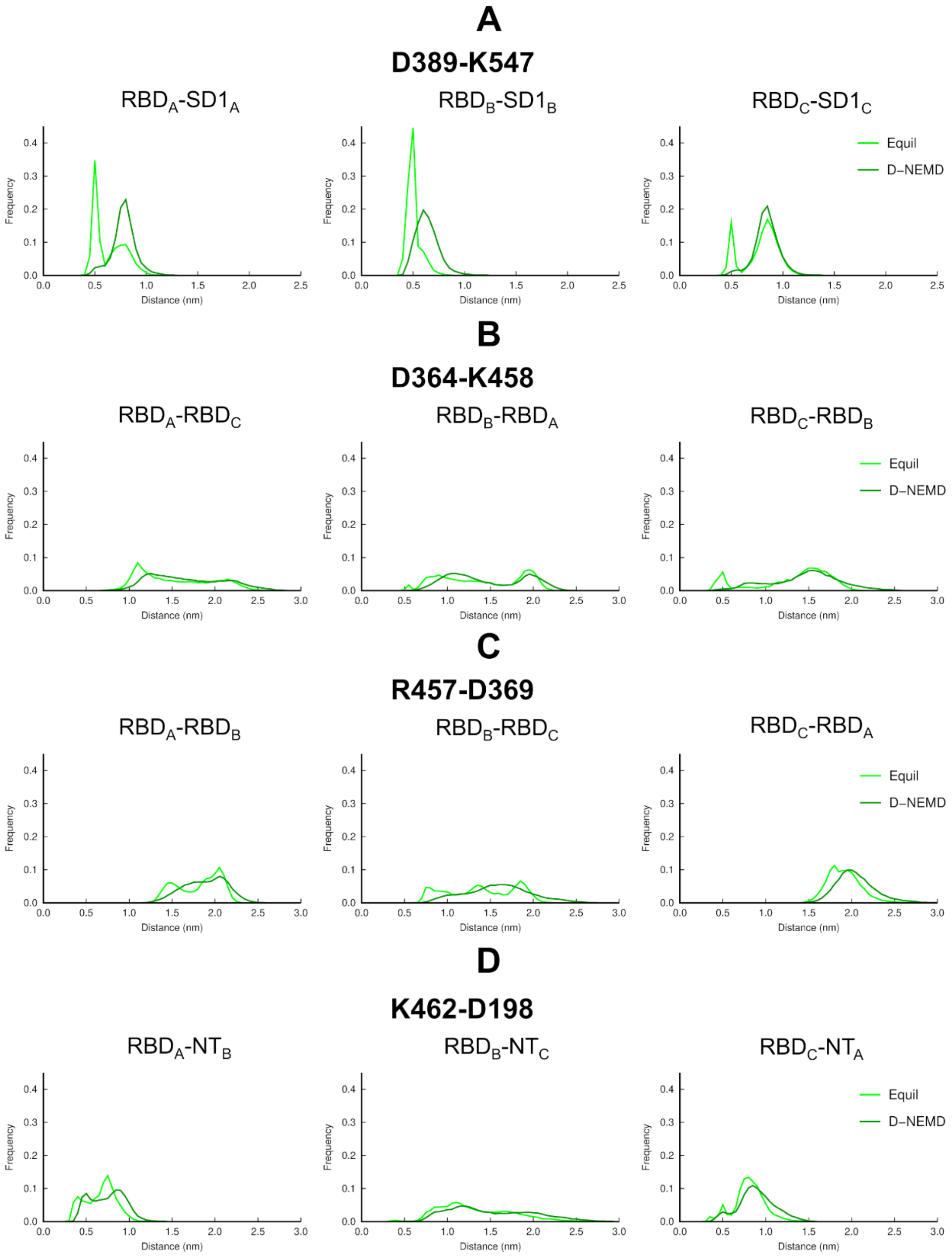


**Figure S40. Distribution of distances between residues forming salt bridges between the RBD and other regions of Omicron in equilibrium simulations at physiological pH and nonequilibrium simulations under acidic conditions**. Panels show distances between the centres of mass of the side chains of: **(A)** D389 (RBD) and R457 (SD1), **(B)** D364 and K458 (RBD), **(C)** R457 and D364 (RBD), and **(D)** K462 (RBD) and D198 (NTD). Previous weighted ensemble simulations have identified the disruption of salt bridges involving D364, R457, K458, and K462 as part of the RBD opening pathway (3). The observed increase in the distance between salt-bridge-forming residues under acidic conditions reflects conformational changes in Omicron, associated with a higher population of conformations in which the salt bridges are disrupted. Please zoom in for detailed visualisation.
